# Evolutionary Trajectory of Pattern Recognition Receptors in Plants

**DOI:** 10.1101/2023.07.04.547604

**Authors:** Bruno Pok Man Ngou, Michele Wyler, Marc W Schmid, Yasuhiro Kadota, Ken Shirasu

## Abstract

Plants perceive pathogen-associated molecular patterns (PAMPs) via pattern recognition receptors (PRRs) to activate PRR-triggered immunity (PTI). Despite extensive research on PTI in model plant species, the evolutionary trajectory and emergence of PRRs remain elusive. Here we conducted a comparative genomic analysis of cell-surface receptors and downstream signalling components among 350 plant species. Our findings reveal that cell-surface receptors comprise two major classes, receptor-like proteins (RLPs) and receptor-like kinases (RLKs), with RLP being more ancient whereas RLK families have undergone significant expansion. We also demonstrate that multiple downstream signalling components have an ancient origin within the plant lineage. To shed light on the immune-specificity of PRRs, we traced the evolutionary origin of immune-specific leucine-rich repeat-RLPs (LRR-RLPs) in plants. Surprisingly, we discovered that the last four LRR motifs crucial for co-receptor interaction in LRR-RLPs are closely related to those of the LRR-RLK subgroup Xb, which primarily governs growth and development. Functional characterisation further reveals that LRR-RLPs initiate immune responses through their juxtamembrane and transmembrane regions, while LRR-RLK-Xb members regulate development through their cytosolic kinase domains. Our data suggest modular evolution of cell-surface receptors in which immunity- and development-specific cell-surface receptors share a common origin. After diversification, their ectodomains, juxtamembrane, transmembrane, and cytosolic regions have either diversified or stabilised to recognize ligands that activate different downstream responses. We propose that cell-surface receptors and downstream signalling components are ancient, and likely predate the emergence of land plants, subsequently evolving to exhibit greater complexity and specificity within the land plant lineage.

## Introduction

Recognition of pathogen-associated molecular patterns (PAMPs) via pattern-recognition receptors (PRRs) is a fundamental mechanism which allows organisms to detect the presence of pathogens^1,2^. PAMP/PRR-triggered immunity (PTI) is conserved in animals, plants, fungi, and other eukaryotes^3–5^. Plant PRRs play a crucial role in perceiving various PAMPs from pathogens, including peptides, small proteins, lipids, and polysaccharides^2,6^. These receptors can be categorised as either receptor-like proteins (RLPs), which possess a transmembrane domain, or receptor-like kinases (RLKs), which have both transmembrane and kinase domains. The number of RLP and RLK gene families varies among plant species, and is believed to expand over evolutionary time in response to pathogen pressure^7,8^. To date, around seventy plant PRRs with known elicitors have been characterised, primarily in angiosperms^6^, while those in other clades of Viridiplantae remain poorly understood.

Upon ligand or elicitor perception, PRRs undergo dimerization or form heteromeric complexes with other cell-surface receptors. This spatial arrangement brings the cytoplasmic kinase domains of PRRs in close proximity, initiating a cascade of auto- and trans-phosphorylation events^9^. The activated receptor complex subsequently phosphorylates members of the cytoplasmic receptor-like kinases subgroup VII (RLCK-VII)^10^, which, in turn, phosphorylate various cytoplasmic kinases, such as the mitogen-activated protein kinase kinase kinases (MAPKKKs), and plasma membrane-associated proteins, such as calcium channels and NADPH oxidases^9^. The phosphorylation of these proteins collectively triggers transcriptional reprogramming and physiological changes, such as cytoplasmic calcium influx and the accumulation of reactive oxygen species (ROS)^11^. These physiological responses effectively hinder pathogen proliferation during infection (Extended Figure 1). The PTI-signalling pathway has been extensively characterised in model plant species such as Arabidopsis, rice, and tomato. However, the origins of PRRs and PTI-signalling components remain unclear, and the evolutionary relationships between receptors and downstream signalling components have yet to be determined.

While many cell-surface receptors are involved in pathogen recognition, many with similar domain architectures are also engaged in other biological processes (Extended Figure 2). Some of these receptors recognise endogenous molecules, such as phytohormones and phytocytokines, to regulate developmental and reproductive processes^12,13^. Given the striking resemblance in their domain architecture, it is reasonable to infer that immunity-and developmental-related cell-surface receptors share a common origin. However, the evolutionary trajectory that led to their divergence and specialisation in distinct biological processes remains poorly understood.

## Results

### Diversity and immune relevance of cell-surface receptors in Viridiplantae

The PRRs known to participate in immunity include LRR-RLKs, LRR-RLPs, G-lectin-RLKs, Wall-associated kinase (WAK)-RLKs, WAK-RLPs, Domain of Unknown Function 26 (Duf26)-RLKs (or cysteine-rich kinases; CRKs), L-lectin-RLK, Lysin motif (LysM)-RLKs, and Malectin-RLKs^6^. LRR-RLKs are further classified into 20 subgroups based on their kinase domains, with subgroup XII specifically implicated in PAMP recognition^14^. PRRs with LRR-, G-lectin-, WAK-, and LysM-ectodomains recognise PAMPs, while others perceive self-molecules or unidentified ligands. These ligands encompass a wide range of substances such as small molecules, peptides, full-length proteins, lipids, and polysaccharides/glycans (Extended Figure 2). Recognition of the diverse array of ligands is likely to be accomplished by variable structures and combinations of different ectodomains (Main figure 1a-g).

**Main figure 1.**
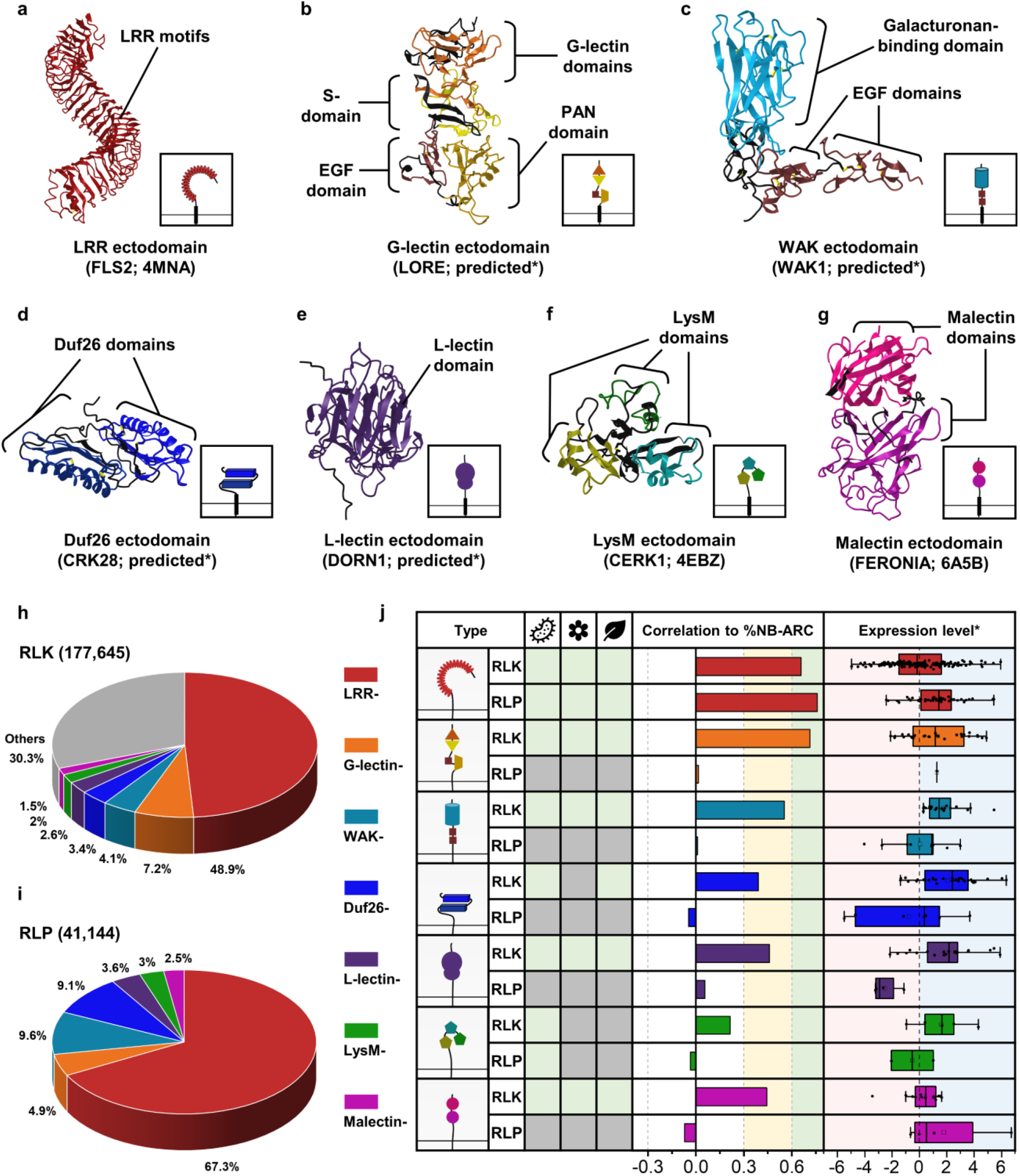
The distribution of cell-surface receptors in plants. Ectodomain structure of a) an LRR receptor, b) a G-lectin receptor, c) an L-lectin receptor, d) a LysM receptor, e) a Malectin receptor, f) a WAK receptor, and g) a Duf26 receptor. Structures of FLS2, CERK1 and FERONIA were published^17–19^. Structures of LORE, DORN1, WAK1 and CRK28 were predicted by Alphafold2*,^20^. Ectodomains are visualized in iCn3D^21^. **H)** Ectodomain distribution of RLKs in plants. Each fraction represents the percentage (%) of ectodomain out of all the RLKs from 350 species (177,645). **I)** Ectodomain distribution of RLP in plants. Each fraction represents the percentage (%) of ectodomains out of all the RLPs with those seven ectodomains (41,144). **J)** Table of RLKs and RLPs with LRR (red), G-lectin (orange), WAK (turquoise), Duf26 (blue), L-lectin (purple), LysM (green), and Malectin (magenta) ectodomains. Characterised receptors involved in microbial interaction (bacteria icon), reproduction (flower icon), and development (leaf icon) are indicated with light green boxes. Grey boxes indicate that the receptor class has not been reported to be involved in that biological process. For details, refer to Extended Figure 2. Correlations between different classes of cell-surface receptors and NB-ARC in 300 angiosperms are indicated with bars. Strong positive correlations are indicated by extension to the light green area (Pearson’s r >0.6) and medium positive correlations are within the yellow area (Pearson’s r between 0.3-0.6). Expression level* refers to the expression of each class of cell-surface receptors during NLR-triggered immunity (NTI) in *Arabidopsis thaliana*. Light blue area represents increased expression and light pink area represents decreased expression during NTI. X-axis values represent log_2_ (fold change during ETI relative to untreated samples). Boxplot elements: center line, median; bounds of box, 25th and 75th percentiles; whiskers, 1.5 × IQR from 25th and 75th percentiles. RNA-seq data analysed here were reported previously, where NTI was activated by estradiol-induced expression of AvrRps4 in *A. thaliana* for 4 hours^16^. For the expression of each class of cell-surface receptors during PTI in *A. thaliana*, refer to supplementary figure 1h-n.

To investigate the phylogenetic relationships of the cell-surface receptor-gene family in Viridiplantae, we extracted RLKs containing both a transmembrane (TM) and a kinase domain (KD) from 350 publicly available genomes as previously reported^8^. In total, we identified approximately 177,645 RLKs meeting this criterion. We subsequently pinpointed RLKs with LRR-, G-lectin-, WAK-, Duf26-, L-lectin-, LysM-, and Malectin-ectodomains. Among these, the most prominent RLK family are the LRR-RLKs, which account for 48.9% of the identified RLKs, followed by G-lectin-RLKs (7.2%), WAK-RLKs (4.1%), Duf26-RLKs (3.4%), L-lectin-RLKs (2.6%), LysM-RLKs (2%) and Malectin-RLKs (1.5%) (Main figure 1h). Collectively, these seven families constitute nearly 70% of all identified RLKs. Previous reports have suggested a positive correlation between the gene family sizes of cell-surface immune receptors (LRR-RLK subgroup XII and LRR-RLPs) and intracellular immune receptors (NLRs, NB-ARC family) across the angiosperms^8^. To test this observation, we examined the correlation between the percentage (%; number of identified genes / numbers of searched genes × 100) of the seven RLK families and the NB-ARC family in each genome. Notably, both %LRR-RLK and %G-lectin-RLK exhibit robust positive correlations (Pearson’s r > 0.6) with %NB-ARC (Main figure 1j). Additionally, %WAK-RLK, %Duf26-RLK, %L-lectin-RLKs and %Malectin-RLKs demonstrate moderate positive correlations (0.6 > Pearson’s r > 0.3) with %NB-ARC (Main figure 1j). Thus, it is likely that members of these RLK families play significant roles in immunity.

Next, we extracted protein sequences with TM and LRR-/G-lectin-/WAK-/Duf26-/L-lectin-/LysM/Malectin-ectodomains but lacking KDs (RLPs). In total, we identified 41,144 RLPs. Again, the largest RLP family comprises LRR-RLKs (67.3%), followed by WAK-RLPs (9.6%), Duf26-RLPs (9.1%), G-lectin-RLPs (4.9%), L-lectin-RLPs (3.6%), LysM-RLPs (3%) and Malectin-RLPs (2.5%) (Main figure 1i). The RLK repertoire surpasses that of RLPs, indicating either the expansion of RLK families, presumably by gene duplication, or contraction of RLP families throughout the plant lineage. We therefore conducted correlation analyses to assess the relationship between the percentages of the seven RLP families and NB-ARC family in each genome. Surprisingly, with the exception of LRR-RLPs (Pearson’s r = 0.759), the sizes of the remaining six RLP families exhibit negligible correlations with %NB-ARC (Pearson’s r < 0.1) (Main figure 1j), suggesting that, apart from LRR-RLPs, the majority of members in these six RLP families are unlikely to be involved in immunity. This observation aligns with the absence of reports regarding the role of G-lectin-RLPs, WAK-RLPs, Duf26-RLPs, L-lectin-RLPs, and Malectin-RLPs in immunity (Extended Figure 2). To further test this supposition, we evaluated the expression levels of all 14 cell-surface receptor families during PTI and NLR-triggered immunity (NTI) in *Arabidopsis thaliana*^15,16^. Notably, RLKs, except for LRR- and Malectin-containing cell-surface receptors, generally exhibit higher expression levels compared to RLPs during immunity (Main figure 1j; supplementary figure 1).

### The origin and expansion of cell-surface receptors in the plant lineage

To trace the origins of different receptor classes within the plant lineage, we examined the presence of ectodomains (LRR-, G-lectin-, WAK-, Duf26-, L-lectin-, LysM- and Malectin-ectodomains lacking TM or KD), RLPs (TM-bound ectodomains) and RLKs (ectodomains encompassing both TM and KD) in Glaucophyta, red algae, green algae, Bryophytes and Tracheophytes (Main figure 2; Supplementary figure 1). Ectodomains exhibit an ancient heritage, with LRR-, WAK-, LysM-, Malectin-, and L-lectin-domains dating back to the era of Glaucophyta. Similarly, relatively ancient counterparts such as LRR-RLPs, WAK-RLPs, LysM-RLPs, Malectin-RLPs, and L-lectin-RLPs are found in both Glaucophyta and Rhodophyta. In contrast, RLKs emerged more recently. Green algae harbour WAK-RLKs, Malectin-RLKs, and G-lectin-RLKs, and LysM-RLKs, L-lectin-RLKs, and Duf-26-RLKs are exclusive to Embryophytes (Main figure 2).

**Main figure 2.**
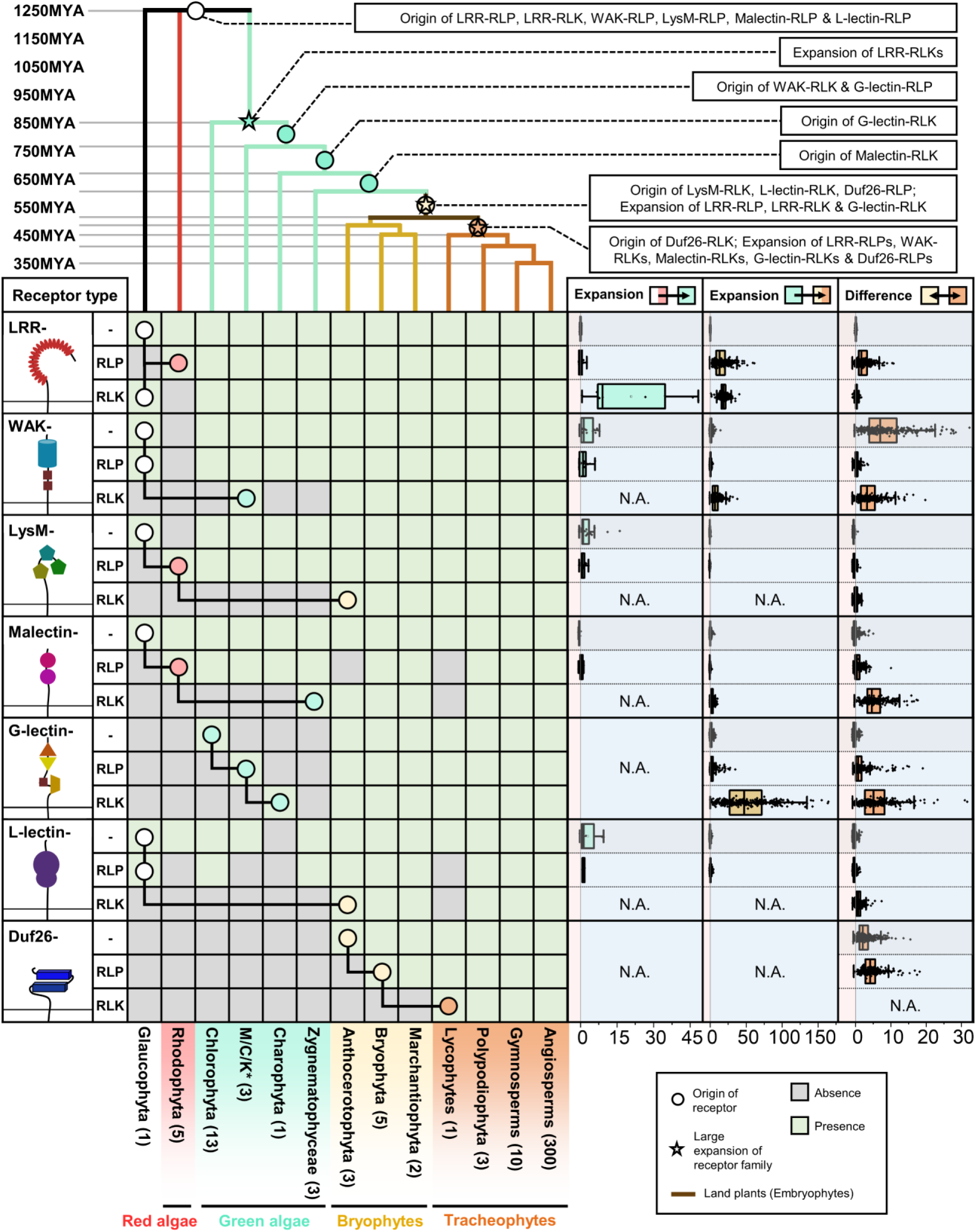
The origin and expansion of cell-surface receptors in plants. The top panel represents a phylogenetic tree of multiple algal and plant lineages. Circles (○) and stars (⋆) indicate the origin and expansion of receptor families. The timescale (in millions of years; MYA) of the phylogenetic tree was estimated by TIMETREE5^23^. The bottom panel represents the presence or absence of different receptor classes in algal and plant lineages. ‘-’ represents ectodomains with no transmembrane or kinase domain, ‘RLP’ represents ectodomains with a transmembrane domain but no kinase domain, ‘RLK’ represents ectodomains with both transmembrane and kinase domains. *M/C/K represents Mesostigmatophyceae, Chlorokybophyceae, and Klebsormidiophyceae. The number of species available from each algal and plant lineage are indicated by the numbers within respective boxes. A grey box indicates the absence of receptors and a green box indicates their presence in each lineage. The origin of a receptor is indicated with a circle (○). The origins of ‘-’, ‘RLP’, and ‘RLK’ are connected by black lines. Expansion rates of receptor classes are indicated by boxplots. The cyan boxplot represents the expansion rate from Glaucophyta and Rhodophyta to green algae. The yellow boxplot represents the expansion rate from green algae to Embryophytes and the orange boxplot represents the expansion rate from early land plants to Tracheophytes. Light blue area represents expansion and light pink area represents contraction of the gene family. X-axis values represent expansion rate (×). Boxplot elements: center line, median; bounds of box, 25th and 75th percentiles; whiskers, 1.5 × IQR from 25th and 75th percentiles.

Except for LRR-RLPs, all six families of RLP are basal to the RLK families. This suggests the intriguing possibility that numerous RLKs may have evolved directly from RLPs through the integration of kinase domains. Such integration events likely occurred at various stages of plant evolution, given the presence of RLKs in diverse plant lineages. Alternatively, it remains plausible that RLKs are equally ancient as RLPs, and that their absence from certain lineages is a result of loss over time. Nevertheless, it is highly probable that both RLPs and RLKs originated from ectodomains that underwent multiple gene duplication and domain-swapping events to incorporate TM and/or kinase domains, thereby diversifying their functional repertoire^22^.

We also examined the expansion patterns of different receptor classes across various plant lineages. Our analysis involved in calculating the median percentage (%) of cell-surface receptors in i) Glaucophyta and Rhodophyta, ii) green algae, and iii) Bryophytes and then determining the percentage increase (% increase) from Glaucophyta and Rhodophyta to green algae; from green algae to Embryophytes; and between Bryophytes and Tracheophytes. We observed substantial expansions in specific receptor families across these lineages. Green algae exhibited a significant expansion of LRR-RLKs, while Embryophytes displayed expansions in LRR-RLPs, LRR-RLKs, WAK-RLKs, and G-lectin-RLKs. Tracheophytes had further expansions in LRR-RLPs, WAK-RLKs, Malectin-RLKs, G-lectin-RLKs and Duf26-RLPs (Main figure 2; Supplementary figure 1). Overall, RLKs demonstrate greater expansion compared to RLPs, with notable expansions observed in LRR-RLK, WAK-RLK, and G-lectin-RLK. These findings align with the substantial size of these receptor families and their involvement in recognising PAMPs (Main figure 1h; Extended Figure 2a). In particular, the LRR-RLK-XII subgroup exhibits a considerably higher expansion rate compared to other subgroups (Extended Figure 3; Supplementary figure 1), reinforcing the idea that cell surface immune receptors underwent extensive expansions as the plant lineage diversified and evolved to adapt to a wide range of environments. It is worth noting that LRR-RLKs and LRR-RLPs have been reported to recognise a number of different types of ligands such as peptides, full-length proteins, small molecules, and phytohormones, whereas WAK-RLKs perceive glycans and full-length proteins, and G-lectin-RLKs perceive lipids and full-length proteins (Extended Figure 2a). Thus, it is plausible that these receptor families expanded throughout plant evolution to accommodate a broader range of ligands.

### The origin and expansion of PTI-signalling components in the plant lineage

Upon ligand perception, PRRs engage in homodimerization or heteromeric complex formation with co-receptors, such as BAK1 (a member of the Somatic embryogenesis receptor-like kinases (SERK) family) and SOBIR1, to initiate downstream signalling cascades. Cytoplasmic kinases, such as RLCK-VIIs, Mitogen-activated protein kinases (MAPKs), and Calcium-dependent protein kinases (CDPKs), undergo phosphorylation to transmit immune signals. Likewise, PM-localised downstream signalling components, such as cyclic nucleotide-gated channels (CNGCs), hyperosmolality-gated calcium-permeable channels (OSCAs), and the respiratory burst oxidase homolog NADPH oxidases (RBOHs) are also phosphorylated to elicit a defense response^9^. We identified these co-receptors and signalling components from 350 genomes and determined their absence or presence across the plant lineage (Extended Figure 4; Supplementary figure 2-26). SERKs, acting as cell-surface co-receptors for multiple LRR-RLKs and LRR-RLPs are present in Zygnematophyceae and Embryophytes^24^ (Main figure 3a), suggesting their emergence during or prior to the appearance of land plants. Immune-related LRR-RLPs lack intracellular kinase domains, thus require another LRR-RLK co-receptor, SOBIR1, to activate downstream signalling^25^. Similar to BAK1, SOBIR1 is also present in Embryophytes (Main figure 3a). Thus, co-receptors for cell-surface receptors likely evolved during or before the emergence of land plants. On the other hand, cytoplasmic kinases (RLCKs, CDPKs, MAPKKKs, MAPKKs, and MAPKs) are ancient, as are the PM-localised downstream signalling components (CNGCs, OCSAs, and RBOHs), found across all plant lineages (Main figure 3a). Although the exact function of these proteins in algal species remains unclear, their immune-related orthologs are present in green algae (Main figure 3; Supplementary figure 2-26). This suggests that they underwent specialisation within the immune activation pathway prior to the emergence of land plants (Extended Figure 4). The EP proteins (EDS1, PAD4, and SAG101) and RPW8-NLRs (NRG1 and ADR1) that are essential for both TIR-NLR and LRR-RLP mediated-immunity^26,27^, are only present in gymnosperms and angiosperms (seed plants)^28^. Considering the ancient nature of LRR-RLPs, it is plausible that EP-protein and helper-NLRs were integrated into the LRR-RLP-signalling pathway, forming a robust immune network in seed plants.

**Main figure 3.**
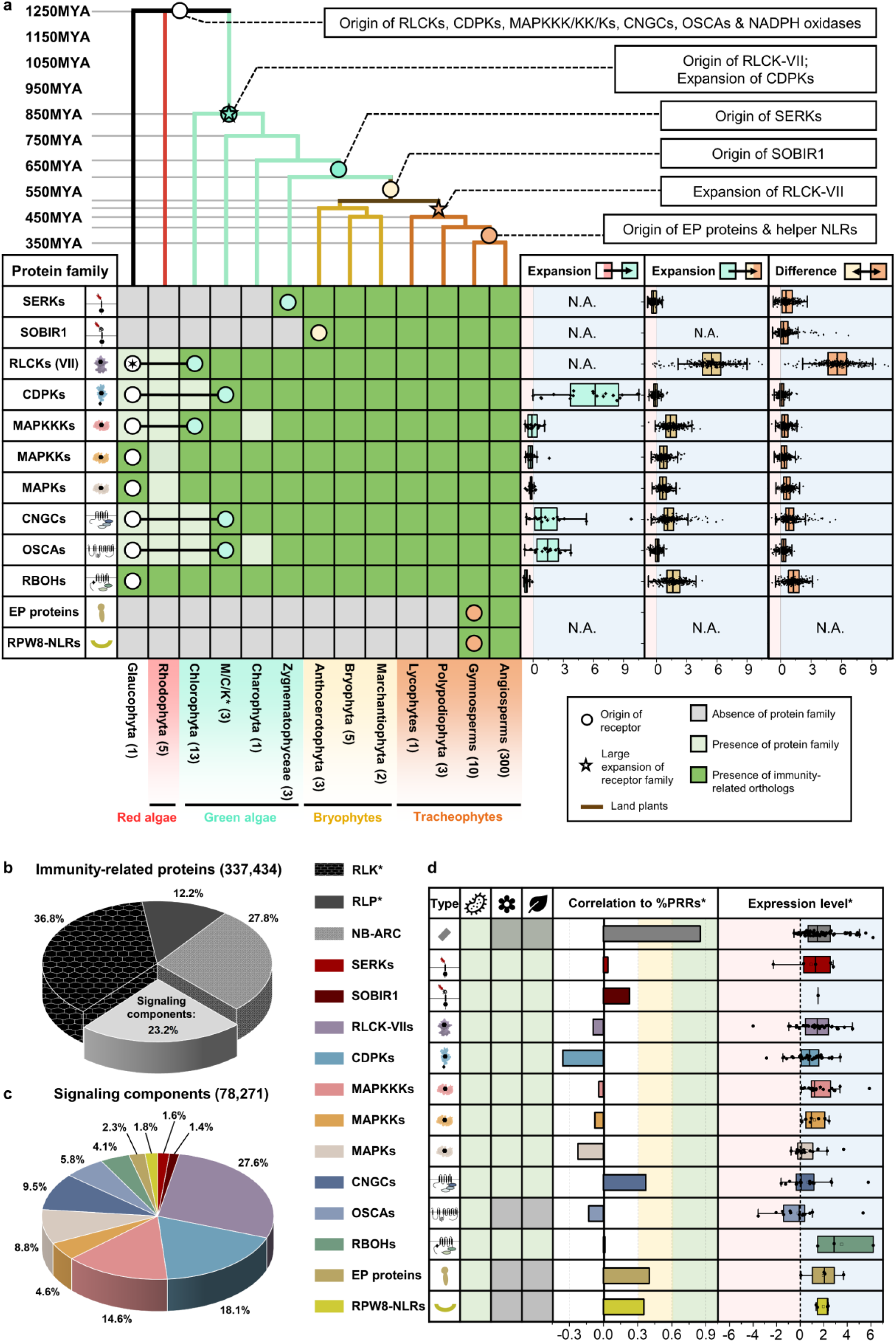
The origin and evolution of PTI-signalling component in plants. **(a)** The top panel is a phylogenetic tree of multiple algal and plant lineages. Circles (○) and stars (☆) indicate the origins and expansion of receptor families, respectively. The timescale (in million year; MYA) of the phylogenetic tree was estimated by TIMETREE5^23^. The bottom panel displays the presence or absence of receptor classes in different algal and plant lineages. *M/C/K represents Mesostigmatophyceae, Chlorokybophyceae, and Klebsormidiophyceae. The number of available species from each algal and plant lineage is indicated within the respective boxes. A grey box indicates the absence, while a green box indicates the presence of a given protein family in each lineage. Dark green indicates the presence of orthologs of immunity-related (PTI) signalling components within that protein family (see also Supplementary figure 4-26). The origin of a protein family is indicated with a circle (○), followed by another circle indicating the origin of the orthologs of PTI-signalling component. The expansion rates of the protein family members are indicated by boxplots. The cyan boxplot represents the expansion rate from Glaucophyta and red algae to green algae. The yellow boxplot represents the expansion rate from green algae to Embryophytes, and the orange boxplot represents the expansion rate from early land plants to Tracheophytes. Light blue area represents expansion and light pink area represents contraction of the gene family. X-axis values represent expansion rate (×). **b)** Distribution of RLKs, RLPs, NB-ARCs, and PTI-signalling components in plants. Each fraction represents the percentage (%) of ectodomain out of all four protein classes from 350 species (337,434). *Note that RLKs and RLPs here represent RLKs and RLPs with LRR-, G-lectin-, L-lectin-, LysM-, Malectin-, WAK- and Duf26-ectodomains. **c)** Distribution of PTI-signalling components in plants. Each fraction represents the percentage (%) of each protein family out of all the families combined (78,271). The colour codes for **b)** and **c)** are indicated on the right. **d)** Characterised PTI-signalling components involved in microbial interaction (bacteria icon), reproduction (flower icon), and development (leaf icon) are indicated with green boxes. A grey box indicates that receptor class has not been reported to be involved in that particular biological process (see also Extended Figure 4). The sizes of each signalling component family are given as a percent of all the proteins in the genomes of 350 plant species. Correlations between different classes of signalling components and PRR* in 300 angiosperms are indicated by the lengths of bars. *Note that PRRs here represent the combined sum of LRR-RLK-XII and LRR-RLPs, since many characterised members of these two classes are involved in pathogen recognition. A moderate positive correlation is represented in yellow (Pearson’s r between 0.3-0.6). Expression level* refers to the expression of each class of PTI-signalling component during NTI in *A. thaliana*. The blue shade represents increased expression, and the red shade represents decreased expression during NTI. The X-axis represents log_2_ (fold change during NTI relative to untreated samples). RNA-seq data analysed here were reported previously^16^, where NTI was activated by estradiol-induced expression of *AvrRps4* in *A. thaliana* for 4 hours. For the expression of each class of PTI-signalling component during PTI in *A. thaliana*, refer to supplementary figure 27. The boxplot elements in **(a)** and **(d)**: centre line, median; bounds of box, 25th and 75th percentiles; whiskers, 1.5 × IQR from 25th and 75th percentiles.

Our investigation of the expansion rate of signalling components within the plant lineage indicated an expansion of CDPKs in green algae and expansion of RLCK-VIIs in Tracheophytes (Main figure 3a; Supplementary figure 6-9). However, other families of signalling components exhibit more limited expansions, compared to PRRs. This is also consistent with the considerably larger family sizes of cell-surface receptors and NLRs in comparison to the signalling components (Main figure 3b-c). Furthermore, we examined the correlation between the percentages of signalling components and PRRs (LRR-RLK-XIIs + LRR-RLPs) across genomes. With the exception of CNGCs, EP proteins, and RPW8-NLRs (0.6 > Pearson’s r > 0.3), most signalling component families do not exhibit co-expansion or co-contraction with PRRs (Main figure 3d). Thus, the expansion of PRR gene families does not necessarily result in the expansion of PTI signalling components. The RLCK-VIIs are further classified into ten subgroups which are differentially required for RLKs and RLPs to activate downstream responses^29,10,30–32^ (Supplementary figure 6-7). Similarly, CDPKs fall into 4 subgroups (Supplementary figure 8-9). RLCK-VII subgroup families are differentially required by different PRRs to activate downstream immune responses^33–36^. Pathogens often target RLCK-VIIs through secreted effectors to suppress immunity^37–39^. Thus, redundancy among RLCK-VII subgroups serves as a protective mechanism for the downstream signalling pathway against effector targeting. In addition, plants have evolved RLCK-VII pseudokinases, or ‘decoys’, to guard functional RLCKs through NLRs^37,40–43^. Together, it has become apparent that the expansion of RLCK-VII families may have been driven by pathogenic pressure, thereby contributing to the enhanced robustness of the immune signalling network.

### The origin of immunity-related LRR-RLPs

To understand the origin of immune-specificity in cell-surface receptors, we sought to trace the evolutionary origin of LRR-RLPs in plants. Among RLPs, LRR-RLPs constitute the largest family, comprising more than twenty characterised members that perceive PAMPs or apoplastic effectors to activate immunity (Extended Figure 5). The ectodomain of LRR-RLPs encompasses additional domains known as N-loopouts (NLs) and island domains (IDs) interspersed between the LRR motifs. Typically, NLs are located closer to the N-termini of the ectodomain, whereas IDs are positioned closer to C-termini. NLs are present in most immunity-related LRR-RLPs while IDs are present in all immunity-related LRR-RLPs (Extended Figure 5a; Supplementary figure 28). NL positioning is relatively more flexible, occurring either before the first LRR motif or between the first few LRR motifs. Conversely, ID positioning is less flexible, located mostly before the last 4 LRR motifs within the ectodomain (Extended Figure 5a; Supplementary figure 28). This observation implies a functional necessity for the specific placement of IDs in LRR-RLPs.

To investigate the functional necessity of IDs, we analysed ectodomains of LRR-RLPs and LRR-RLKs from 350 species (113,794) (Extended Figure 6a-b). Employing multiple prediction programs, we identified gaps between LRR motifs ranging from 10-29 or 30-90 amino acids (AA) (Extended Figure 6a). Since NLs typically span 6-30AA and IDs range from around 40-75AA, we focused on small gaps (10-29AA) corresponding to NLs, and large gaps (30-90AA) indicative of IDs (Extended Figure 6a and supplementary information). Small or large gaps are relatively infrequent in LRR-RLKs (10.6% and 5.43%, respectively) (Extended Figure 6c). In contrast, both small and large gaps are more prevalent in LRR-RLPs (28.3% and 61.6%, respectively) (Extended Figure 6c). Furthermore, both LRR-RLKs and LRR-RLPs typically have only one gap, which can be either small or large (Extended Figure 6d). Our analysis also showed that small gap positions within the ectodomains of both LRR-RLKs and LRR-RLPs are not fixed, and may be distributed randomly. Conversely, larger gaps are predominantly positioned before the last four LRR motifs in the ectodomain (51.2% for LRR-RLKs and 86.9% for LRR-RLPs) (Extended Figure 6e-f; Supplementary figure 28-32). Thus, our findings suggest functional requirement for IDs to be positioned before the last four LRRs.

An analysis of the distribution of LRR-RLK subgroups with IDs indicated that over 55% belong to the Xb subgroup (Main figure 4a). Furthermore, 94.7% of LRR-RLKs with IDs positioned before the last four LRRs (ID+4LRR) belong to the Xb subgroup (Main figure 4a), suggesting that both LRR-RLK-Xb and LRR-RLPs share the ID+4LRR motif. Among the characterised members of LRR-RLK-Xb are important components of growth and development regulation, including the BRASSINOSTEROID INSENSITIVE 1 (BRI1) family members, PHYTOSULFOKIN RECEPTOR 1 (PSKR1) and PSY1 RECEPTOR (PSY1R) family members, EXCESS MICROSPOROCYTES1 (EMS1) and NEMATODE-INDUCED LRR-RLK 1 (NILR1/GRACE)^44–48^. Interestingly, we observed that both LRR-RLPs and LRR-RLK-Xbs with ID+4LRR motifs are present in land plants (Embryophyte) but not in other lineages (Main figure 4b). Considering the similarity in structural motifs between LRR-RLPs and LRR-RLK-Xbs, it is likely that these two receptor classes share a common origin.

**Main figure 4.**
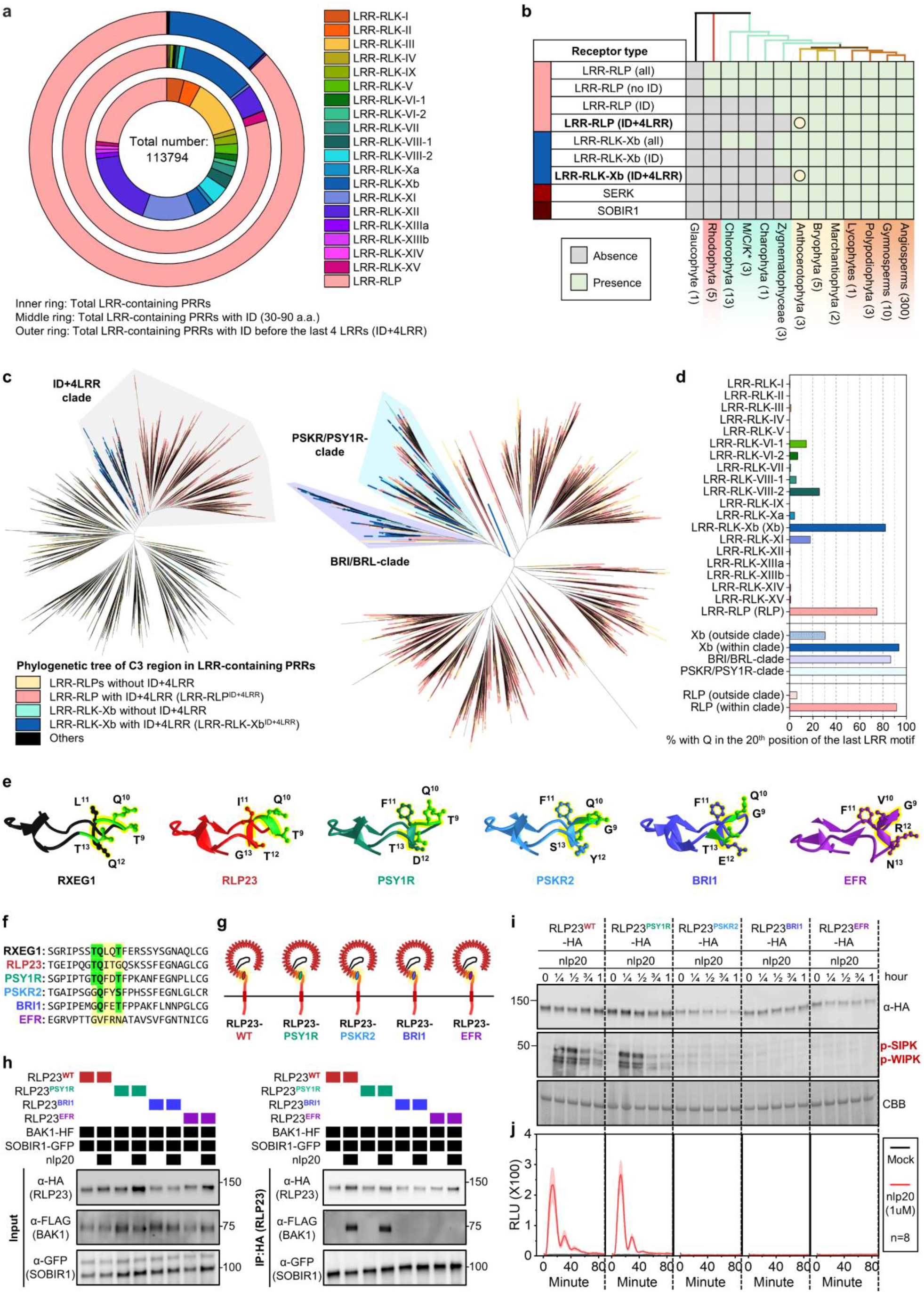
The origin and evolution of LRR^ID+4LRR^ in plants. **(a)** The concentric ring pie chart presents the percentage of LRR-containing cell-surface receptors (PRRs) from 350 species. The inner ring represents all LRR-containing cell-surface receptors (113,794); the middle ring represents LRR-containing PRRs with ID (20,556); the outer ring represents LRR-containing PRRs with an ID preceding the last 4 LRR (ID+4LRR) at the C terminus (16,885). **(b)** The presence or absence of receptor classes in various algal and plant lineages. *M/C/K represents Mesostigmatophyceae, Chlorokybophyceae, and Klebsormidiophyceae. A grey box indicates the absence, and a green box indicates the presence of a given receptor class in each lineage. The origin of LRR-RLP and LRR-RLK-Xb with ID+4LRR is indicated with a circle (○). **(c)** Phylogenetic tree of the C3 region (last 4 LRRs) from all LRR-containing PRRs of 350 species. Branches are colour-labelled as indicated. The light grey area indicates clustering of LRR-RLK-Xb and LRR-RLP with ID+4LRR. The pruned phylogenetic tree on the right corresponds to the light grey area in the left tree, with clades labelled in dark grey areas accordingly. **(d)** The percentage of receptors with glutamine (Q) in the 9^th^ position of the terminal LRR. Percentages (%) are calculated as (the number of LRR-RLKs or LRR-RLPs in the subgroup with Q) / (the number of LRR-RLKs or LRR-RLPs in the subgroup without Q) ×100. RLP/RLK-Xb (outside clade) refers to receptors outside the light grey clade in **(c)**. RLP/RLK-Xb (within clade) refers to receptors inside the light grey clade in **(c)**. The number of cell-surface receptors (n) in each LRR-RLK subgroup: I, n = 752; II, n = 682; III, n = 6572; IV, n = 1033; V,n = 8; VI-1, n = 84; V1-2, n = 146; VII, n = 1720; VIII-1, n = 195; VIII-2, n = 411; IX, n = 70; Xa, n = 96; Xb, n = 3182; XI, n = 8807; XII, n = 12863; XIIIa, n = 739; XIIIb, n = 465; XIV, n = 241; XV, n = 548; Xb (outside clade), n = 580; Xb (within clade), n = 2527; BRI/BRL clade, n = 1170; PSKR/PSY1R clade, n = 1347. The number of cell-surface receptors (n) in each LRR-RLP subgroup: LRR-RLP (RLP), n = 24860; RLP (outside clade), n = 5000; RLP (within clade), n = 19970. **(e)** Structure of the terminal LRR motif of *Nicotiana benthamiana* (Nb)RXEG1, *A. thaliana* (At)RLP23, AtPSY1R, AtPSKR2, AtBRI1 and AtEFR. Structures of NbRXEG1 and AtBRI1 were published^53–56,51,57^. Structures of AtRLP23, AtPSY1R, AtPSKR2 and AtEFR were predicted by Alphafold2*,^20^. Ectodomains are visualised in iCn3D^21^. **(f)** Alignment of amino acids in the last LRR motifs from NbRXEG1, AtRLP23, AtPSY1R, AtPSKR2, AtBRI1, and AtEFR. Amino acid residues involved in the interaction between NbRXEG1 and BAK1 are highlighted in green. The QxxT motif positions are highlighted in yellow. **(g)** Design of AtRLP23 chimeras. The last LRR motif of AtRLP23 is exchanged with either AtPSY1R, AtPSKR2, AtBRI1, or AtEFR. **(h)** Immuno-precipitation to test interactions between AtRLP23 chimeras, AtBAK1 and AtSOBIR1. Nb leaves expressing the indicated constructs were treated with either mock or 1μM nlp20 for 5 minutes. **(i)** Functionality testing of AtRLP23 chimeras. Nb leaves expressing the indicated constructs were treated with 1μM nlp20 and samples were collected at indicated time points. Phosphorylation of NbSIPK and NbWIPK was detected with p-P42/44 antibody. **(j)** Nb leaf discs expressing the indicated constructs were collected and treated with either mock or 1μM nlp20, and ROS production was measured for 90 min. For details of experiential design in **(h)**, **(i)**, and (**j)**, refer to the methods section.

To test this hypothesis, we conducted phylogenetic and structural analyses on the ectodomains of LRR-RLPs and LRR-RLKs. First, we aligned the last four LRR motifs (referred to as the C3 region^49^) from LRR-RLKs and LRR-RLPs (60,240). LRR-RLPs and LRR-RLK-Xbs with an ID+4LRR cluster together in the C3 phylogenetic tree, indicating a common origin (Main figure 4c). Within this cluster (indicated as the ID+4LRR clade), we observed distinct subclades, including the BRI1/BRI1-LIKE(BRL) clade, the PSKR/PSY1R clade, and mostly LRR-RLPs (Main figure 4c). The closely similar C3 regions indicate a conserved function. BRI1 recognises brassinosteroids (BRs) and interacts with the co-receptor BAK1 to induce BR responses^50,51^. Similarly, PSKR1 perceives phytosulfokine (PSK) and interacts with the co-receptor SERK1/2. LRR-RLPs, on the other hand, perceive PAMPs or apoplastic effectors and engage both co-receptor BAK1 and SOBIR1 to initiate immune responses^52^. Structural studies have elucidated the interaction mechanism of BRI1, PSKR1, and the LRR-RLP RXEG1 with SERKs^53–56,51,57^ (Extended Figure 6a-d). Interestingly, the C3 regions of BRI1, PSKR1, and RXEG1 play a crucial role in SERK interactions^55,56,51,57^, differentiating them from other LRR-RLKs. BRI1, PSKR1 and RXEG1 contain specific amino acid residues within the terminal two LRR motifs (within the C3 region) to interact with BAK1/SERK1 (Extended Figure 6b-d), while the flagellin peptide (flg22) receptor FLS2 (an LRR-RLK-XII), relies on the 3^rd^-12^th^ last LRR for BAK1 (SERK3) interactions^18^ (Extended Figure 6a). The N-terminal region (or N-terminal cap) of BAK1 or SERK1 is important for FLS2, BRI1, PSKR1, and RXEG1 interactions, while the 1^st^ LRR inner surface of BAK1 or SERK1 is primarily involved in BRI1, PSKR1, and RXEG1 interactions, and the 2^nd^ and the 4^th^ LRR inner surface of BAK1 is involved in FLS2 interactions (Extended Figure 6a-e). There are also striking similarities in interaction network maps between PSKR1-SERK1 and RXEG1-BAK1 interactions. Additionally, the C3 region of BRI1 participates in both BR binding^55,56^ and co-receptor binding, whereas the C3 region of PSKR1 and RXEG1 exclusively engages in co-receptor interactions (SERK1/2 and BAK1, respectively)^51,57^. By aligning the C3 regions of various LRR-RLKs, including BRI1/BRL-orthologs, PSKR orthologs, PSY1R, and multiple LRR-RLPs, overlapping residues that are required for SERK interactions within PSKR/PSY1R, and LRR-RLP clades can be discerned (Extended Figure 8a). For example, the glutamic acid (E) residue at the second position of the penultimate LRR motif is involved in both PSKR1-SERK1 and RXEG1-BAK1 interactions^57^. This E residue is highly conserved in both clades (PSKR/PSY1R clade: 69.9%, LRR-RLP: 86%; Extended Figure 8b). Similarly, the phenylalanine (F) residue at the last position of the penultimate LRR motif contributes to both PSKR1-SERK1 and RXEG1-BAK1 interactions^51^. This F residue is conserved in the BRI1/BRL (62.2%), PSKR/PSY1R (99.7%), and LRR-RLP (64.1%) clades (Extended Figure 8b). There is also a conserved motif crucial for SERKs interactions within the last LRR motifs of BRI/BRL, PSKR/PSY1R, and LRR-RLP clades. This motif, Glutamine-x-x-Threonine/Serine (QxxT/S) loop, is conserved in BRI1/BRL (Q:86.6%; T/S:98.9%), PSKR/PSY1R (Q:99.9%; T/S:92.5%) and LRR-RLP (Q:91.8%; T/S:88.4%) clades, but it is not conserved in other LRR-RLPs outside of the ID+4LRR clade or LRR-RLK subgroups (Main figure 4d; Extended Figure 8a). Structural analysis of BRI1-BAK1, PSKR1-SERK1/2, and RXEG1-BAK1 further supports the importance of residues within and around the QxxT/S loop for SERK interactions^55,56,51,57^ (Extended Figure 7b-d). In conclusion, our findings indicate that the C3 regions of LRR-RLK-Xb and LRR-RLPs share a conserved function for interacting with SERKs.

To further validate the functional conservation of C3 regions in LRR-RLK-Xb and LRR-RLPs, we performed functional analysis of the QxxT/S motif in the LRR-RLP RLP23 from *A. thaliana*. RLP23 forms heteromeric complexes with the LRR-RLK co-receptor SOBIR1, and upon the perception of the nlp20 peptide, BAK1 is recruited into the complex, leading to activation of the SOBIR1 KD to induce immunity^25,52,58^. Similar to RXEG1 (with T<u>Q</u>LQ<u>T</u>), RLP23 possesses a TQITG motif (T<u>Q</u>xxx), while LRR-RLK-Xb members, such as PSY1R, PSKR2, and BRI1, feature T<u>Q</u>FD<u>T</u>, G<u>Q</u>FY<u>S</u>, and G<u>Q</u>FE<u>T</u> motifs, respectively (all QxxT/S). Notably, the LRR-RLK-XII member EFR lacks the QxxT/S motif in that position, having GVFRN instead (Main figure 4e-f). We generated chimeric constructs of RLP23 with the terminal LRR motifs swapped between PSY1R, PSKR2, BRI1, and EFR (Main figure 4g). By immuno-precipitation assays, we tested the ability of these chimeras to interact with BAK1 upon ligand perception. Both wildtype RLP23 (RLP23^WT^) and RLP23-PSY1R (RLP23^PSY1R^) chimeras can interact with BAK1 upon nlp20 treatment, whereas RLP23-BRI1 (RLP23^BRI1^) and RLP23-EFR (RLP23^EFR^) cannot. This suggests that the terminal LRR motif of RLP23 and PSY1R bind BAK1 in a similar manner, but the terminal LRR motif of BRI1 binds to BAK1 differently. Importantly, all chimeras can interact with SOBIR1 regardless of the presence of nlp20, indicating that the last LRR motif may not be involved in SOBIR1 interactions (Main figure 4h). Furthermore, upon nlp20 treatment, both RLP23^WT^ and RLP23^PSY1R^ can trigger immune responses, while RLP23^PSKR2^ and RLP23^BRI1^ cannot (Main figure 4i). We speculate that this may be due to the absence of a specific T residue before the Qxxx motif in RLP23, which is relatively less prevalent within the LRR-RLP clade (31.1%; Extended Figure 8). Multiple *A. thaliana* LRR-RLPs contain this residue, but it is not in other studied LRR-RLPs, such as the tomato Cf proteins (Extended Figure 8), suggesting that this residue evolved in some species after the divergence of LRR-RLK-Xb and LRR-RLPs. Nevertheless, our results strongly support the functional conservation of C3 regions in LRR-RLK-Xbs and LRR-RLPs, specifically their ability to interact with SERKs.

The conservation of other regions within the ectodomain of LRR-RLK-Xb and LRR-RLPs was tested by aligning the IDs extracted from LRR-RLKs and LRR-RLPs (20,246). Remarkably, the ID clusters of LRR-RLPs and LRR-RLK-Xb are found in close proximity, mirroring the C3 phylogenetic tree (Extended Figure 9a). Again, the BRI1/BRI1-LIKE(BRL) and PSKR/PSY1R clusters are close to the LRR-RLPs (Extended Figure 9a). These results are also consistent with a previous report that PSKRs are closely related to some LRR-RLPs in Arabidopsis and rice^49^. A recent review identified two conserved lysine (K)-containing motifs, Yx8KG and Kx5Y, in the ID of LRR-RLPs^59^. The lysine residue in the Kx5Y motif from RXEG1 is required for its interaction with BAK1^57^. We extracted both Yx8KG and Kx5Y motifs in the extracted IDs (before the last 4LRR motifs) of LRR-RLKs and LRR-RLPs. More than 75% of IDs from LRR-RLPs contain at least one of these lysine motifs, while less than 5% of IDs from LRR-RLK-Xb have either motif (Extended Figure 9b). This is consistent with structural data indicating that IDs from BRI1 and PSKR1 employ distinct residues for their interactions with BAK1^55,56,51^. However, IDs from LRR-RLPs that are closely related to those from LRR-RLK-Xbs retain the Kx5Y motifs (Supplementary figure 33). It is therefore possible that the common ID ancestor may have originally harboured lysine motifs that were subsequently lost from the BRI1 and PSKR/PSY1R clades.

To further explore the relatedness of the IDs between LRR-RLK-Xb group and LRR-RLPs, we formed clusters of highly similar IDs and examined their subgroup/family affiliations. Overall, we identified more than 2,822 clusters, with the majority (2,734) consisting of LRR-receptors of a single subgroup/family (Extended Figure 9c). Among the 61 clusters containing LRR-receptors from two different subgroups/families, 80% (49 clusters) consisted of LRR-receptors from LRR-RLK-Xb and LRR-RLPs (Extended Figure 9d). The enrichment of LRR-RLK-Xb and LRR-RLP pairings provides further evidence for the relatedness of their ectodomains. The substantial number of LRR-RLP-only ID clusters (2,383) also suggests that the IDs of LRR-RLPs have undergone extensive diversification, in the process providing a broad scope for the recognition of PAMPs and apoplastic effectors. Most of clusters contain relatively a small number of IDs, predominantly from species within the same order or family (Extended Figure 9c). Consequently, tracing back the origin of IDs is challenging due to their considerable diversity. We therefore propose that the IDs of LRR-RLK-Xbs and LRR-RLPs likely originated from a common ancestor, with the IDs of LRR-RLPs expanding and diversifying after the divergence of LRR-RLK-Xb and LRR-RLPs.

Within the BRI1/BRI1-LIKE (BRL) and PSKR/PSY1R clusters, we found LRR-RLP counterparts of BRI1, PSKR, and PSY1R. To further test their relatedness, we aligned the ectodomains from LRR-RLKs and LRR-RLPs and extracted the branches containing BRI1, PSKR, and PSY1R from the phylogenetic tree (Extended Figure 9d-f; Supplementary figure 34). Within the BRI1/BRL-ectodomain clade, we identified 63 LRR-RLPs and 1,188 LRR-RLK-Xbs. The IDs of these LRR-RLPs are highly similar to BRL1/BRL3 with residues crucial for BL binding^53,54^ (Extended Figure 9e). Within the PSKR-ectodomain subclade, we found 343 LRR-RLPs alongside 791 LRR-RLK-Xbs. Similarly, the IDs of the LRR-RLPs are remarkably similar to PSKR2, with residues essential for PSK binding and SERK1/2 interactions^51^ (Extended Figure 9f). Finally, within the PSY1R-ectodomain subclade, we found 454 LRR-RLPs accompanied by 227 LRR-RLK-Xbs. The IDs of these LRR-RLPs share remarkable similarity with AtPSY1R (Extended Figure 9g). Moreover, AtRLP3 and AtPSY1R have over 70% sequence identity and 85% similarity in their ectodomains. Although PSY1R is not the receptor for PSY peptide^60^, it is possible that RLP3, and PSY1R recognise similar or identical ligands^61^. Currently, the functions of these BRL-, PSKR- and PSY1R-like LRR-RLPs remain unclear. RLP3 confers resistance against the vascular wilt fungus *Fusarium oxysporum* f. sp. *matthioli*^61^, while PSY1R is involved in growth and development^46^. We propose that these RLPs may either recognise endogenous molecules to activate growth and development, or participate in the recognition of pathogen-mimicking molecules to trigger immune signalling.

### Specialisation of cell-surface receptors in different biological processes

Given the common origin of the ectodomains of LRR-RLPs and LRR-RLK-Xbs, it is likely that these receptors have undergone specialisation in immune- and developmental processes following their divergence. While LRR-RLK-Xbs activate downstream responses through the Xb kinase domain, LRR-RLPs recruit SOBIR1 to trigger immunity^52,58^. Because LRR-RLPs lack a kinase domain, the juxtamembrane (JM) and TM regions may activate immune responses. We therefore aligned the eJM (JM region before TM), TM, and cJM (JM region after TM, but with the absence of kinase domain from RLKs) from LRR-RLKs and LRR-RLPs (62,896). Interestingly, the eJM-TM-cJM region of LRR-RLPs is not closely related to LRR-RLK-Xbs (Main figure 5a). Previous studies have reported the requirement of Glycine-X-X-X-Glycine (GxxxG) motifs in SOBIR1 for its association with LRR-RLPs, and the potential contribution of negatively charged amino acids in the eJM region of LRR-RLPs to their interaction with SOBIR1^62^. We aligned the eJM-TM region of LRR-RLKs, BRI1/BRL-orthologs, PSKR orthologs, PSY1R, and multiple LRR-RLPs, and analysed them for the presence of positively/negatively charged amino acids in eJM and GxxxG motifs in TMs (Extended figure 10a). Consistent with previous reports^62^, LRR-RLPs are strongly negatively charged at the end of the eJM region, whereas LRR-RLKs, including SOBIR1, are positively charged at the end of the eJM region (Extended figure 10a). Most LRR-RLPs have a single GxxxG motif, with some having two (GxxxGxxxG) or three (GxxxGxxxGxxxG) consecutive motifs. Conversely, GxxxG motifs are relatively less common in LRR-RLKs, but can be found in SOBIR1, BAK1, and the PSKR/PSY1R clade (Extended figure 10a). We further examined overall charges in the eJM region of the LRR-RLP and LRR-RLK subgroups. All LRR-RLK subgroups, except for LRR-RLK-VIII-2, feature positively charged eJM regions, whereas LRR-RLPs have negatively charged eJM regions. Importantly, negatively charged eJM regions are only present in LRR-RLPs within the ID+4LRR clade (Main figure 5b). The percentage of TMs with GxxxG or GxxxGxxxG motifs in the LRR-RLP and LRR-RLK subgroups varies from about 60% of subgroup members from LRR-RLK-I, LRR-RLK-VI-1, LRR-RLPs possess GxxxG motifs in TM, but more than 80% of LRR-RLPs within the ID+4LRR clade contain these motifs (Main figure 5b). Moreover, GxxxGxxxG motifs are primarily found in LRR-RLPs and are relatively rare in LRR-RLKs. Interestingly, both GxxxG and GxxxGxxxG motifs are enriched in the PSKR/PSY1R clade, indicating that the TM regions of LRR-RLPs and PSYR/PSY1R clade might share a common origin. (Main figure 5b; Supplementary figure 35-37).

**Main figure 5.**
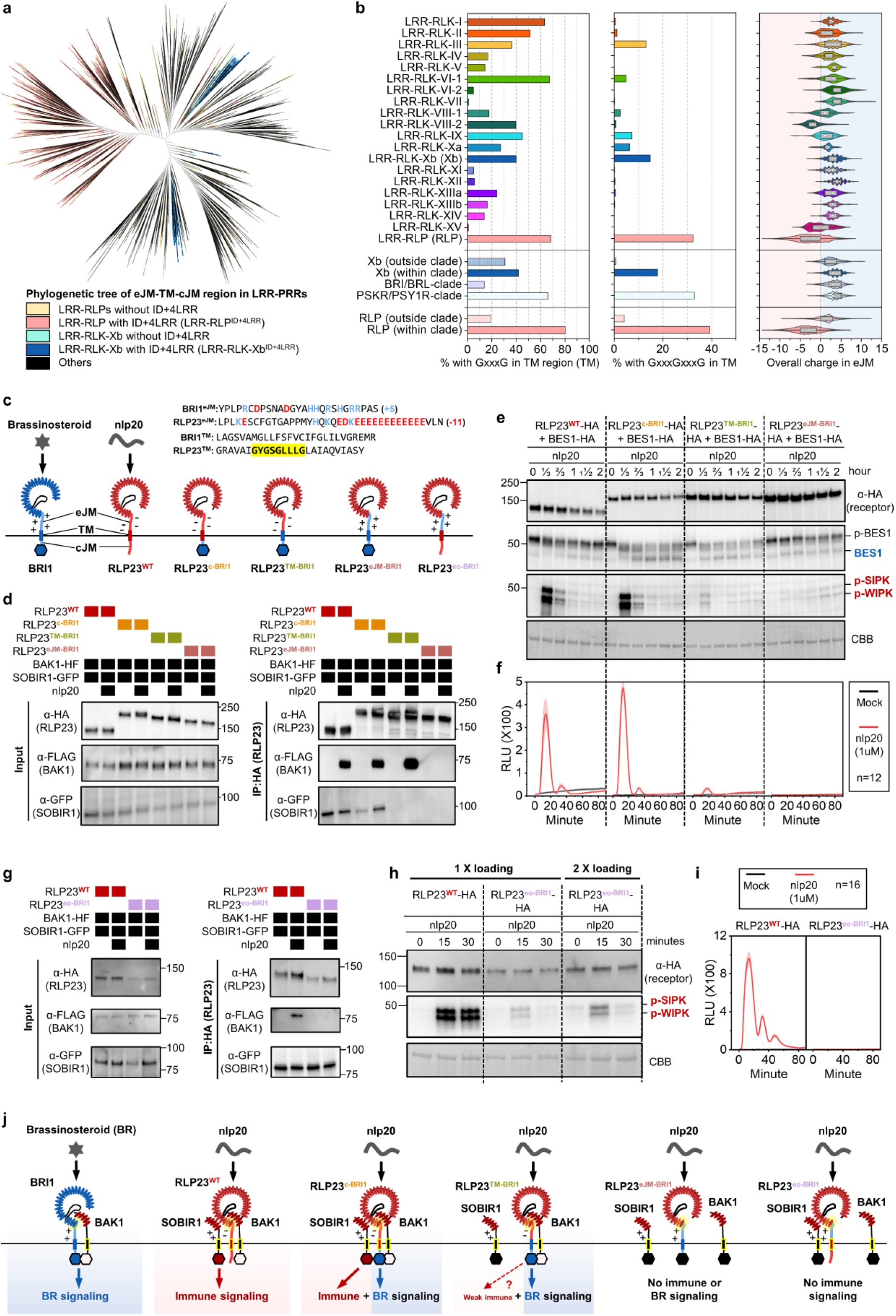
Adaptation of LRR^ID+4LRR^ to differential downstream signalling pathways. **(a)** Phylogenetic tree of eJM-TM-cJM regions from all LRR-containing PRRs of 350 plant species. Branches are labelled in colours as indicated. **(b)** Percentage of receptors with GxxxG in the TM region (left), GxGxG in the TM region (middle) and overall charge distribution in eJM (right). LRR-RLP/RLK-Xb (outside clade) refers to receptors outside the light grey clade in Figure 4c. Percentages (%) are calculated by the number of LRR-RLKs or LRR-RLPs in the subgroup with the stated residue divided by the number of LRR-RLKs or LRR-RLPs in the subgroup without the stated residue × 100. RLP/RLK-Xb (outside clade) refers to receptors outside the light grey clade in Figure 4c. RLP/RLK-Xb (within clade) refers to receptors inside the light grey clade in Figure 4c. Number of cell-surface receptors (n) in each LRR-RLK subgroup: I, n = 752; II, n = 682; III, n = 6572; IV, n = 1033; V, n = 8; VI-1, n = 84; V1-2, n = 146; VII, n = 1720; VIII-1, n = 195; VIII-2, n = 411; IX, n = 70; Xa, n = 96; Xb, n = 3182; XI, n = 8807; XII, n = 12863; XIIIa, n = 739; XIIIb, n = 465; XIV, n = 241; XV, n = 548; Xb (outside clade), n = 580; Xb (within clade), n = 2527; BRI/BRL clade, n = 1170; PSKR/PSY1R clade, n = 1347. Number of cell-surface receptors (n) in each LRR-RLP subgroup: LRR-RLP (RLP), n = 24860; RLP (outside clade), n = 5000; RLP (within clade), n = 19970. **(c)** Design of AtRLP23-BRI1 chimeras. Different regions (cytosolic, TM+cytosolic, and eJM+TM+cytosolic) of BRI1 were swapped into AtRLP23 as indicated. The alignment of amino acids in the eJM and TM regions from AtRLP23 and AtBRI1 is shown. Amino acid residues with negative charges are in red and amino acids with positive charges are in blue. The GxxxG motif in the TM region is highlighted in yellow. **(d, g)** Immuno-precipitation to test interactions between AtRLP23 chimeras, AtBAK1 and AtSOBIR1. Nb leaves expressing the indicated constructs were treated with either mock or 1μM nlp20 for 5 minutes. **(e, h)** Functionality testing of AtRLP23 chimeras. Nb leaves expressing the indicated constructs were treated with 1μM nlp20 and samples were collected at indicated time points. Dephosphorylation of BESI1-HA was detected with HA antibody. Phosphorylation of NbSIPK and NbWIPK was detected with p-P42/44 antibody. For **(h)**, twice the sample of RLP23^oe-BRI1^ was loaded as a reference, because RLP23^oe-BRI1^ protein accumulation is weaker than that of RLP23^WT^. **(f, i)** Nb leaf discs expressing the indicated constructs (without BES1-HA) were collected and treated with either mock or 1μM nlp20 and ROS production was measured for 90 minutes. For details of experiential design, refer to the methods section. **(g)** Schematic model of the interaction between LRR-RLK-Xb^ID+4LRR^ and LRR-RLP^ID+4LRR^ with co-receptors to induce differential downstream signalling. Both receptor classes utilise the last 4 LRRs (highlighted in yellow) to interact with SERKs (BAK1). LRR-RLP evolved to interact with SOBIR1 with the GxxxG motifs in TM (highlighted in yellow outline). Coloured hexagons on RLKs indicate activated kinases and black hexagon indicates an inactivated kinase.

To assess the functionality of eJM-TM-cytosolic regions in LRR-RLK-Xbs and LRR-RLPs, we generated multiple chimeras of eJM, TM, and cytosolic regions of BRI1 and RLP23 (Main figure 5c). In RLP23^c-BRI1^, the cytosolic region (following TM) of BRI1 was swapped into RLP23. In RLP23^TM-BRI1^, the TM+ cytosolic region of BRI1 was swapped into RLP23. In RLP23^eJM-BRI1^, the eJM + TM + cytosolic region of BRI1 was swapped into RLP23 (Main figure 5c). Immuno-precipitation assays showed that both RLP23^WT^ and RLP23^c-BRI1^ exhibit constitutive interactions with SOBIR1, and specifically interact with BAK1 upon nlp20 treatment. RLP23^TM-BRI1^ does not interact with SOBIR1 but still interacts with BAK1 upon nlp20 treatment, suggesting that the RLP23 TM region with GxxxG motifs is necessary for SOBIR1-, but not BAK1-interactions (Main figure 5d). Consistent with these immunoprecipitation assays, RLP23^WT^ and RLP23^c-BRI1^ can activate immune responses, while RLP23^c-BRI1^ and RLP23^TM-BRI1^ can activate developmental responses as indicated by the dephosphorylation of BRI1-EMS-SUPPRESSOR 1 (BES1)^63^ (Main figure 5e-f). This confirms that the BRI1 kinase domain is specifically required for the activation of BR responses. Interestingly, RLP23^TM-BRI1^ can also activate weak immune responses, likely independent of SOBIR1 (Main figure 5e-f). RLP23^eJM-BRI1^ does not interact with SOBIR1 or BAK1, and is unable to activate either immune or developmental responses. These data suggest that the eJM region of RLP23 is necessary for RLP23-BAK1 interactions (Main figure 5d). This conclusion was reinforced by generating RLP23^eo-BRI1^ chimera, in which the eJM region of RLP23 is replaced with the BRI1 eJM (Main figure 5c). RLP23^eo-BRI1^ consistently accumulates less protein than RLP23^WT^. Moreover, RLP23^eo-BRI1^ constitutively interacts with SOBIR1, but fails to interact with BAK1 with nlp20 treatment (Main figure 5g). RLP23^eo-BRI1^ only weakly activates MAPKs, and fails to trigger ROS production with nlp20 treatment (Main figure 5h-i). Thus, we conclude that the eJM regions of LRR-RLPs are important for protein accumulation and interaction with BAK1 upon ligand perception.

## Discussion

The ectodomains of LRR-RLPs and LRR-RLK-Xbs appear to share a common evolutionary origin, suggesting an ancestral adoption of the ID+4LRR architecture for ligand recognition and interactions with co-receptors such as the SERKs. Since both LRR-RLPs and LRR-RLK-Xbs rely on the terminal 4LRR/C3 region for SERK interaction^55,56,51,57^, it is plausible that the C3 region has undergone stabilizing evolution to preserve this functionality (Extended figure 10b). Consequently, a portion of the C3 region in LRR-RLK-Xb and LRR-RLP can be interchanged without loss of function. However, the remaining ectodomain, including the LRRs preceding the ID and the ID itself, has undergone diversification and adaptation to recognise distinct ligands. Notably, certain LRR-RLPs exhibit significant sequence similarities to LRR-RLK-Xbs (BRI1/BRL, PSKR, and PSY1R), though their specific functions are unexplored. Following diversification, the eJM, TM, and cytosolic regions of LRR-RLK-Xb and LRR-RLP acquired distinct roles, primarily in development and immunity, respectively (Main figure 5j; Extended figure 10b). The eJM and TM regions of LRR-RLP specifically facilitate constitutive interactions with SOBIR1 and ligand-dependent interactions with BAK1, whereas the LRR-RLK-Xb group lacks such specialisation. Since the recruitment of SOBIR1 to BRI1 would lead to immune activation, there should be negative selection for negatively charged eJMs and GxxxGs in the TM regions to prevent immune activation by LRR-RLK-Xbs. Given the different roles of various domains and regions of LRR-RLKs and LRR-RLPs, we propose a model in which different domains or regions of cell-surface receptors undergo modular evolution to either diversify or to maintain their original functions. Modular evolution allows the specialisation of cell-surface receptors to recognize different ligands and to activate distinct downstream signal responses or maintain interactions with co-receptors (Extended figure 10b).

The study of PRRs and the core PTI-signalling pathway has primarily focused on model plants, such as Arabidopsis and rice. Recent investigations into chitin perception by LysM-RLKs in the Bryophyte *Marchantia polymorpha* have revealed a high degree of conservation in the PTI-signalling pathway across land plants^64,65^. In fact, most cell-surface receptors and PTI signalling components in the Tracheophytes are conserved in Bryophytes, with the exception of Duf26-RLKs, EP proteins, and RPW8-NLRs (Main figure 6a). Therefore, the most recent common ancestor of land plants is likely to possess a considerable number of cell-surface receptors and basic components of immune signalling network, facilitating adaptation to terrestrial environments (Main figure 6a). The presence of diverse cell-surface receptor classes and multiple PTI signalling components in algal species suggests that algae may also have PTI system (Main figure 6a), as indicated by the MAMPs triggering of defense responses in some algal species. For instance, treatment with lipopolysaccharides (LPS) and oligosaccharides in multiple red algae species (Rhodophyta) stimulated ROS production, activation of nitric oxide (NO) signalling, changes in protein expression and hypersensitive responses (HR)^66^. Land plants perceive LPS and oligosaccharides through G-lectin- and WAK-/LysM-receptors ^67,68^. In Rhodophyta, we identified the presence of LysM-RLPs, but not WAK- or G-lectin-containing cell-surface receptors, suggesting that other PRRs perceive LPS in algae. Nevertheless, PTI is likely to be present in multiple algal lineages. Further investigation into PRRs and signalling network in the Glaucophyta, Rhodophyta, and green algal species holds promise for shedding light on the evolution PTI in Viridiplantae.

**Main figure 6.**
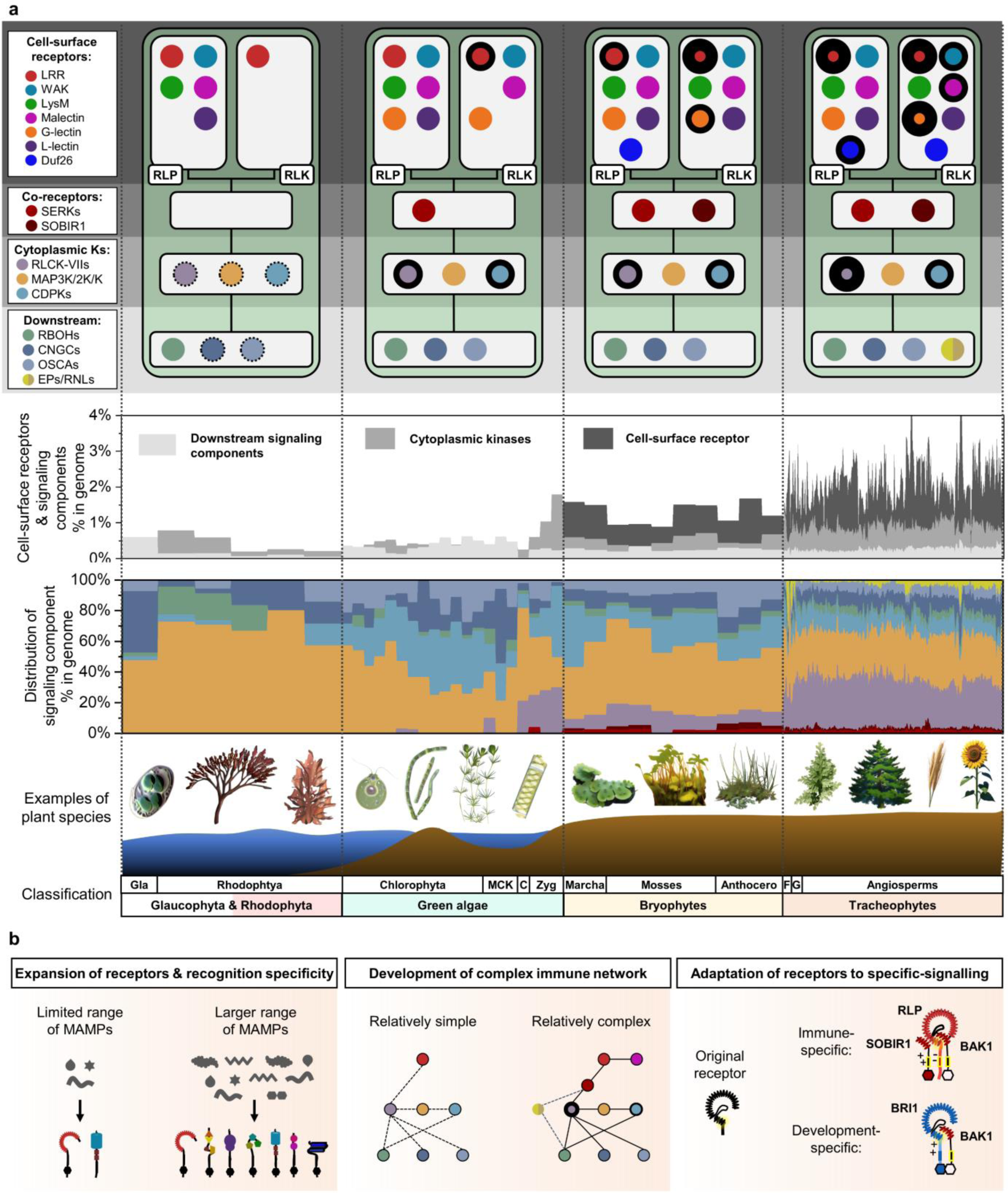
The evolutionary trajectory of PTI in plants. **(a)** The top panel depicts the presence of cell-surface receptors, co-receptors, cytoplasmic kinases (cytoplasmic Ks) and downstream signalling components (downstream) in Glaucophyta and Rhodophyta, green algae, Bryophytes and Tracheophytes. Nodes are labelled in colours as indicated on the left. Absence of a node indicates the absence of a gene family from the lineage. Nodes with dotted outlines indicate the presence of a gene family, but the absence of immunity-related orthologs. Nodes with thick outlines indicate the expansion of gene families. Repeated expansion is indicated with thicker outlines. The middle panel (top) displays the percentages (%) of cell-surface receptors and signalling components in the genome of each species within the lineage. Bars are labelled in colours as indicated. Middle panel (bottom) represents the distribution of signalling components, including co-receptors, cytoplasmic kinases, and downstream signalling components, in the genome of each species within a lineage. Bars are labelled in colours as indicated on top left. The bottom panel shows examples of plant species and the classification of the plant lineages. Abbreviations: Gla: Glaucophyta; MCK: represents Mesostigmatophyceae, Chlorokybophyceae, and Klebsormidiophyceae; C: Charophyta; Zyg: Zygnematophyceae; Marcha: Marchantiophyta; Mosses: Bryophyte; Anthocero: Anthocerotophyta; F: Lycophytes and Polypodiophyta; G: Gymnosperms. **(b)** Evolution of PTI in plants. Left panel: expansion of PRR family gene repertoires throughout the plant lineage, which leads to recognition of a larger range of MAMPs. Middle panel: PRR co-receptors, EP proteins and helper NLRs are absent from many algal species. A more complex immune network involving these signalling components apparently developed in vascular plants. Right panel: An ancient PRR with LRR^ID+4LRR^ with unknown function evolved into LRR-RLK-Xbs and LRR-RLPs, which are involved in development- and immune-signalling, respectively. eJM-TM-cJM region of LRR-RLPs evolved to allow interactions with SOBIR1 to induce immunity (negatively charged eJM and GxxxG). LRR-RLK-Xbs utilize Xb kinase domains to induce distinct downstream responses.

Our work has uncovered multiple evolutionary mechanisms underlying PTI: i) expansion of the number of receptors and their recognition specificity; ii) development of an increasingly complex immune network; iii) adaptation of existing receptors for specific signalling (Main figure 6b). i) Many PRRs are lineage-specific, with the characterised LRR-RLK-XII and LRR-RLPs found only in specific lineages within angiosperms, such as the Brassicaceae (EFR) and Solanaceae (Cf2/4/5/9). This expansion of PRR subgroups, including LRR-RLKs, LRR-RLPs, G-lectin-RLKs, and WAK-RLKs, has enabled plants from different lineages to recognise a broader range of molecules specific to certain pathogens, exemplified by Xa21-Raxx21 and Cf2/4/5/9-Avr2/4/5/9 interactions^69–73^. ii) Cell-surface co-receptors, such as SERKs and SOBIR1 emerged during or around the time of land plant evolution. Moreover, several cell-surface receptors involved in signalling regulation, such as malectin-RLKs (FERONIA)^74^ and LRR-RLK-Xa (BIR1, BIR2)^75–78^ are found exclusively in Embryophytes. PRRs differentially activate downstream signalling components, including RLCK-VIIs, CDPKs and EP proteins, which fine-tune the magnitude of immune responses^10,79^. RLCK-VIIs are expanded in the Embryophytes, while EP proteins and RPW8-NLRs are specific to Tracheophytes. Collectively it seems that increasingly intricate and specialised PTI-signalling networks enhance the flexibility and regulation of immune responses to keep up with the changes taking place in rapidly evolving pathogens^80^. iii) The structural similarities between LRR-RLPs and LRR-RLK-Xbs implies a common origin between immune-specific cell-surface receptors and development-specific cell-surface receptors. The exact nature of the common ancestral form of these receptors, whether an RLK or an RLP, remains, and perhaps will always remain uncertain. Both LRR-RLK-Xbs and LRR-RLPs with ID+4LRR can be found in land plants, so it is conceivable that LRR-RLK-Xb, with its kinase domain predating land plant emergence, evolved from an integration of an LRR-RLP containing an ID+LRR into an Xb kinase domain^81^. In this scenario, the common ancestral receptors could have recognised a PAMP, with the peptide sequence of this PAMP possibly converted to serve as a phytocytokine to regulate plant developmental processes. Multiple phytocytokines, such as PSK, PSY, SCOOPs and CLE peptides, are present in plant pathogens and pests^82–86^. Whether the perception of phytocytokines evolved from the perception of PAMPs, or pathogens developed phytocytokine-mimics to repress immune responses remains an open question.

## Methods

### RLK, RLP, and ectodomain identification

LRR-RLK, LRR-RLP, LysM-RLKs and Nb-ARC proteins were identified as described previously^8^. For the other RLK and RLPs described in this study, only primary gene model proteins longer than 150 AA were used^8^. Based on the presence of a kinase domain (KD) and/or a trans-membrane domain (TM), the proteins fell into three major groups: 1) RLKs with KD and TM, 2) RLPs without KD but with TM, and 3) ectodomains without KD or TM.

To identify RLK candidates, we first searched for the presence of a protein kinase domain (PFAM: PF00069.26) with hmmer (version 3.1b2, option -E 1e-10^87^). If multiple hits were found, only the best match was kept. Potential signalling peptides were removed with SignalP (version 5.0b^88^) to avoid identifying and including signal peptides as TMs. TMs were searched with tmhmm (version 2.0^89^). The location of the KD and the TM were used to split the protein sequence into the endodomain (with KD) and ectodomain (without KD). Given that LRR-RLKs were previously identified ^8^, we searched and removed proteins with LRR repeats in their ectodomains with predict-phytolrr^90^ (obtained in December 2021).

To identify RLPs and candidate ectodomains (without KD or TM), we first removed all proteins with a kinase domain match (hmmer with the option -E1e1000) and removed potential signalling peptides to avoid their inclusion with TMs. We then searched for TMs with tmhmm and kept proteins with no TM (ectodomains) or 1-2 TM (RLPs). Proteins containing LRR repeats were identified with predict-phytolrr and removed from the RLP/ectodomain set.

Finally, we searched candidates from all three major groups for the presence of Duf26 (PFAM: PF01657.18), malectin or malectin-like (PFAM: PF11721.9 and PF12819.8), G-type lectin (PFAM::PF01453.25), L-type lectin (PFAM: PF00139.20), and WAK domains: EGF-like (PFAM: PF00008.28), Calcium-binding EGF (PFAM: PF07645.16), and Wall-associated receptor kinase galacturonan-binding (PFAM: PF13947.7) with hmmer (option -E 10). Hits from all searches were combined for each group, and proteins were assigned to the hit with the highest score.

### Identification of signalling components

Signalling components were identified using different approaches, but always using primary gene model proteins longer than 150 AA^8^. 150AA cut-off was used to eliminate truncated proteins.

CNGCs, OSCAs, and RBOHs were identified by hmmer searches (option -E 10) for specific domains. Ion transport protein domains (PFAM: PF00520.32) were used for CNGCs, PHM7_cyt (PFAM: PF14703.7) and RSN1_TM (PFAM: PF13967.7) were used for OSCAs, and FAD-binding domains (PFAM: PF08022.13), ferric reductase like transmembrane components (PFAM: PF01794.20), and ferric reductase NAD binding domains (PFAM: PF08030.13) were used for RBOHs.

EP proteins (EDS1, PAD4, and SAG101) were identified by hmmer searches (option -E 10) for the Lipase (class 3) domain (PFAM: PF01764.26) and the Enhanced disease susceptibility 1 protein EP domain (PFAM: PF18117.2). To further classify the candidates among the known EDS1, PAD4, and SAG101 candidates, we clustered all candidates with MMSeq2 (Release 14-7e284, options --min-seq-id 0.3 -c 0.3^91^). All known EDS1 (*AT3G48090*, *Niben101Scf06720g01024.1*, and *Solyc06g071280*), PAD4 (*AT3G52430*, *Niben101Scf02544g01012.1*, and *Solyc02g032850*), and SAG101 (*AT5G14930*,

*Niben101Scf00271g02011.1*, *Niben101Scf01300g01009.1*, *Solyc02g069400*, and *Solyc02g067660*) proteins were found in exactly one cluster each. Hence, we used the three matching clusters as EDS1, PAD4, and SAG101 proteins.

RPW8-NLRs (NRG1 and ADR1) were identified similarly using the NB-ARC (PFAM: PF00931.23) and RPW8 (PFAM: PF05659.12) domains. After clustering as described above, we found all known NRG1 and ADR1 sequences in one single cluster. This allowed us to extract and re-cluster these sequences with more stringent parameters (options --min-seq-id 0.3 -c 0.75). After that, we found all known NRG1 (*AT5G66900*, *AT5G66910*, and *Niben101Scf02118g00018.1*) and ADR1 (*AT1G33560*, *AT4G33300*, *AT5G04720*, and *Niben101Scf02422g02015*) proteins in exactly one cluster each, indicating that the two matching clusters could be considered as NRG1 and ADR1 proteins.

The remaining signalling component candidates (SOBIR1, RLCK-VII, CDPK, MAPK, MAPKK, and MAPKKK) were identified using previously published HMM profiles^92^ using hmmer (option -E 10). The following patterns were used for the families of interest: SOBIR1: RLK-Pelle_LRR-XI-2, RLCK-VII: RLK-Pelle_RLCK-VIIa-1, RLK-Pelle_RLCK-VIIa-2, RLK-Pelle_RLCK-VIIb, CDPK: CAMK_CDPK, MAPK: CMGC_MAPK, MAPKK: STE_STE7, and MAPKKK: STE_STE11.

### Expansion rate of cell-surface receptors and signalling components

The percentages (%) of cell-surface receptors and signalling components from each genome was calculated as (number of identified genes/number of searched genes × 100). Next, the percentages from each species within a lineage (e.g, Rhodophtya or green algae) were grouped and the median percentage was calculated. Median value was used instead of mean to avoid outliers within the lineages. The expansion rate within a species is calculated by ((% cell surface receptors or signalling components in that species)-(median))/(median). For example, the expansion rate of LRR-RLP family in *Marchantia polymorpha* from green algae is calculated by ((%LRR-RLP in *Marchantia polymorpha*)-(median %LRR-RLP in green algae)/(median %LRR-RLP in green algae). Values larger than 0 indicate expansion; values equal to 0 indicate no expansion, and values below 0 indicate contraction. Note that the reliability of the expansion rate is dependent on the number of species used to calculate the median, which is also dependent on the available genomes in Glaucophyta, red algae (Rhodophyta), green algae, and Bryophytes.

### Identification of N-loopouts (NLs) island domains (IDs)

To identify N-loopouts (NLs) or island domains (IDs) in LRR-RLK and LRR-RLP proteins, we used a dataset of previously described LRR-RLKs and LRR-RLPs^8^. We searched the LRR-RLKs again for kinase domains (PFAM: PF00069.26) with hmmer (option -E 1e-10), and kept only the best match for each protein. LRR-motifs and transmembrane domains were searched in both groups with predict-phytolrr^90^ and tmhmm^89^, respectively. LRR-RLKs were filtered for the presence of internal KD motifs, one or two TM, and at least two external LRR repeats (‘internal’ was defined as the side with the kinase domain). LRR-RLPs were filtered for the absence of a KD and the presence of one or two TM and at least two external LRR-motifs as defined by the site with more LRR repeats). The outer LRR-motifs were then used to identify NLs and IDs: Individual repeats were grouped into LRR-regions if they were less than 13 AA apart from each other. Gaps between LRR-regions or LRR-motifs that were 15-29 AA or 30-90 AA long were extracted as NL and ID candidates, respectively. After extracting gap sequences, all sequences were again checked for the presence of LRR-motifs using predict-phytolrr and hmmer using all LRR patterns as described previously^8^, and LRRsearch^93^. Only NLs and IDs without any LRR match were included in the final dataset. Locations of the NLs and IDs in relation to LRR-motifs and LRR-regions were determined with a custom R-script. ID sequences were aligned to each other with FAMSA^94^ without trimming^95^ and phylogenetic trees were inferred with FastTree^96^ (version 2.1.11 SSE3, option -wag). Trees were rooted with gotree^97^ (v0.4.2) using one sequence belonging to the most basal species as outgroup, according to the taxonomic tree.

The phylogenetic tree of the IDs was used to cluster the proteins: the tree was converted into a distance matrix using the function cophenetic.phylo() from the R-package ‘ape’^98^ (version 5.6-2). Distances smaller than 0.2 (*i.e.* less than 0.2 substitutions per site on average) were extracted, converted to similarities, and used as edges in a network. Communities within this similarity network were identified with the function cluster_louvain^99^ implemented in the R-package ‘igraph’^100^ (version 1.2.6).

### In-depth phylogeny of the ectodomains from LRR-RLPs and LRR-RLKs

In-depth analysis of the ectodomains from LRR-RLPs and LRR-RLKs was done using the LRR-RLKs and LRR-RLPs from the NLs and IDs search (see above). We first searched for the C3 domain in each sequence^49,8^ with hmmer and selected the best hit. We then pruned the sequences to include everything from the C3 domain to the C terminus. For the LRR-RLKs, we further searched sequences for kinase domains and removed sequences upstream of the start of the kinase domain. That is, for all LRR-RLKs and LRR-RLPs, we extracted regions with C3, eJM, TM, or eJM domains. These sequences were aligned with FAMSA. Specific domains (*e.g*. C3 or eJM) were subsequently extracted from this alignment. After extraction, the phylogenetic trees of specific domains were constructed as described above (FAMSA, FastTree, gotree).

### Taxonomic trees

Taxonomic trees used in this study were identical to the ones described previously^8^: The taxonomic tree for visualizing the entire data set and selecting outgroups was obtained from NCBI (https://www.ncbi.nlm.nih.gov/Taxonomy/CommonTree/wwwcmt.cgi). The tree used for testing the relationship between the fraction of candidates found and phylogenetic distances, was obtained from a previous report^101^. The latter contained 238 out of the 351 genomes analysed. Phylogenetic trees were visualised and pruned, and figures were generated with iTOL^102^.

### Test for similarities in fraction of proteins and phylogenetic relationships

Tests for similarities in fraction of proteins and phylogenetic relationship were done as described previously^8^: To test whether the fraction of certain proteins found per species correlated with predicted phylogenetic relationships, we converted the fractions and the phylogenetic tree to distance matrices and tested for correlation with mantel tests (R-package vegan, version 2.5-7 with 10’000 permutations). Analogously, we also tested for correlation between distance matrices obtained for two different sets of proteins. P-values were corrected for multiple testing to reflect false discovery rates (FDRs^103^).

## Vector construction

The CDS regions of AtRLP23, AtBRI1, AtPSKR2, AtPSY1R, AtEFR, and AtBES1 were amplified by PCR with KoD one (Toyobo, Japan), and the PCR products were cloned into the epiGreenB5 (3×HA) vector between the *Cla*I and *Bam*HI restriction sites with In-Fusion HD Cloning Kit (Clontech, USA) to generate p35S::BES1-HA or p35S::cell-surface receptor-HA (epiGreenB5-Cauliflower mosaic virus (CaMV) p35S:gene of interest-3×HA). The constructs were then transformed into *Agrobacterium tumefaciens* strain AGL1 for transient expression in *Nicotiana benthamiana*. All chimeric cell-surface receptors generated in this study contain the EFR signal peptide to ensure consistency between the constructs and expression levels.

### Transient expression in *Nicotiana benthamiana*

*Agrobacterium tumefaciens* strain AGL1 carrying the binary expression vectors described above were grown on LB agar plates amended with selection antibiotics. Cultures were pelleted, centrifugated, and then resuspended in infiltration buffer (10mM MgCl_2_, 10mM MES at pH5.6, and 100μM acetosyringone). The concentration of AGL1 was then adjusted to OD_600_ = 0.5 and syringe-infiltrated into *Nicotiana benthamiana* leaves.

### Protein extraction and immunoprecipitation

Protein extraction for immunoprecipitation was performed as previously described^104^. Three days after transient expression, three to four grams of *N. benthamiana* leaves were treated with elicitors and snap-frozen. The tissues were then ground in liquid nitrogen and extracted in extraction buffer (50mM Tris-HCl at pH 7.5, 150mM NaCl, 10% glycerol, 5mM DTT, 2.5mM NaF, 1mM Na_2_MoO_4_•2H2O, 0.5% polyvinylpyrrolidone (w/v), 1% Protease Inhibitor Cocktail (P9599; Sigma-Aldrich), 100μM phenylmethylsulphonyl fluoride and 2% IGEPAL CA-630 (v/v; Sigma-Aldrich), and 2mM EDTA) at a concentration of 3mL/g tissue powder. Samples were then incubated at 4 °C for an hour and debris was removed by centrifugation at 13,000rpm for 10min at 4 °C. Supernatants were collected, protein concentrations were adjusted to 5mg/mL, then incubated with rotation for an hour at 4 °C with 50μL anti-HA magnetic beads (Miltenyi Biotec) for immunoprecipitation. Magnetic beads were then washed twice with extraction buffer and the HA-tagged protein was eluted with sodium dodecyl sulphate (SDS) sample buffer at 95 °C.

### Immunoblotting

Protein extractions were performed as previously described^16^. *N. benthamiana* leaves were infiltrated with elicitors and snap-frozen at indicated time points. The tissues were then lysed in liquid nitrogen and extracted in 1×NuPAGE™ LDS Sample Buffer (Invitrogen™) with 10mM DTT at 70 °C for 10 minutes. Total proteins were then separated by SDS-PAGE and blotted onto a nitrocellulose membrane (Trans-Blot Turbo Transfer System, Bio-Rad). The membrane was then blocked in a solution of either 5% skimmed milk (for BES1 and cell-surface receptor detection) or 5% bovine serum albumin (BSA; for MAPK detection) in Tris-buffered saline, 0.1% Tween 20 detergent (TBST) for an hour. Phosphorylated MAPKs were detected using α-phospho-p44/42 MAPK rabbit monoclonal antibody (D13.14.4E, in 1:2000, Cell Signalling Technology, USA) in a solution of 5% BSA in TBST overnight at 4 °C. HA-tagged BES1 or cell surface receptors were detected using Anti-HA-Peroxidase, High Affinity, rat IgG_1_ antibody (Roche) in a solution of 5% skimmed milk in TBST overnight at 4 °C. For detection of MAPKs, this was followed by incubation with α-rabbit IgG-HRP-conjugated secondary antibodies (1:10000, Roche, USA) in a solution of 5% BSA in TBST for an hour at room temperature. HRP signal was then detected by Clarity Western ECL Substrate (Bio-Rad) with a LAS 4000 system (GE Healthcare, USA). Nitrocellulose membranes were stained with Coomassie Brilliant Blue (CBB) to ensure equal loading.

### ROS assays

ROS burst assays were performed as described previously^104^. *N. benthamiana* leaf discs were collected with a 4-mm-diameter cork borer and placed in 96-well plates with 120μl deionised water overnight in the dark (abaxial surface of the leaves facing down). *N. benthamiana* leaf discs were then treated with either mock (water) or 1 μM nlp20 in 20 mM luminol (Wako, Japan) and 0.02 mg ml^−1^ horseradish peroxidase (Sigma-Aldrich). Luminescence was then measured over indicated periods of time with a Tristar2 multimode reader (Berthold Technologies, Germany).

## Contributions

B.P.M.N., Y.K. and K.S. conceived and conceptualised the study; B.P.M.N., Y.K., M.W.S. and M.W. designed the bioinformatic analyses; M.W.S. and M.W. performed the bioinformatic analyses; B.P.M.N. performed the vector construction with assistance from Y.K.. B.P.M.N. performed protein extractions, immunoprecipitation, immunoblotting and ROS assays; B.P.M.N. wrote the original draft; and B.P.M.N., M.W.S., M.W., Y.K., an K.S. reviewed and edited the manuscript.

## Acknowledgments

We thank Yoko Nagai, Naomi Watanabe, and Mizuki Yamamoto for providing technical support, Cyril Zipfel, Simon Snoeck and Jonathan Jones for discussions and critical reading of the manuscript. We also thank Markus Albert for sharing the information of chimeric receptors and BES1-HA experiments. B.P.M.N. is an International Research Fellow of Japan Society for the Promotion of Science (Postdoctoral Fellowships for Research in Japan (Standard), 21F21793). The research was financially supported by MEXT/JSPS KAKENHI Grant Numbers, 20H02994, 21K19128, JPJSBP1-20193222 (to Y.K.), 20H05909 and 22H00364 (to K.S.).

---

**Extended figure 1.**
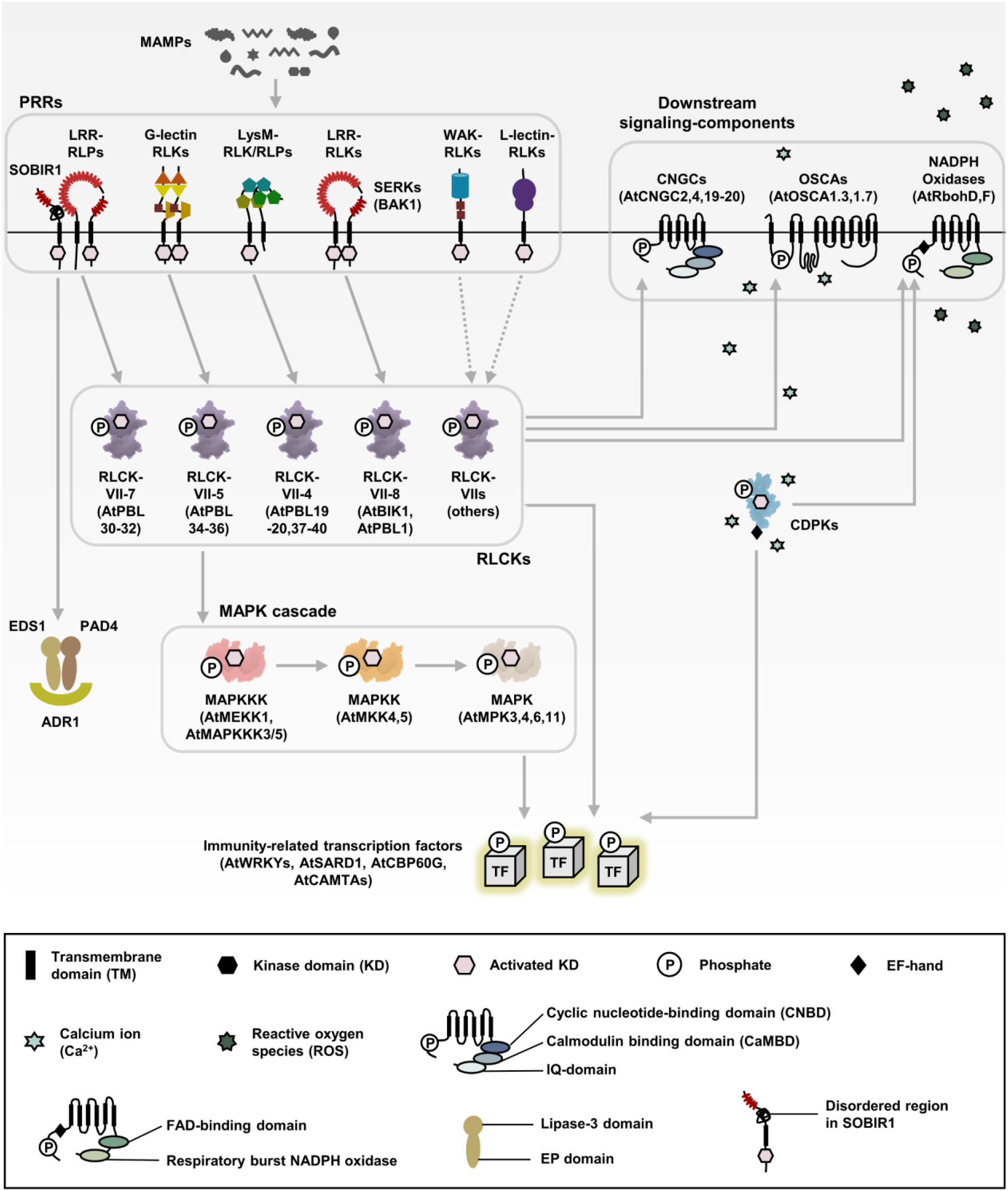
PTI signalling pathway in plants. Schematic showing the PTI signalling pathway in plants. The architecture of different signalling components is defined in the lower box. Grey arrow represents the activation of downstream signalling components by upstream signalling components following the perception of PAMPs/MAMPs by PRRs. Multiple signalling components are activated through phosphorylation by an activated kinase domain (KD). Downstream signalling components are activated to produce ROS and cytosolic calcium influx. Multiple immune-related transcription factors (TF) are also activated by cytoplasmic protein kinases, which then lead to defence-related gene expression.

**Extended figure 2.**
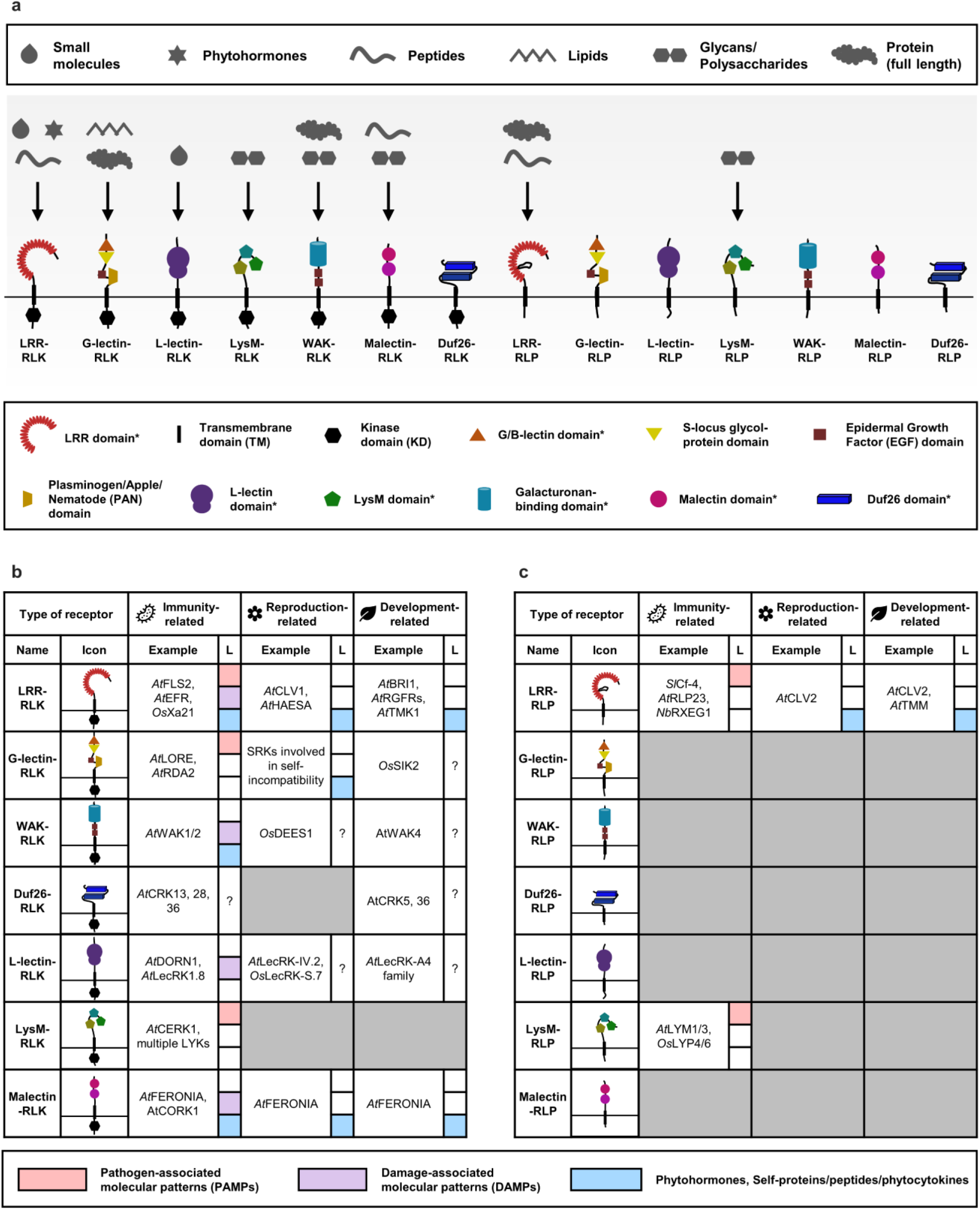
Domain architecture and roles of the major plant cell-surface receptor classes. **(a)** Schematic displays the domain architecture of different classes of receptor-like kinase (RLKs) and receptor-like proteins (RLP) in plants. Arrows represent the ligands which these receptor classes have been reported to perceive or recognize. The upper box defines the ligands recognized by different receptors. The lower box defines the domains within the receptor classes. ‘**\*’** represents the definitive domains in these receptors. Note that these receptors may be able to recognise other unidentified ligands. The number of definitive domains and the domain architecture in each receptor are variable. **(b-c)** Characterised (b) RLKs and (c) RLPs in plants; the biological processes in which the characterised members are involved, and type of ligand (L) they perceive. A grey box indicates that no receptor class has been reported to be involved in that biological process. For ligands, a red box indicates PAMPs from pathogens, a purple box indicates DAMPs released from damage, and a blue box indicates phytohormones, self-protein/peptide or phytocytokines. A question mark (?) indicates an unidentified ligand. Abbreviations for plant species: *A. thaliana*, *At*; *S. lycopersicum*, *Sl*; *O. sativa*, *Os*; *N. benthamiana*, *Nb.* References to the genes are included in the supplementary information.

**Extended figure 3.**
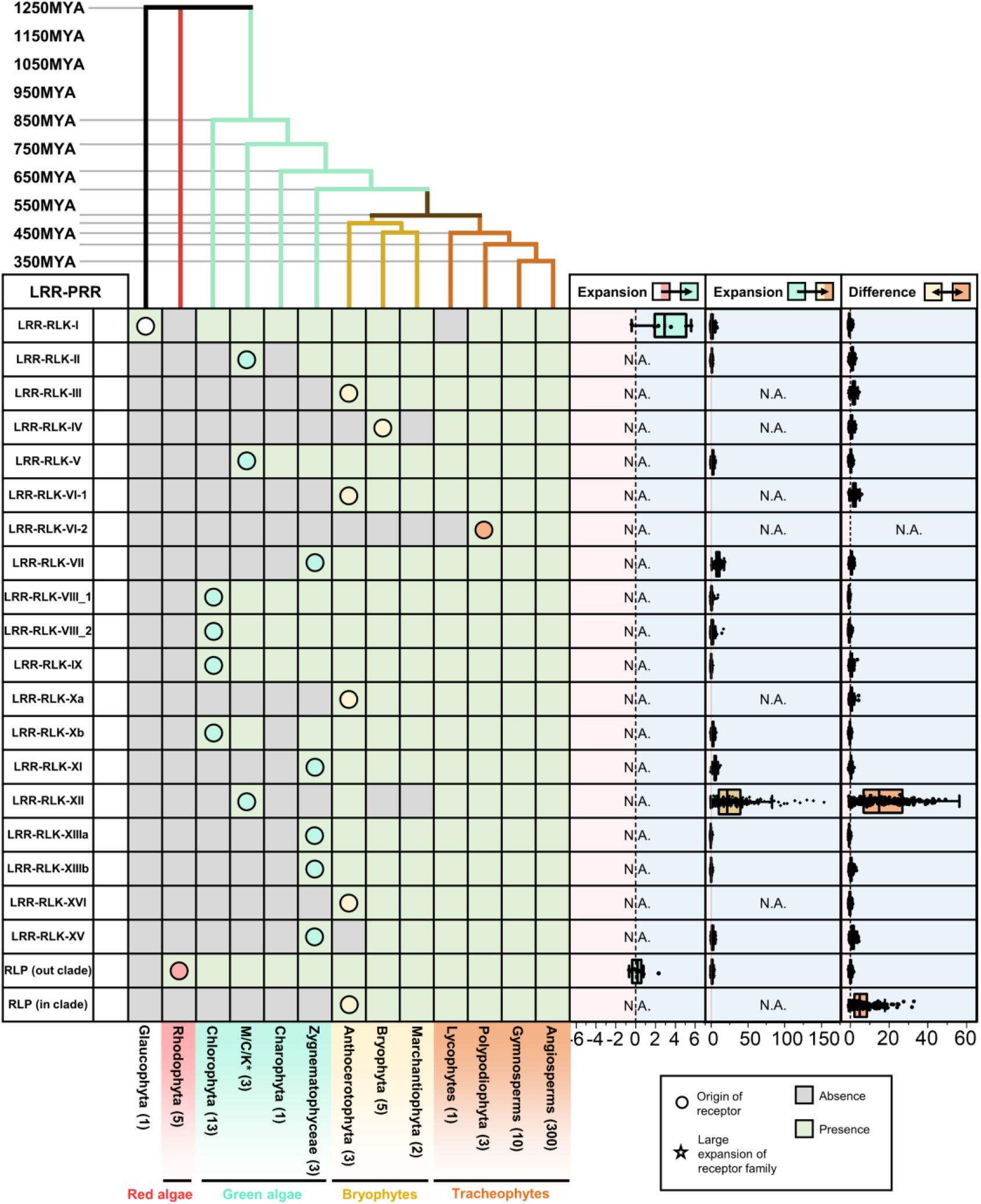
The origin and expansion of LRR-containing cell-surface receptors in plants. Top panel represents the phylogenetic tree of multiple algal and plant lineages. The timescale (in million years; MYA) of the phylogenetic tree was estimated by TIMETREE5^23^. Bottom panel represents the presence or absence of different receptor classes in algal and plant lineages. RLP represents LRR-RLPs. ‘Out clade’ refers to LRR-RLPs outside the ID+4LRR clade in Figure 4c and ‘in clade’ refers to LRR-RLPs inside the ID+4LRR clade in Figure 4c. *M/C/K represents Mesostigmatophyceae, Chlorokybophyceae, and Klebsormidiophyceae. The number of available species from each algal and plant lineage is indicated by numbers in the boxes. A grey box indicates the absence of receptor, and a green box indicates the presence of receptors in each lineage. Receptor origin is indicated with a circle (○). Expansion rates of receptor classes are indicated by boxplots. The cyan boxplot represents the expansion rate from Glaucophyta and Rhodophyta to green algae. The yellow boxplot represents the expansion rate from green algae to embryophytes and the orange boxplot represents the differences between Bryophytes and Tracheophytes. Light blue area represents expansion and light pink area represents contraction of the gene family. X-axis values represent expansion rate (×). Boxplot elements: center line, median; bounds of box, 25th and 75th percentiles; whiskers, 1.5 × IQR from 25th and 75th percentiles.

**Extended figure 4.**
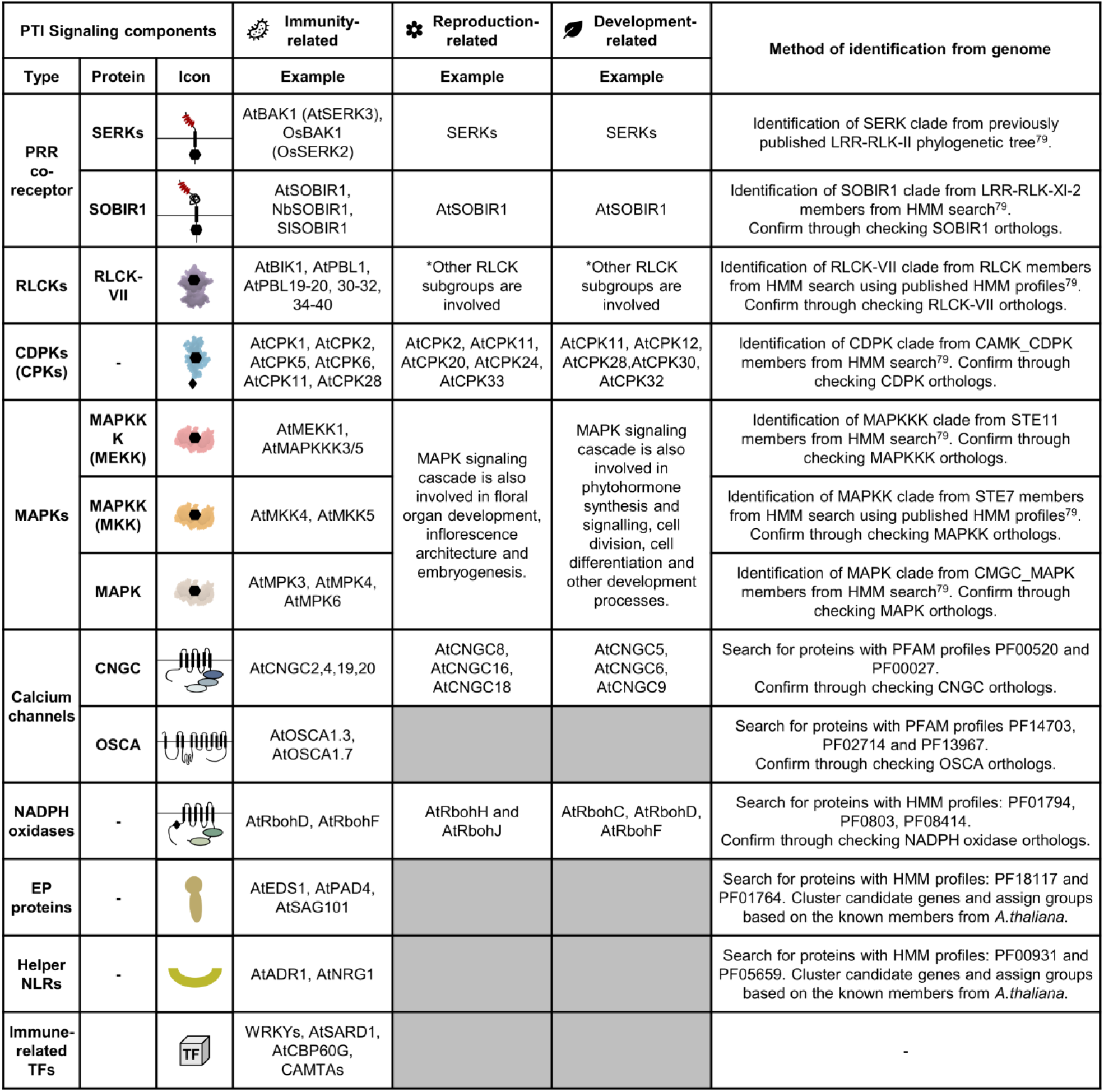
Roles of signalling components in different biological processes in plants. Characterised PTI signalling components in plants and the biological processes they are involved in. Characterised examples of each signalling component involved in different biological processes are given. A grey box indicates that no signalling component has been reported to be involved in that biological process. The methods of identifying these signalling components from genomes are also given. Abbreviations for plant species: *A. thaliana*, *At*; *S. lycopersicum*, *Sl*; *N. benthamiana*, *Nb.* References to the genes are included in Supplementary information.

**Extended figure 5.**
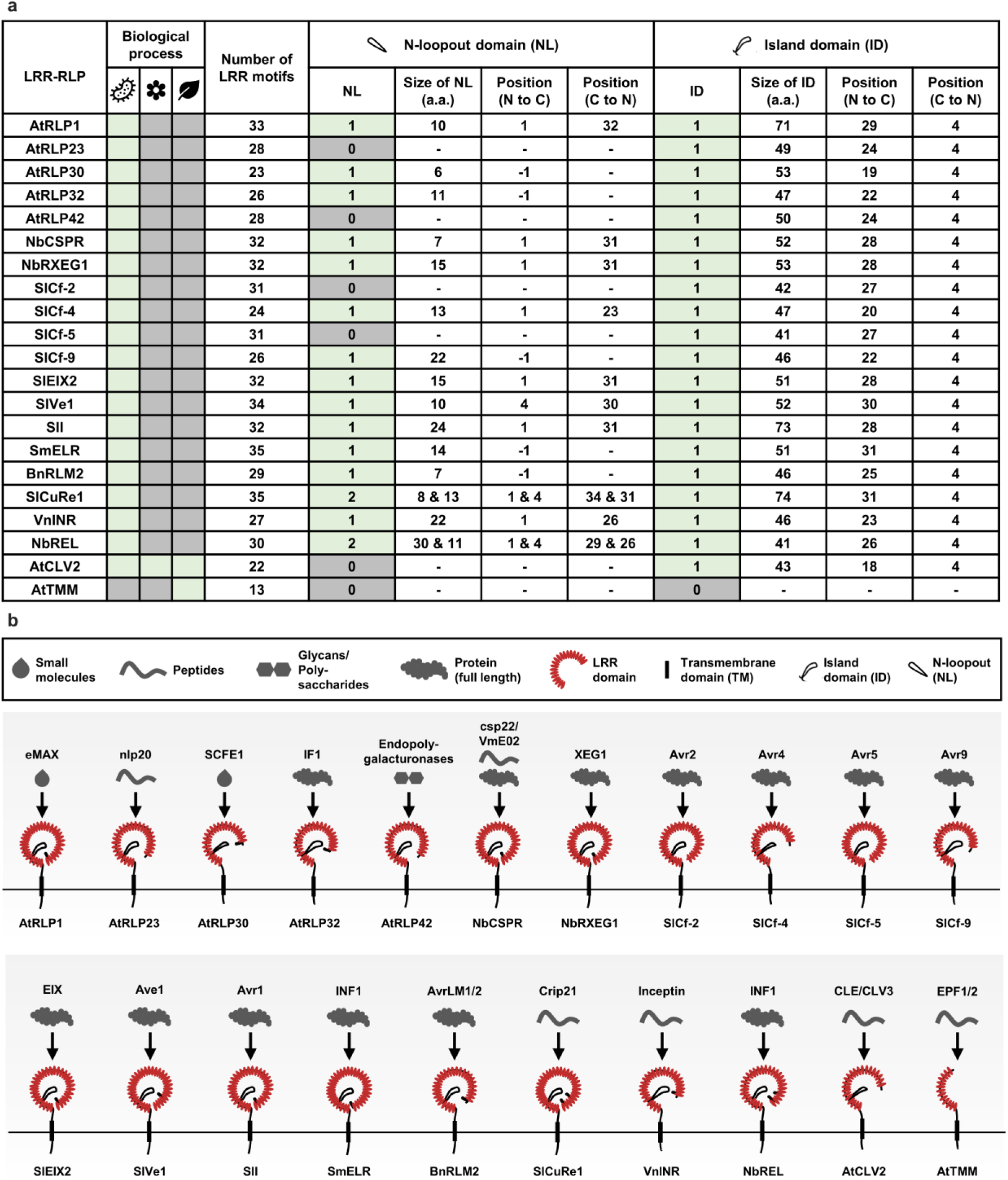
Domain architecture of the characterized LRR-RLP in plants. **(a)** Domain architecture of the characterised LRR-RLPs in plants. Characterised receptors involved in microbial interactions (bacteria icon), reproduction (flower icon) and development (leaf icon) are indicated with green boxes. A grey box indicates that no receptor class has been reported to be involved in that biological process. The numbers of LRR motifs, N-loopouts (NLs) and island domains (IDs), and the position of NL/ID (from N-to-C terminal or C-to-N terminal) are obtained from structures shown in supplementary figure 28. For NL, -1 position from N-to-C terminal indicates that the NL is positioned before the LRR motifs in the ectodomain. **(b)** Domain architecture schematic of the characterised LRR-RLPs in plants. Arrows represent the ligands of each reported receptor class. Upper box defines ligands recognised by different LRR-RLPs and the domains in the LRR-RLPs. Number of LRRs and position of ID/NL were obtained from structures shown in supplementary figure 28. Note that these receptors may be able to recognise additional unidentified ligands. References to the genes are included in Supplementary information.

**Extended figure 6.**
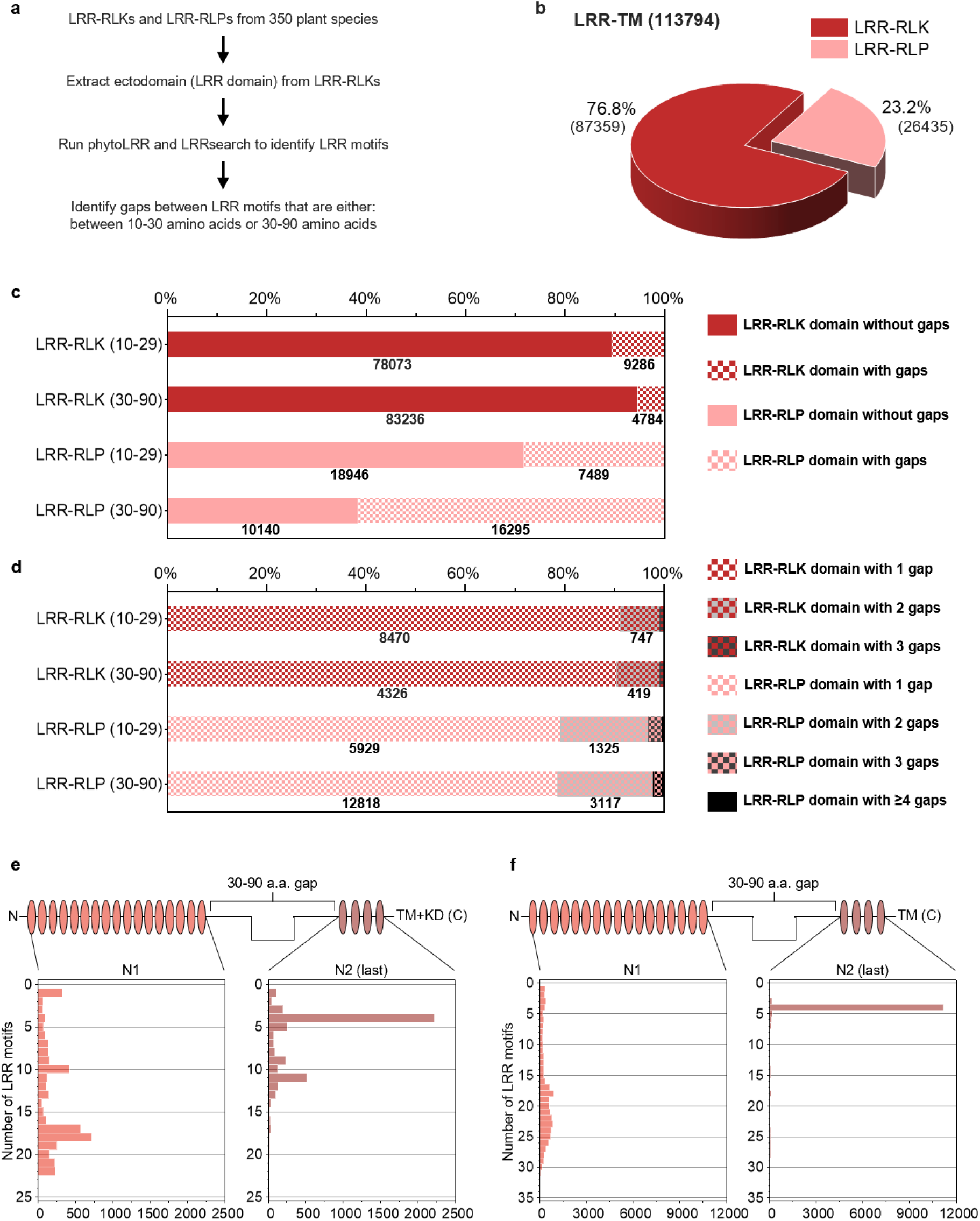
Distribution of gaps within LRR motifs in LRR-containing cell-surface receptors. **(a)** Method of gap identification from LRR-containing cell-surface receptors. **(b)** Distribution of LRR-RLKs and LRR-RLPs in LRR-containing cell surface receptors (LRR-TM) from 350 plant species. **(c)** Distribution of LRR-RLPs and LRR-RLKs with or without gaps ranging from 10-29 AA (10-29) or 30-90 (30-90). AA Labels are indicated on the left. **(d)** Distribution of LRR-RLPs and LRR-RLKs with one, two, three, or >3 gaps of 10-29 AA (10-29) or 30-90 (30-90). AA labels are indicated on the left. **(e-f)** Position of large gaps (IDs; 30-90 AA) in **(e)** LRR-RLKs and **(f)** LRR-RLPs with a single large ID. N1 represents the number of LRR motifs before the IDs and N2 represents the number of LRR motifs after the ID. For positions of gaps in LRR-RLKs and LRR-RLPs with multiple gaps, refer to supplementary figure 29-32.

**Extended figure 7.**
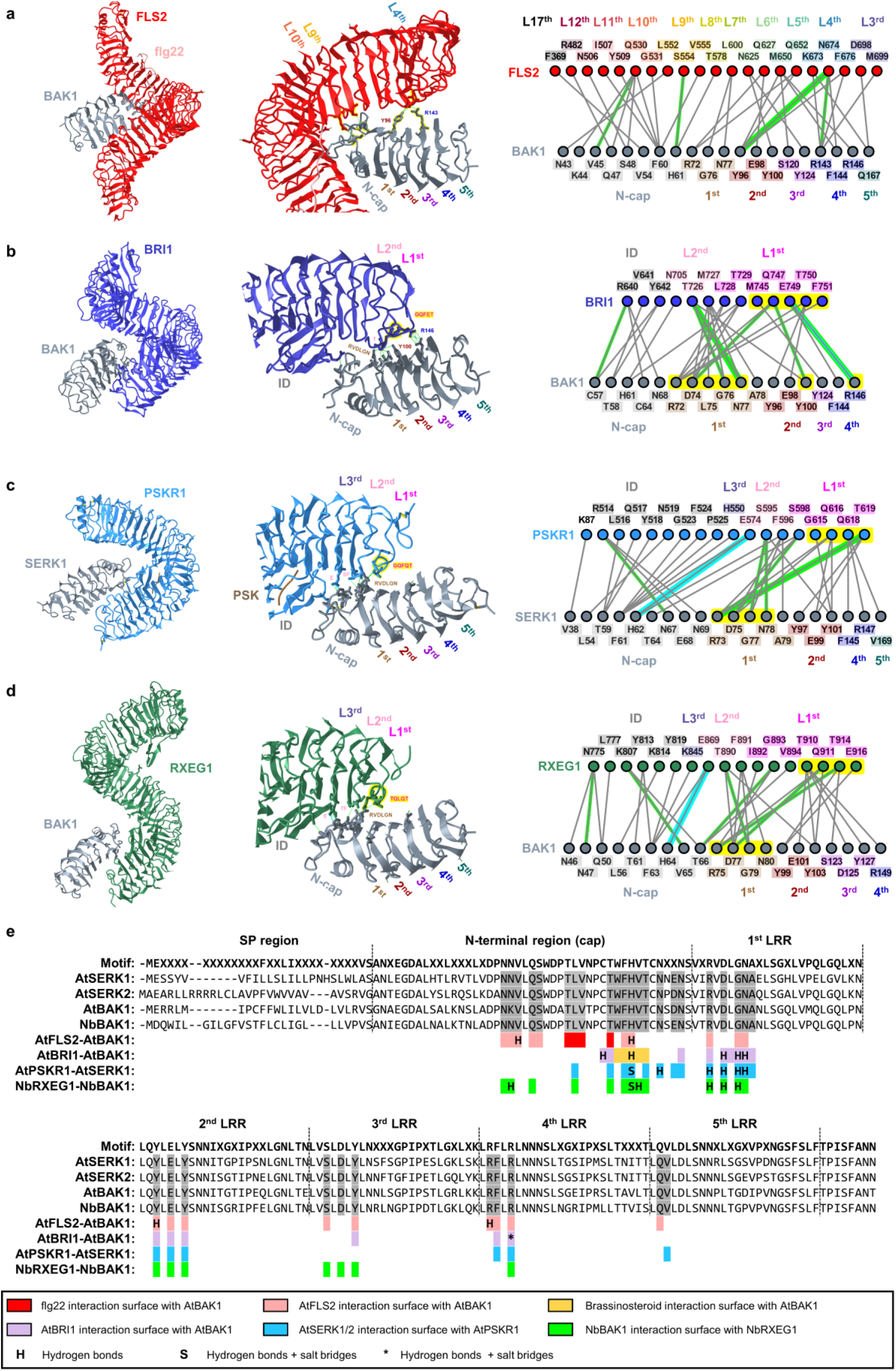
Interaction surfaces between SERKs, LRR-RLKs and LRR-RLP. (a-d) Structures and interaction interfaces of LRR-RLKs and LRR-RLPs with SERKs. Published structures of **(b)** AtFLS2-AtBAK1^18^, **(c)** AtBRI1-AtBAK1^56^, **(d)** AtPSKR1-AtSERK1^51^, and **(e)** NbRXEG1-NbBAK1^57^ are shown. The left panels show the full structure, and the middle panels show the interaction sites between LRR-RLKs or LRR-RLP and SERKs. Hydrogen bonds are indicated by green dotted lines, and salt bridges are shown as cyan dotted lines. The positions of LRR residues (counting from N to C for SERKs and counting from C to N for LRR-RLKs and LRR-RLP) are shown. Amino acid residues that are important for the interactions are labelled and the QxxT motifs are highlighted in yellow (red text). The right panel represents the 2D interaction network between SERKs and its receptors. Contacts/interactions are shown in grey lines, hydrogen bonds are shown in green lines, and salt bridges are shown in cyan lines. Amino acids are labelled in colours according to their positions in the LRR motifs (counting from N to C for SERKs and counting from C to N for LRR-RLKs and LRR-RLP (L)). Residues around and within the QxxT motifs in BRI1, PSKR1, and RXEG1 are highlighted in yellow. Residues in SERKs that are involved in the interactions with QxxT motifs are also highlighted in yellow. Structures were visualized in iCn3D^21^. For **(a-e)**, the interaction sites are calculated by iCn3D with the following thresholds: hydrogen bonds: 4.2Å; salt bridges/ionic bonds: 6Å; contacts/interactions: 4Å. **(e)** Alignment of the ectodomains of SERKs from *A. thaliana* (At) and *N. benthamiana* (Nb). The amino acid residues between the interaction interfaces of AtFLS2-AtBAK1, AtBRI1-AtBAK1, AtPSKR1-AtSERK1, and NbRXEG1-NbBAK1 are highlighted in different colours as indicated in the boxes below. Hydrogen bonds and salt bridges/ionic bonds are also indicated.

**Extended figure 8.**
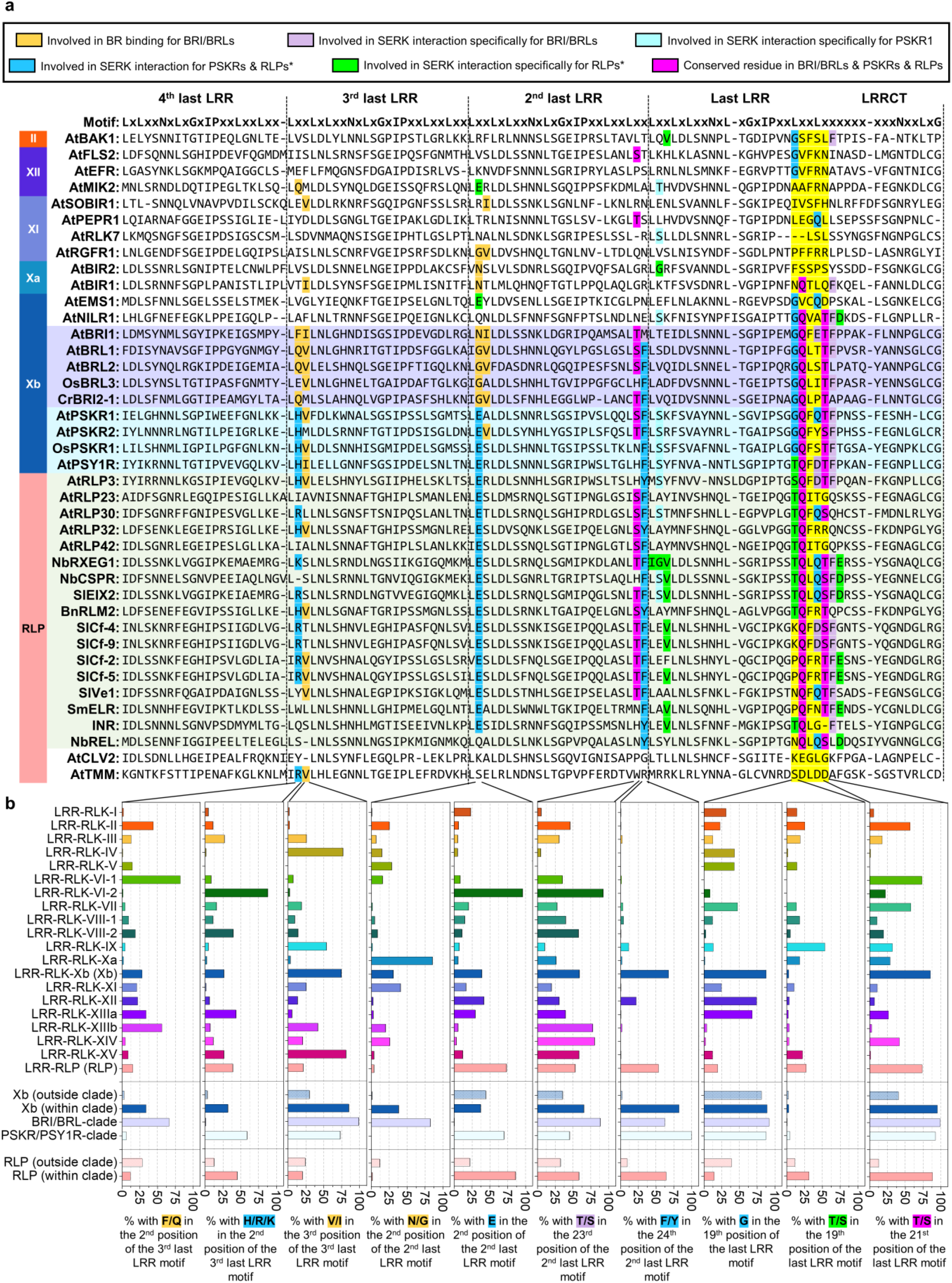
Alignment and features of the terminal four LRRs (C3) in LRR-RLKs and LRR-RLPs. **(a)** Alignment of the C3 region in LRR-RLKs (from subgroups II, XII, XI, Xa, Xb) and LRR-RLPs. The BRI1/BRL clade is highlighted in purple; PSKR/PSY1R clade is highlighted in cyan and LRR-RLP with the ID+4LRR clade (see Figure 4c) is highlighted in light green. Amino acid residues involved in brassinosteroid (BR) binding for BRI/BRL and residues required for SERK interaction for BRI1/BRL, PSKR1 and RXEG1 (LRR-RLP) are highlighted. The colour code for each highlight is indicated in the box on top. The interaction sites were calculated using iCn3D^21^ with the following thresholds: hydrogen bonds: 4.2Å; salt bridges/ionic bonds: 6Å; contacts/interactions: 4Å. For details, please refer to Extended Figure 7. **(b)** Percentages of LRR-RLKs or LRR-RLPs with the stated amino acid residues in the corresponding position in **(a)**. Percentages (%) were calculated by the number of LRR-RLKs or LRR-RLPs in the subgroup with the stated residue divided by the number of LRR-RLKs or LRR-RLPs in the subgroup without the stated residue × 100. RLP/RLK-Xb (outside clade) refers to receptors outside the light grey clade in Figure 4c. RLP/RLK-Xb (within clade) refers to receptors inside the light grey clade in Figure 4c. Number of cell-surface receptors (n) in each LRR-RLK subgroup: I, n = 752; II, n = 682; III, n = 6572; IV, n = 1033; V,n = 8; VI-1, n = 84; V1-2, n = 146; VII, n = 1720; VIII-1, n = 195; VIII-2, n = 411; IX, n = 70; Xa, n = 96; Xb, n = 3182; XI, n = 8807; XII, n = 12863; XIIIa, n = 739; XIIIb, n = 465; XIV, n = 241; XV, n = 548; Xb (outside clade), n = 580; Xb (within clade), n = 2527; BRI/BRL clade, n = 1170; PSKR/PSY1R clade, n = 1347. Number of cell-surface receptors (n) in each LRR-RLP subgroup: LRR-RLP (RLP), n = 24970; RLP (outside clade), n = 5000; RLP (within clade), n = 19970.

**Extended figure 9.**
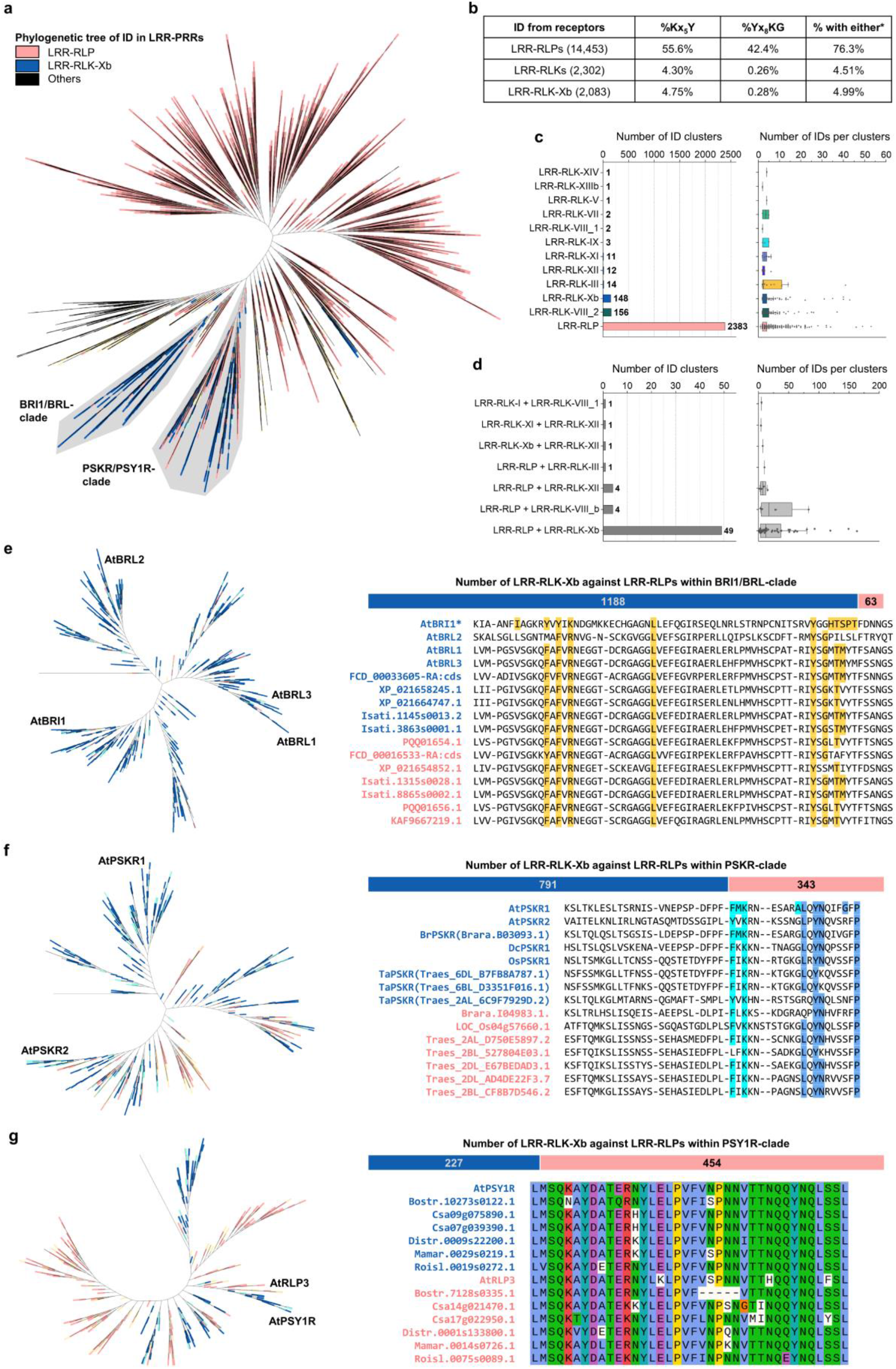
Ectodomains shared by LRR-RLK-Xbs and LRR-RLPs. **(a)** Phylogenetic tree of IDs of all LRR-containing PRRs from 350 species. Branches are labelled in colours as indicated. **(b)** Percentage (%) of IDs (before the last four LRR motifs) from LRR-RLP, LRR-RLK, and LRR-RLK-Xb with the Kx_5_Y, Yx_8_KG, or either Kx5Y or Yx8KG (*) motifs. **(c)** Number of ID clusters with LRR-receptors from a single subgroup/family. The left graph shows the number of clusters containing LRR-receptors from each subgroup/family. The right graph shows the distribution of the sizes of each cluster (number of IDs in each cluster). **(d)** Number of ID clusters with LRR-receptors from two different subgroups/families. The graph on the left shows the number of clusters containing LRR-receptors with the combination of subgroups/families. The graph on the right shows the distribution of the sizes of each cluster (number of IDs in each cluster). Boxplot elements: center line, median; bounds of box, 25th and 75th percentiles; whiskers, 1.5 × IQR from 25th and 75th percentiles. **(e)** Phylogenetic tree of ectodomains of BRI1/BRL members from 350 species. Branches are labelled in colours as indicated in **(a)**. The right top bar represents the distribution of LRR-RLK-Xb (blue) against LRR-RLP (pink) within the phylogenetic tree. Bottom alignment represents the alignment of IDs from LRR-RLK-Xb (blue) and LRR-RLP (pink) from the phylogenetic tree. The yellow highlights indicate the amino acid residues required for BR binding for AtBRL1^53,54^. **(f)** Phylogenetic tree of ectodomains of PSKR members from 350 species. Branches are labelled in colours as indicated in **(a)**. The right top bar represents the distribution of LRR-RLK-Xb (blue) against LRR-RLP (pink) within the phylogenetic tree. Bottom alignment represents the alignment of IDs from LRR-RLK-Xb (blue) and LRR-RLP (pink) from the phylogenetic tree. Light blue highlights indicate the amino acid residues required for PSK binding for AtPSKR1 and dark blue highlights indicate the amino acid residues required for SERK1/2 interactions for AtPSKR1^51^. **(g)** Phylogenetic tree of ectodomains of PSY1R members from 350 species. Branches are labelled with colours as indicated in **(a)**. Right top bar represents the distribution of LRR-RLK-Xb (blue) against LRR-RLP (pink) within the phylogenetic tree. Bottom alignment represents the alignment of the IDs from LRR-RLK-Xb (blue) and LRR-RLP (pink) from the phylogenetic tree. Amino acid residues that are conserved are highlighted by their properties (Clustal X^105^). For phylogenetic trees in (d-f), *Arabidopsis thaliana* genes are indicated. For the full ID and ectodomain phylogenetic tree, refer to Supplementary figure 33-34.

**Extended figure 10.**
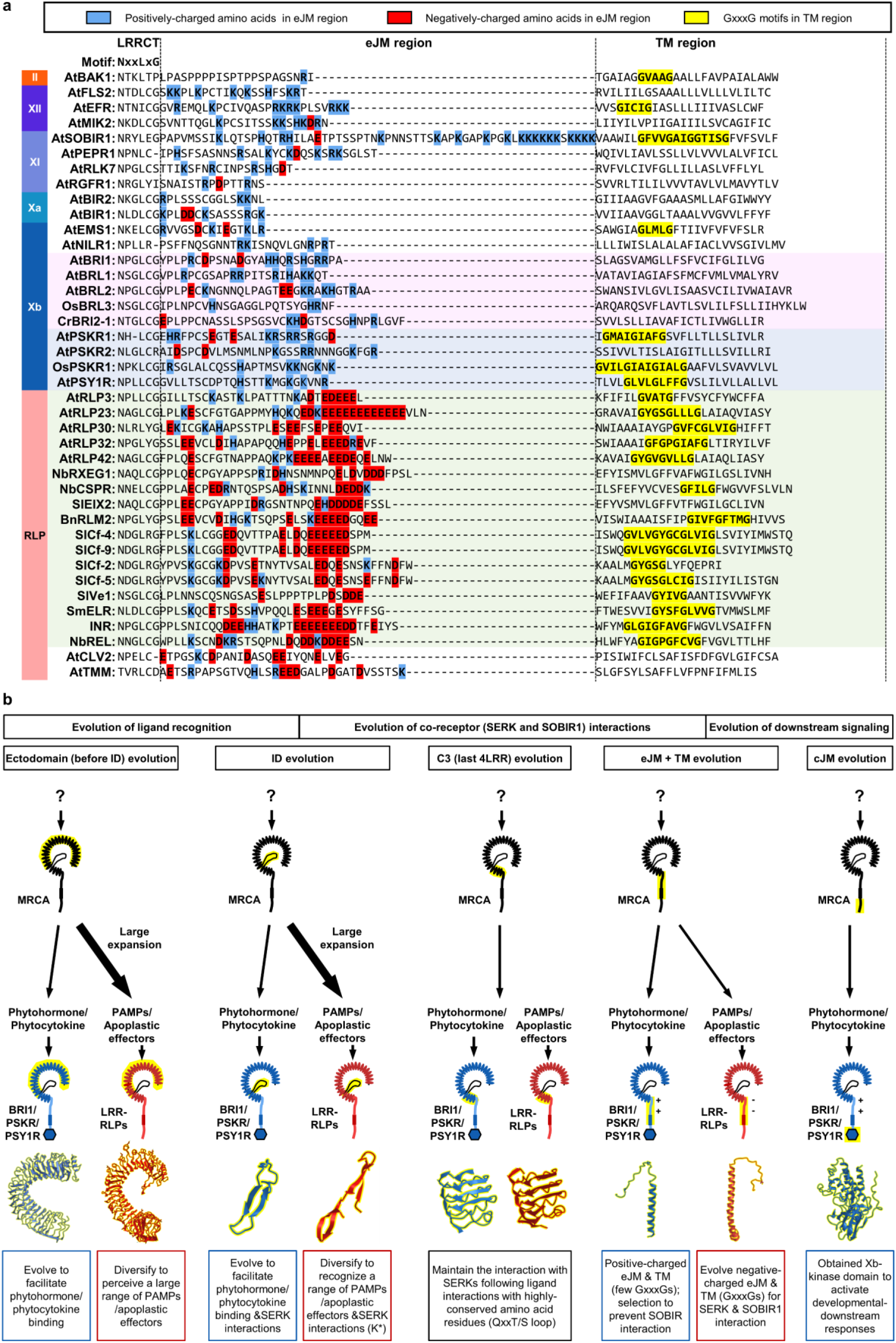
Modular evolution of cell-surface receptors in plants. **(a)** Alignment of the eJM+TM region in LRR-RLKs (from subgroup II, XII, XI, Xa, Xb) and LRR-RLPs. The BRI1/BRL clade is highlighted in purple; The PSKR/PSY1R clade is highlighted in cyan and the LRR-RLP with the ID+4LRR clade (see Figure 4c) is highlighted in light green. Positively charged amino acid residues are highlighted in blue; negatively charged amino acid residues are highlighted in red. The GxxxG motifs in TM are highlighted in yellow. **(b)** Modular evolution of different domains in cell-surface receptors to allow diverse ligand recognition and specificity of downstream signalling. Domains or regions that evolved different functions are highlighted in yellow. Bold arrows represent large expansions and diversifications. K* represents the lysine in Kx_5_Y or Yx_8_KG motifs in ID from LRR-RLPs. Domain or region structures (from left to right) are obtained from: BRI1 ectodomain (3RGX); RXEG1 ectodomain (7W3X); PSKR1 ID (4Z63); RXEG1 ID (7W3X); PSKR1 C3 (4Z63); RXEG1 C3 (7W3X); PSKR1 ejM-TM (predicted from Alphafold2^20^); RLP23 eJM-TM (predicted from Alphafold2^20^); BRI1 kinase (4OH4). Structures were visualized in iCn3D^21^.

