## Supplementary materials for "Evolutionary Trajectory of Pattern Recognition Receptors in Plants"

**Contains additional references and supplementary figure 1-37.**

**Additional references:**

For extended figure 2: *At*FLS2[1], *At*EFR[2], *Os*Xa21[3], *At*LORE[4], AtRDA2[5], AtWAK1/2[6,7], AtCRK13,28,38[8–10], AtDORN1[11], AtLecRK1.8[12], AtCERK1[13], AtFERONIA[14–18], AtCORK1[19], AtCLV1[20], AtHAESA[21], SRKs[22], OsDEES1[23], AtLecRK-IV.2[24], OsLecRK-S.7[25], AtBRI1[26], AtRGFRs[27,28], AtTMK1[29], OsSIK2[30], AtWAK4[31], AtCRK5,36[32,33], AtLecRK-A4 family (LecRKA4.1, LecRKA4.2, LecRKA4.3)[34], SlCf-4[35], AtRLP23[36], NbRXEG1[37], AtLYM1/3[38], OsLYP4/6[39], AtCLV2[40], AtTMM[41].

For extended figure 4: BAK1[42–45], SOBIR1[46], AtBIK1/AtPBL1[47,48], AtPBL19-20, 30-32, 34-40[49,50] AtCPK1,2,5,6,11,28[51–53], AtMEKK1, AtMAPKKK3/5, AtMKK4/5, AtMPK3/4/6[54], AtCNGC2,4,19,20[55–57], AtOSCA1.3, AtOSCA1.7[58], AtRbohD/F[59,60], AtEDS1[61], AtPAD4[62–64], AtSAG101[65], AtADR1[66], AtNRG1[67], AtSARD1, AtCBP60G[68], AtCPK2,11,20,24,33[69–71], AtCNGC8,16,18[72–74], AtRbohH, AtRbohJ[75], AtCPK11,12,28,30,32[76–78], AtCNGC5,6,9[79], AtRbohC[80].

For extended figure 5: AtRLP1[81], AtRLP23[36], AtRLP30[82], AtRLP32[83], AtRLP42[84], NbCSPR[85], NbRXEG1[37], SlCf-2/4/5/9[35,86–90], SlEIX2[91], SlVe1[92,93], SlI[94], SmELR[95,96], BnRLM2[97], SlCuRe1[98], VnINR[99], NbREL[100], AtCLV2[40], AtTMM[41].

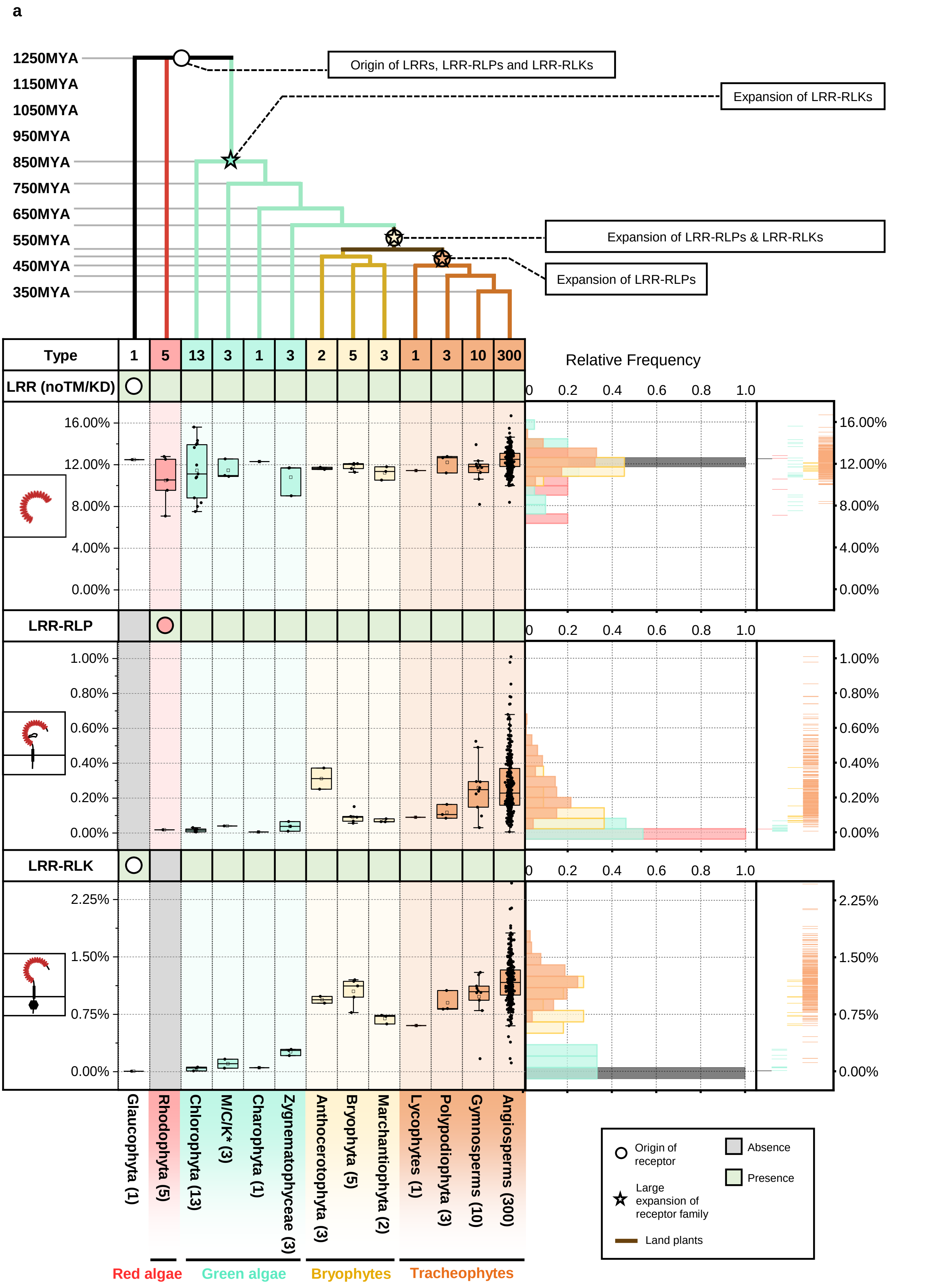
a
1250MYA
1150MYA
1050MYA
950MYA
850MYA
750MYA
650MYA
550MYA
450MYA
350MYA
Origin of LRRs, LRR-RLPs and LRR-RLKs
Expansion of LRR-RLKs
Expansion of LRR-RLPs & LRR-RLKs
Expansion of LRR-RLPs
Type
1
5
13
3
1
3
2
5
3
1
3
10
300
LRR (noTM/KD)
LRR-RLP
LRR-RLK
Relative Frequency
0
0.2
0.4
0.6
0.8
1.0
16.00%
12.00%
8.00%
4.00%
0.00%
1.00%
0.80%
0.60%
0.40%
0.20%
0.00%
2.25%
1.50%
0.75%
0.00%
Glaucophyta (1)
Rhodophyta (5)
Chlorophyta (13)
M/C/K* (3)
Charophyta (1)
Zygnematophyceae (3)
Anthocerotophyta (3)
Bryophyta (5)
Marchantiophyta (2)
Lycophytes (1)
Polypodiophyta (3)
Gymnosperms (10)
Angiosperms (300)
Red algae
Green algae
Bryophytes
Tracheophytes
Origin of receptor
Absence
Presence
Large expansion of receptor family
Land plants

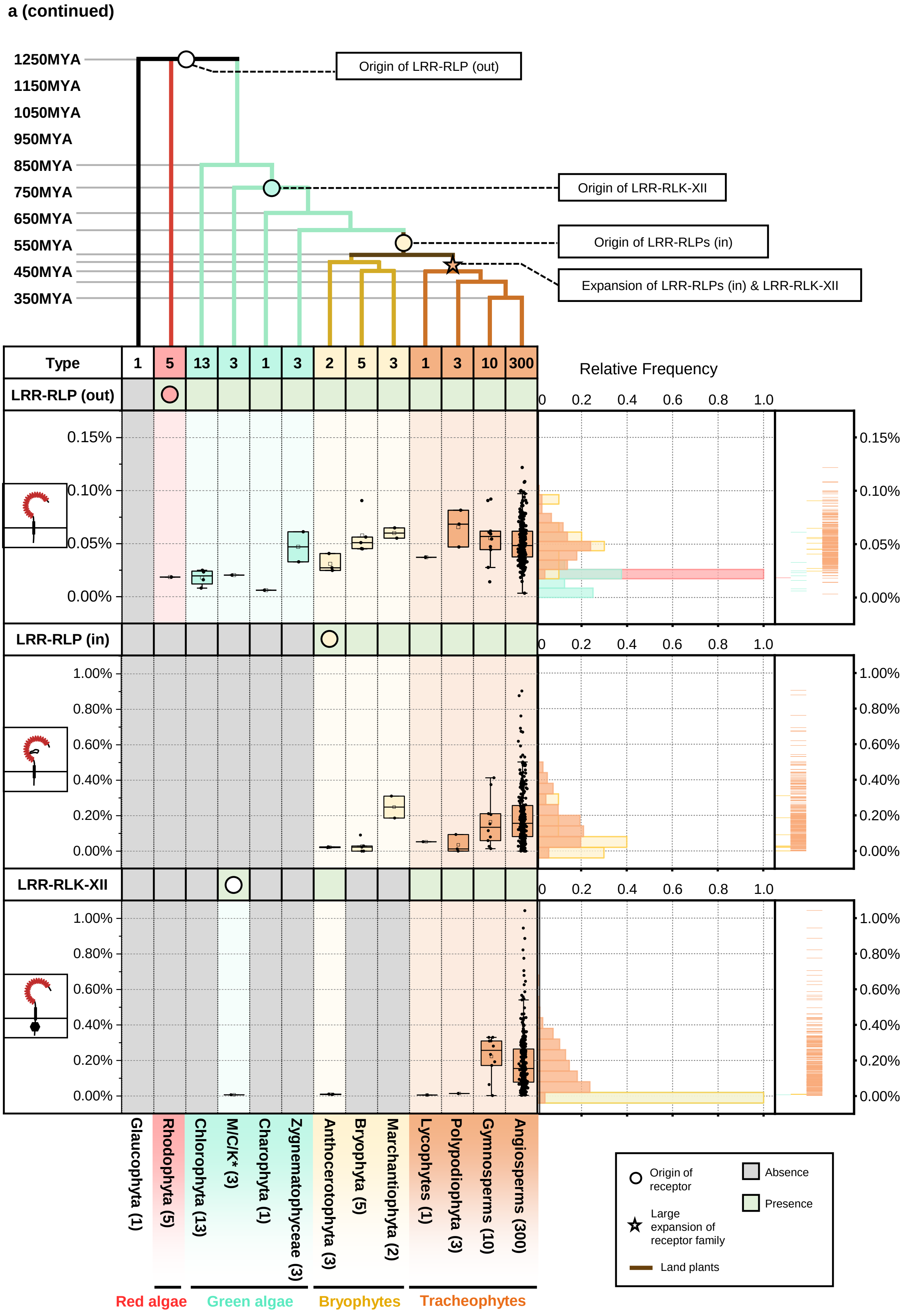
a (continued)
1250MYA
1150MYA
1050MYA
950MYA
850MYA
750MYA
650MYA
550MYA
450MYA
350MYA
Origin of LRR-RLP (out)
Origin of LRR-RLK-XII
Origin of LRR-RLPs (in)
Expansion of LRR-RLPs (in) & LRR-RLK-XII
Type
1
5
13
3
1
3
2
5
3
1
3
10
300
LRR-RLP (out)
LRR-RLP (in)
LRR-RLK-XII
Relative Frequency
0.15%
0.10%
0.05%
0.00%
1.00%
0.80%
0.60%
0.40%
0.20%
0.00%
0
0.2
0.4
0.6
0.8
1.0
Glaucophyta (1)
Rhodophyta (5)
Chlorophyta (13)
M/C/K* (3)
Charophyta (1)
Zygnematophyceae (3)
Anthocerotophyta (3)
Bryophyta (5)
Marchantiophyta (2)
Lycophytes (1)
Polypodiophyta (3)
Gymnosperms (10)
Angiosperms (300)
Red algae
Green algae
Bryophytes
Tracheophytes
Origin of receptor
Large expansion of receptor family
Land plants
Absence
Presence

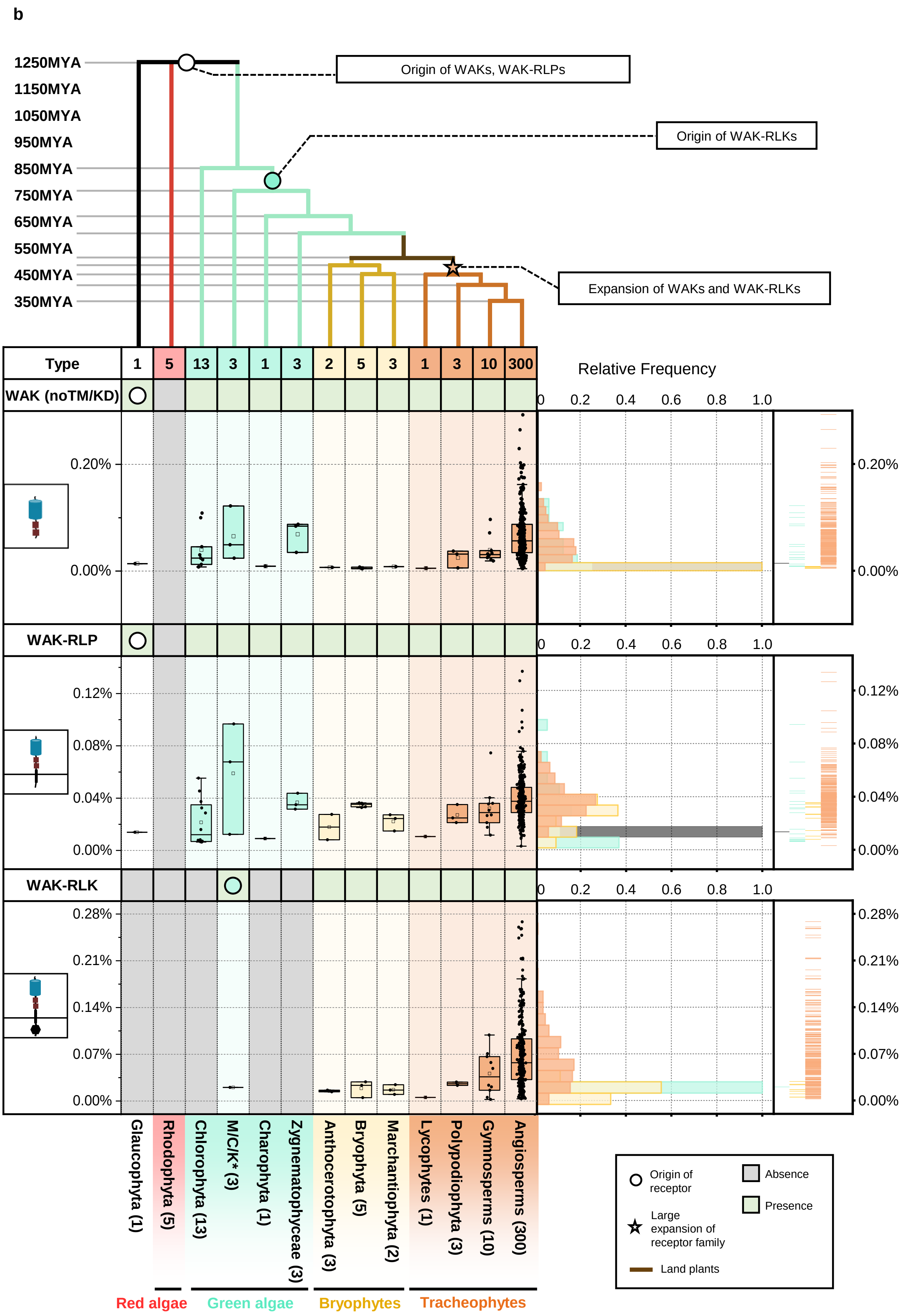
b
1250MYA
1150MYA
1050MYA
950MYA
850MYA
750MYA
650MYA
550MYA
450MYA
350MYA
Origin of WAKs, WAK-RLPs
Origin of WAK-RLKs
Expansion of WAKs and WAK-RLKs
Type
1 5 13 3 1 3 2 5 3 1 3 10 300
WAK (noTM/KD)
WAK-RLP
WAK-RLK
Relative Frequency
0 0.2 0.4 0.6 0.8 1.0
0.20%
0.00%
0.12%
0.08%
0.04%
0.28%
0.21%
0.14%
0.07%
Glaucophyta (1)
Rhodophyta (5)
Chlorophyta (13)
M/C/K* (3)
Charophyta (1)
Zygnematophyceae (3)
Anthocerotophyta (3)
Bryophyta (5)
Marchantiophyta (2)
Lycophytes (1)
Polypodiophyta (3)
Gymnosperms (10)
Angiosperms (300)
Red algae
Green algae
Bryophytes
Tracheophytes
Origin of receptor
Large expansion of receptor family
Land plants
Absence
Presence

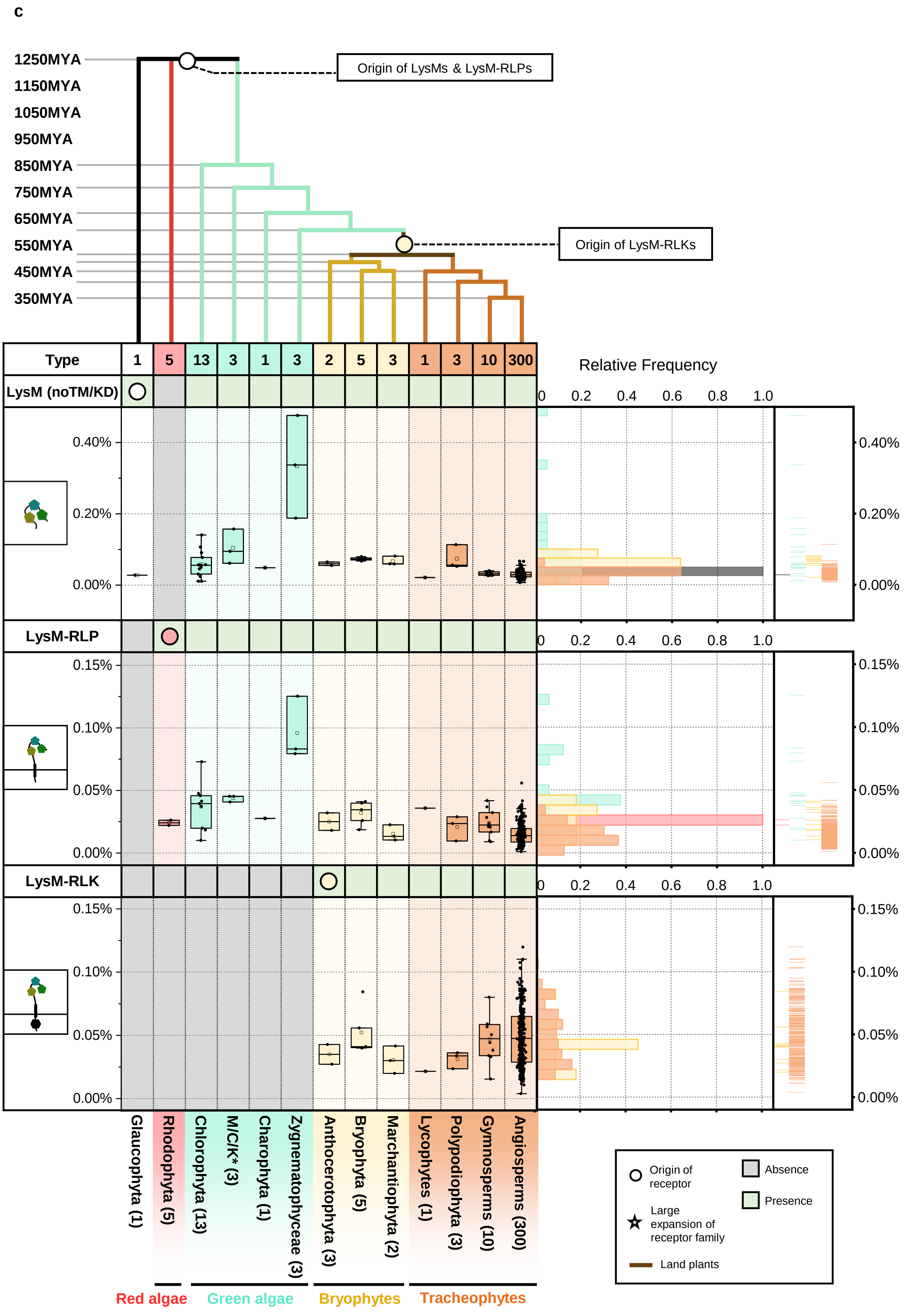
c
1250MYA
1150MYA
1050MYA
950MYA
850MYA
750MYA
650MYA
550MYA
450MYA
350MYA
Origin of LysMs & LysM-RLPs
Origin of LysM-RLKs
Type
1
5
13
3
1
3
2
5
3
1
3
10
300
LysM (noTM/KD)
LysM-RLP
LysM-RLK
Relative Frequency
0 0.2 0.4 0.6 0.8 1.0
0.40%
0.20%
0.00%
0.15%
0.10%
0.05%
Glaucophyta (1)
Rhodophyta (5)
Chlorophyta (13)
M/C/K* (3)
Charophyta (1)
Zygnematophyceae (3)
Anthocerotophyta (3)
Bryophyta (5)
Marchantiophyta (2)
Lycophytes (1)
Polypodiophyta (3)
Gymnosperms (10)
Angiosperms (300)
Red algae
Green algae
Bryophytes
Tracheophytes
Origin of receptor
Absence
Presence
Large expansion of receptor family
Land plants

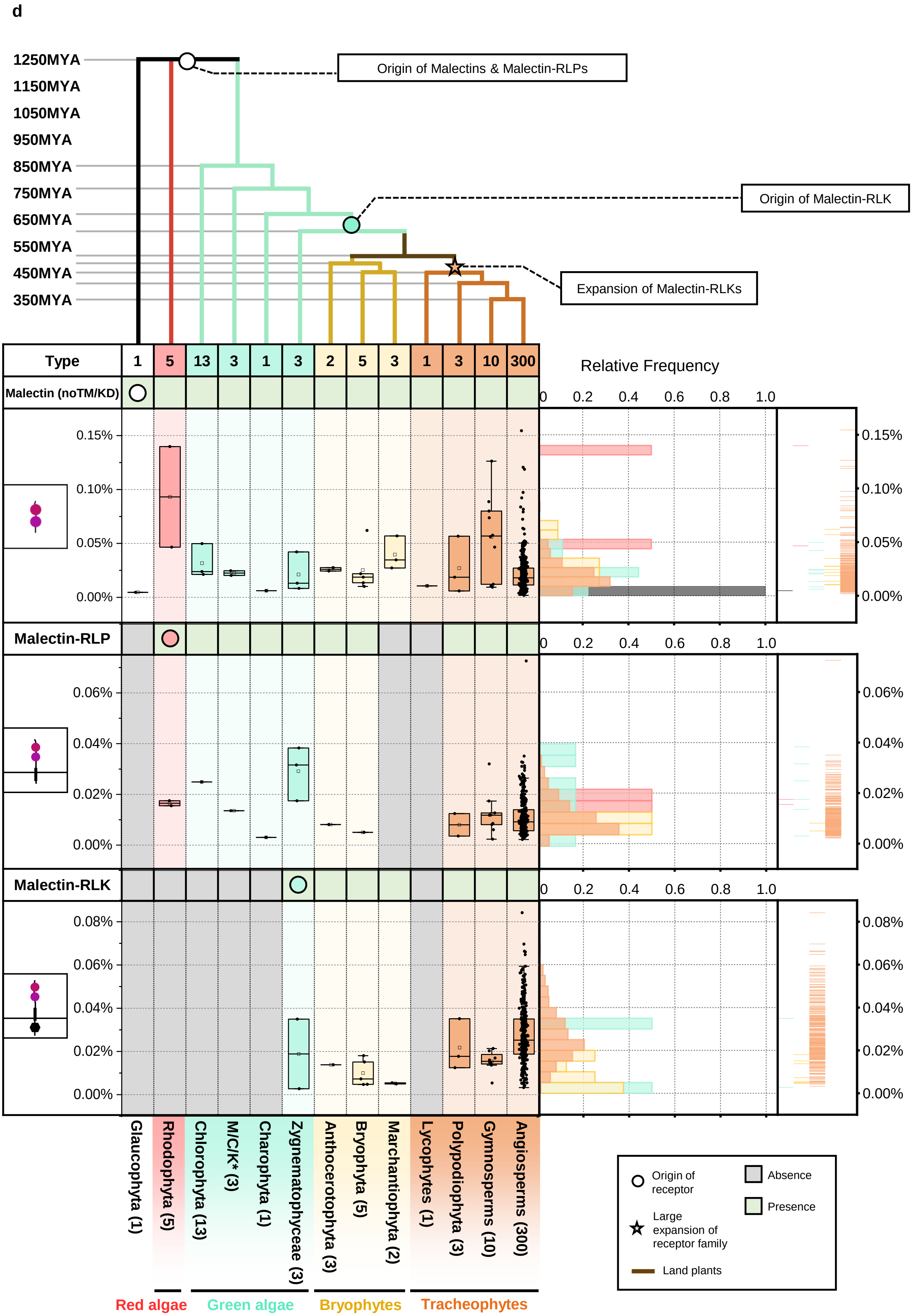
d
1250MYA
1150MYA
1050MYA
950MYA
850MYA
750MYA
650MYA
550MYA
450MYA
350MYA
Origin of Malectins & Malectin-RLPs
Origin of Malectin-RLK
Expansion of Malectin-RLKs
Type
1
5
13
3
1
3
2
5
3
1
3
10
300
Malectin (noTM/KD)
Malectin-RLP
Malectin-RLK
Relative Frequency
0.2
0.4
0.6
0.8
1.0
0.15%
0.10%
0.05%
0.00%
0.06%
0.04%
0.02%
0.08%
Glaucophyta (1)
Rhodophyta (5)
Chlorophyta (13)
M/C/K* (3)
Charophyta (1)
Zygnematophyceae (3)
Anthocerotophyta (3)
Bryophyta (5)
Marchantiophyta (2)
Lycophytes (1)
Polypodiophyta (3)
Gymnosperms (10)
Angiosperms (300)
Red algae
Green algae
Bryophytes
Tracheophytes
Origin of receptor
Large expansion of receptor family
Land plants
Absence
Presence

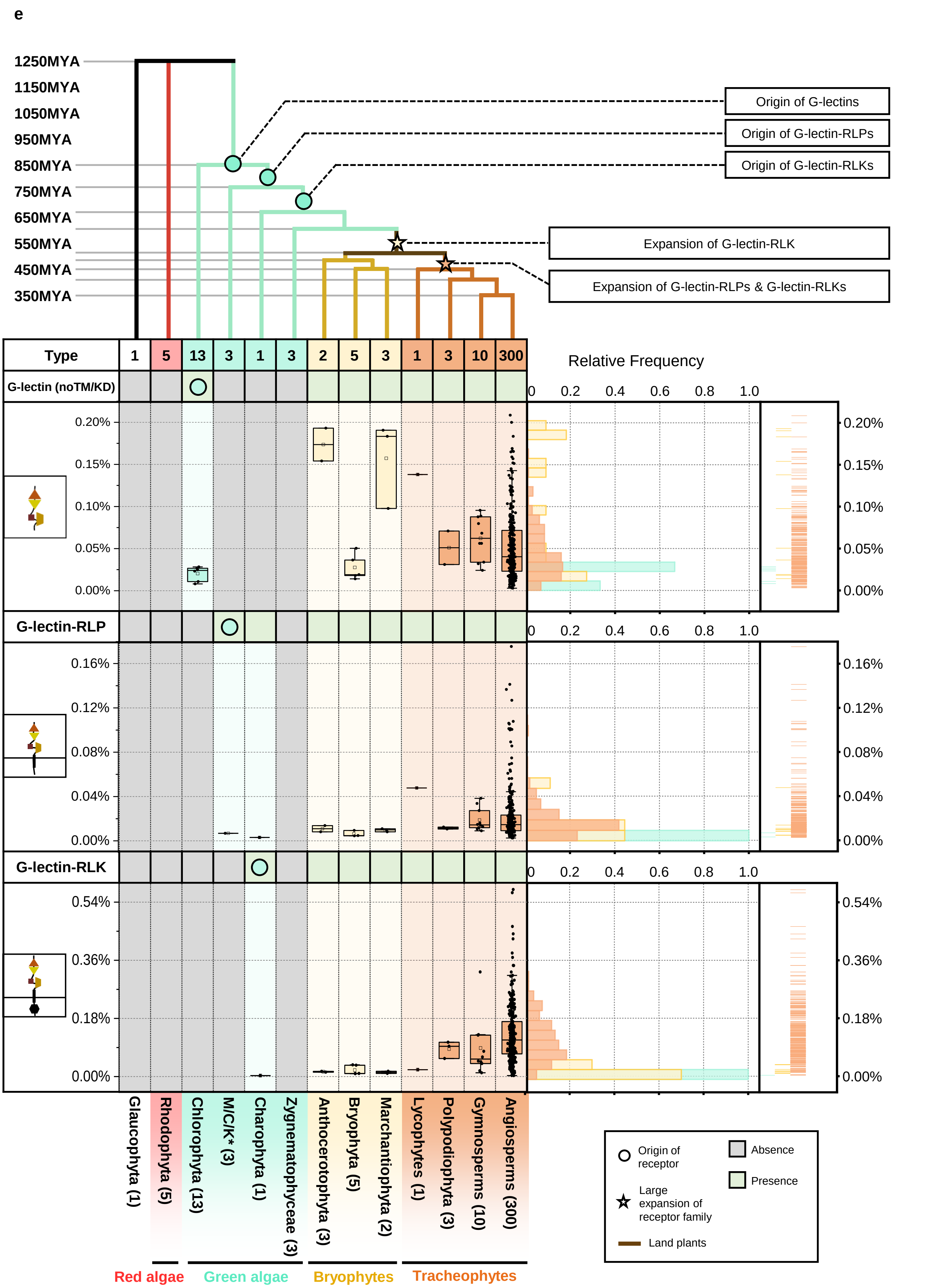
e
1250MYA
1150MYA
1050MYA
950MYA
850MYA
750MYA
650MYA
550MYA
450MYA
350MYA
Origin of G-lectins
Origin of G-lectin-RLPs
Origin of G-lectin-RLKs
Expansion of G-lectin-RLK
Expansion of G-lectin-RLPs & G-lectin-RLKs
Type
1
5
13
3
1
3
2
5
3
1
3
10
300
G-lectin (noTM/KD)
G-lectin-RLP
G-lectin-RLK
Relative Frequency
0 0.2 0.4 0.6 0.8 1.0
0.20%
0.15%
0.10%
0.05%
0.00%
0.16%
0.12%
0.08%
0.04%
0.54%
0.36%
0.18%
Glaucophyta (1)
Rhodophyta (5)
Chlorophyta (13)
M/C/K* (3)
Charophyta (1)
Zygnematophyceae (3)
Anthocerotophyta (3)
Bryophyta (5)
Marchantiophyta (2)
Lycophytes (1)
Polypodiophyta (3)
Gymnosperms (10)
Angiosperms (300)
Red algae
Green algae
Bryophytes
Tracheophytes
Origin of receptor
Large expansion of receptor family
Land plants
Absence
Presence

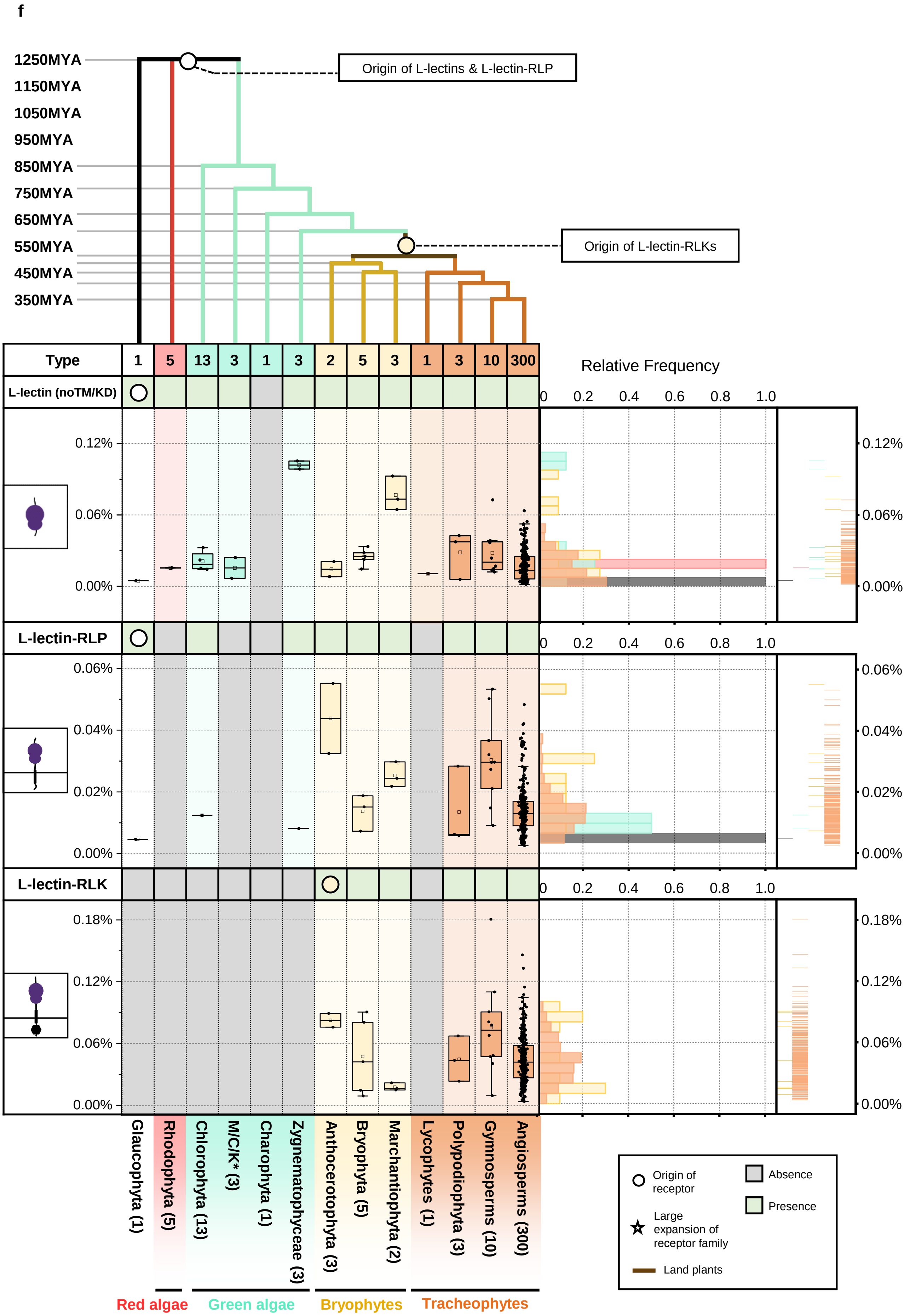
f
1250MYA
1150MYA
1050MYA
950MYA
850MYA
750MYA
650MYA
550MYA
450MYA
350MYA
Origin of L-lectins & L-lectin-RLP
Origin of L-lectin-RLKs
Type
1
5
13
3
1
3
2
5
3
1
3
10
300
L-lectin (noTM/KD)
L-lectin-RLP
L-lectin-RLK
Relative Frequency
0 0.2 0.4 0.6 0.8 1.0
0.12%
0.06%
0.00%
0.04%
0.02%
0.18%
Glaucophyta (1)
Rhodophyta (5)
Chlorophyta (13)
M/C/K* (3)
Charophyta (1)
Zygnematophyceae (3)
Anthocerotophyta (3)
Bryophyta (5)
Marchantiophyta (2)
Lycophytes (1)
Polypodiophyta (3)
Gymnosperms (10)
Angiosperms (300)
Red algae
Green algae
Bryophytes
Tracheophytes
Origin of receptor
Large expansion of receptor family
Land plants
Absence
Presence

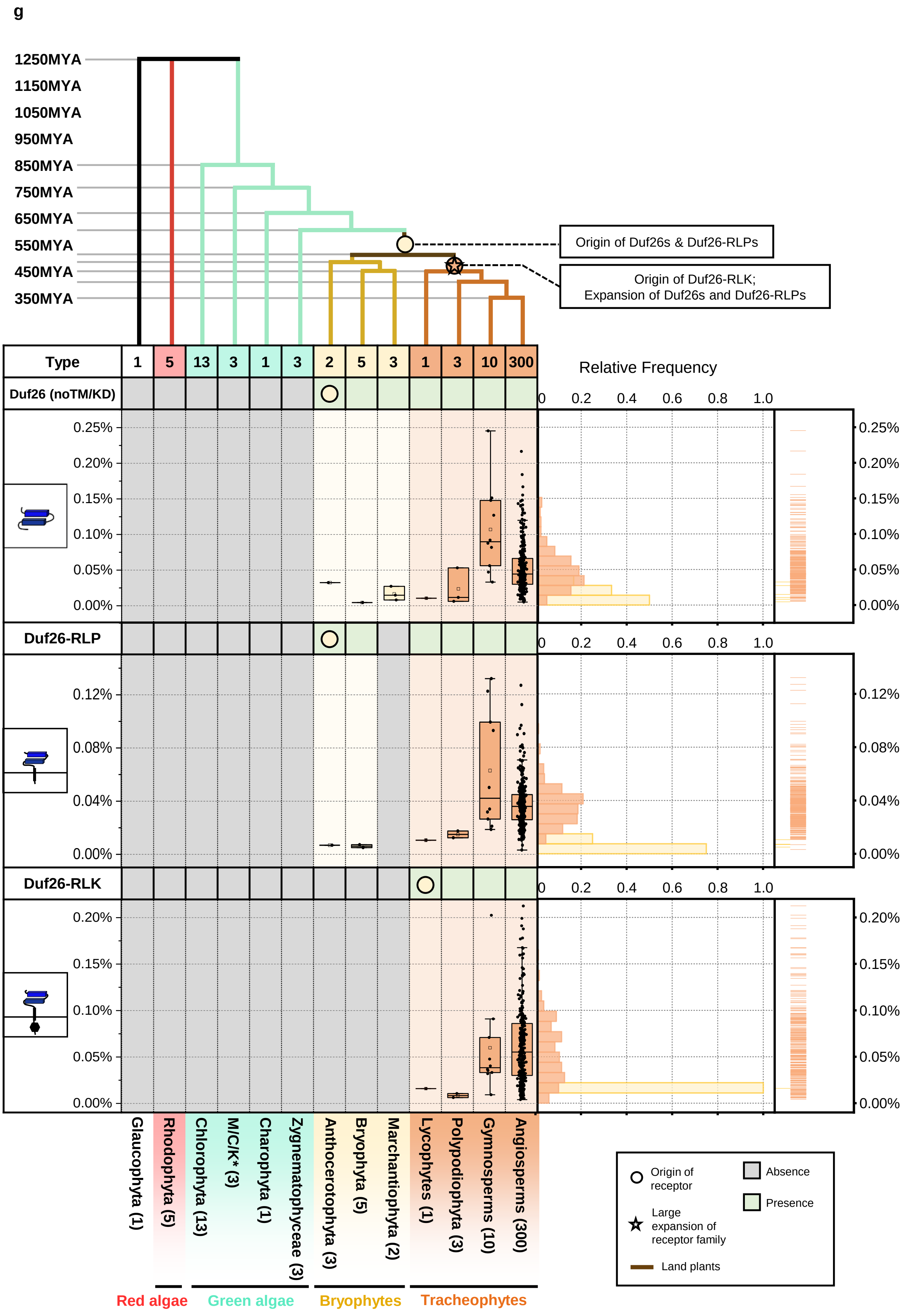
g
1250MYA
1150MYA
1050MYA
950MYA
850MYA
750MYA
650MYA
550MYA
450MYA
350MYA
Origin of Duf26s & Duf26-RLPs
Origin of Duf26-RLK;
Expansion of Duf26s and Duf26-RLPs
Type
1
5
13
3
1
3
2
5
3
1
3
10
300
Duf26 (noTM/KD)
Duf26-RLP
Duf26-RLK
Relative Frequency
0 0.2 0.4 0.6 0.8 1.0
0.25%
0.20%
0.15%
0.10%
0.05%
0.00%
0.12%
0.08%
0.04%
Glaucophyta (1)
Rhodophyta (5)
Chlorophyta (13)
M/C/K* (3)
Charophyta (1)
Zygnematophyceae (3)
Anthocerotophyta (3)
Bryophyta (5)
Marchantiophyta (2)
Lycophytes (1)
Polypodiophyta (3)
Gymnosperms (10)
Angiosperms (300)
Red algae
Green algae
Bryophytes
Tracheophytes
Origin of receptor
Large expansion of receptor family
Land plants
Absence
Presence

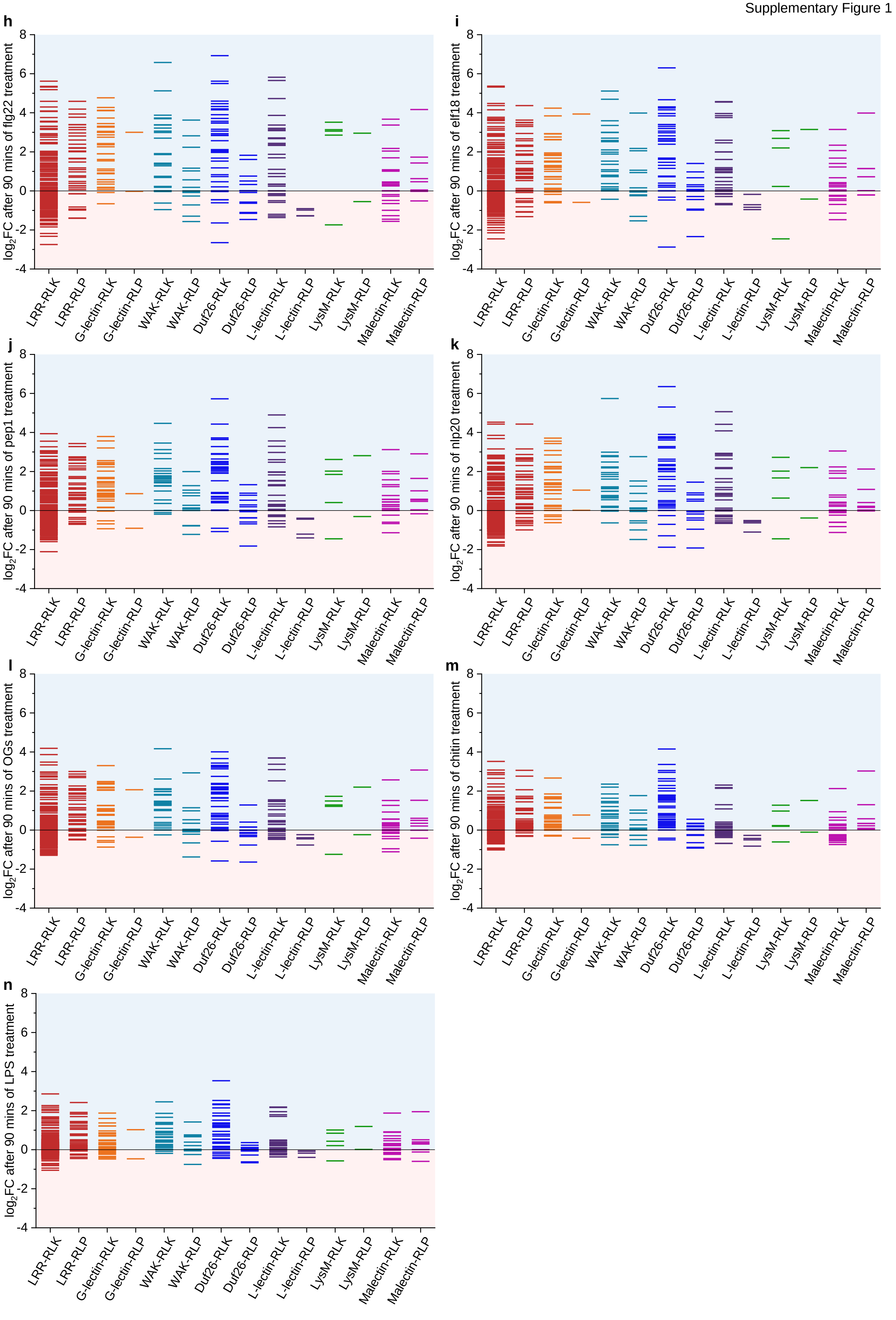
h
log$_2$FC after 90 mins of flg22 treatment
i
log$_2$FC after 90 mins of elf18 treatment
j
log$_2$FC after 90 mins of pep1 treatment
k
log$_2$FC after 90 mins of nlp20 treatment
l
log$_2$FC after 90 mins of OGs treatment
m
log$_2$FC after 90 mins of chitin treatment
n
log$_2$FC after 90 mins of LPS treatment
8
6
4
2
0
-2
-4
LRR-RLK
LRR-RLP
G-lectin-RLK
G-lectin-RLP
WAK-RLK
WAK-RLP
Duf26-RLK
Duf26-RLP
L-lectin-RLK
L-lectin-RLP
LysM-RLK
LysM-RLP
Malectin-RLK
Malectin-RLP

**Supplementary figure 1. The origin, expansion, and expression of cell-surface receptors in Viridiplantae.** The origin and expansion of **(a)** LRR-, **(a continued)** LRR-RLP subgroups and LRR-RLK-XII, **(b)** WAK-, **(c)** LysM-, **(d)** Malectin, **(e)** G-lectin, **(f)** L-lectin, and **(g)** Duf26-domains in plants. Top panel represents a phylogenetic tree of multiple algal and plant lineages. Circles (○) and stars (☆) indicate the origin and expansion of receptor families. The timescale (in million years; MYA) of the phylogenetic tree is estimated by TIMETREE. Bottom panel represents the present/absence of an ectodomain (with no TM/KD), ectodomain-RLP, ectodomain-RLK in different algal and plant lineages. Grey box indicates the absence of receptor and green box indicates the presence of receptors in each lineage. *M/C/K represents Mesostigmatophyceae, Chlorokybophyceae and Klebsormidiophyceae. Number of available species from algae and plant lineages are indicated by numbers in the boxes. Boxplot below represents the percentage (%) of ectodomains (in proteins without TM or KD), ectodomain-RLPs and ectodomain-RLKs in each lineage. Right plot represents the distribution of the relative frequency of the percentage of ectodomains, ectodomain-RLPs and ectodomain-RLKs in each lineage. **(h-n)** The expression of cell-surface receptors during PTI in *Arabidopsis thaliana*. *A. thaliana* seedlings were treated with flg22, elf18, pep1, nlp20, OGs, chitin, or LPS to activate PTI. Light blue represents increased expression and light pink represents decreased expression during PTI. X-axis values represent log2 (fold change during PTI relative to samples at 0 min after PAMP/DAMP treatment). RNA-seq data analyzed here were reported previously, where PTI was activated by different PAMPs/DAMPs in *A. thaliana* for 90mins. RNA-seq data were obtained from Bjornson et al, Nature Plants 2021 (reference 15 in main text).

### SERK phylogenetic tree

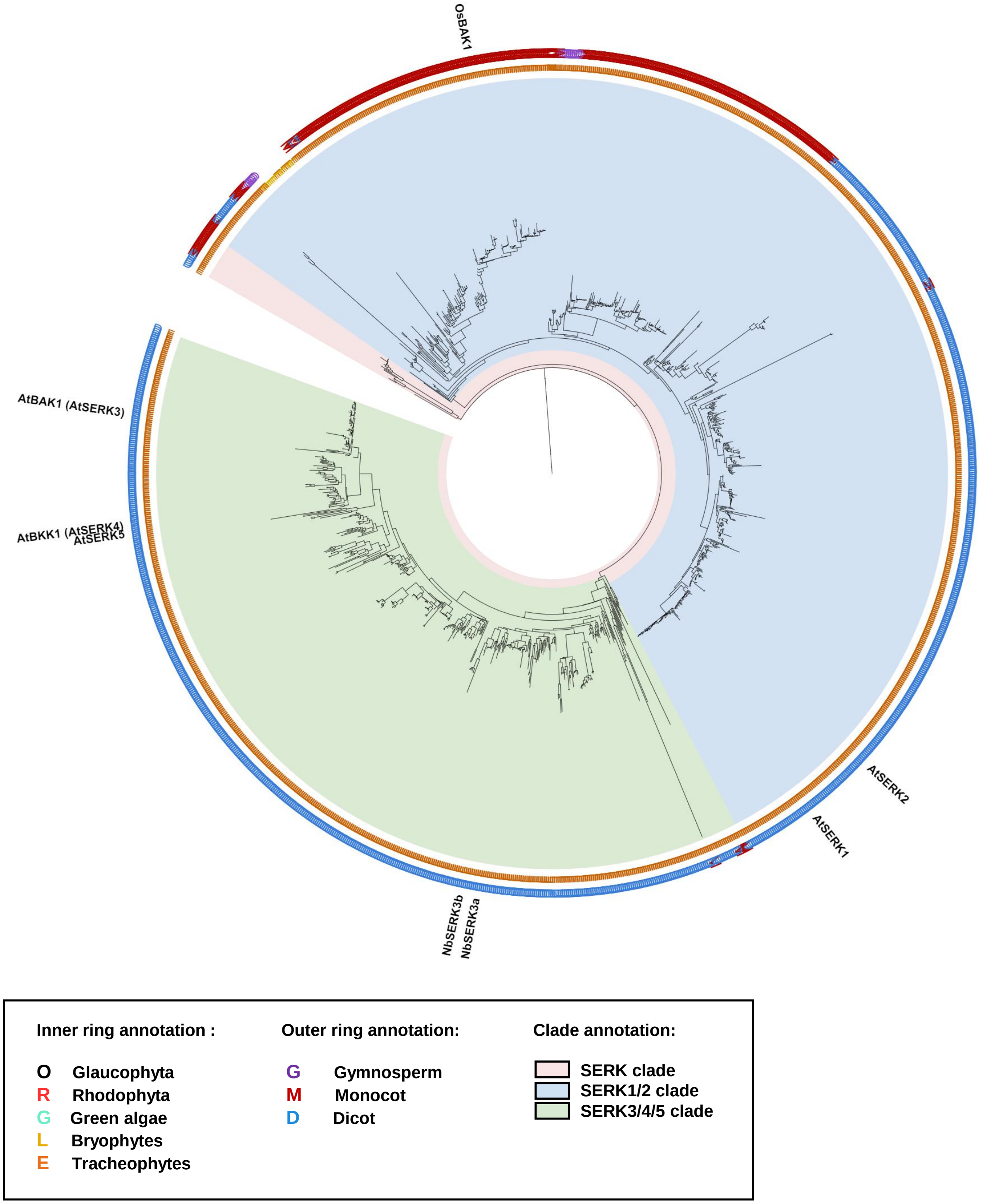


**Supplementary figure 2. Phylogenetic analysis of SERKs in plants.** Full-length phylogenetic tree of SERK members identified from the LRR-RLK-II phylogenetic tree. The inner ring indicates SERK members from Glaucophyta, red algae (Rhodophyta), green algae, Bryophytes or Tracheophytes. The outer ring indicates LRR-RLK-II members from gymnosperm, monocots or dicots. The SERK, SERK1/2 and SERK3/4/5 clades are defined. Characterized SERK members are labelled in the tree. Abbreviations for plant species: *A. thaliana*, At; *N. benthamiana*, Nb; *O.sativa*, Os.

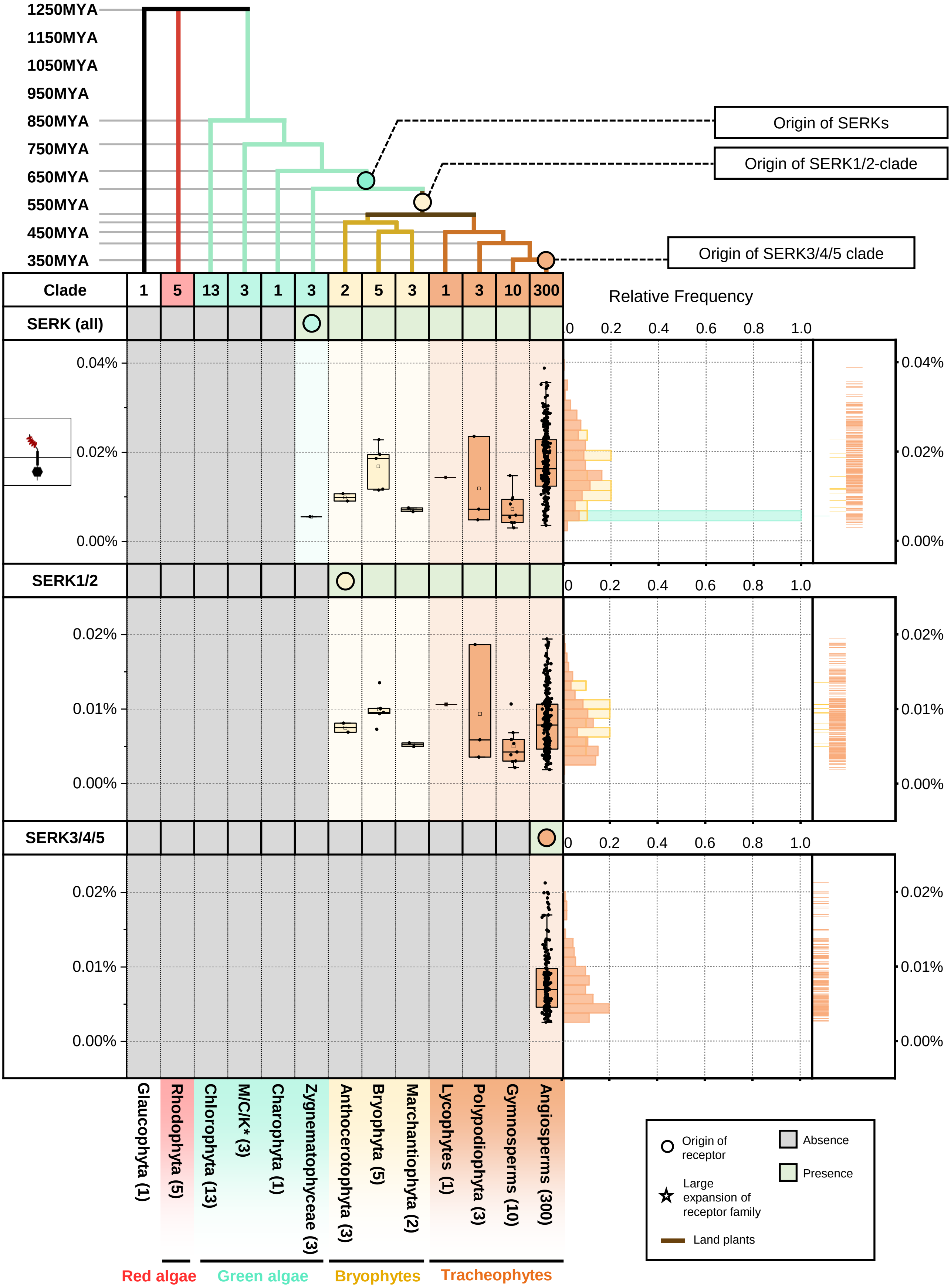


**Supplementary figure 3. The origin and expansion of SERKs in plants.** Top panel represents the phylogenetic tree of multiple algae and plant lineage. With circle (○) and star (☆) indicate the origin and expansion of receptor families. The timescale (in million year; MYA) of the phylogenetic tree is estimated by TIMETREE. Bottom panel represent the present/absence of SERKs, SERK1/2-related and SERK3/4/5-related proteins in different algae and plant lineages. Grey box indicates the absence of receptor and green box indicates the presence of receptors in each lineage. The origin of SERKs are marked with a circle. *M/C/K represents Mesostigmatophyceae, Chlorokybophyceae and Klebsormidiophyceae. Number of available species from each algae and plant lineages are indicated by numbers in the boxes. Boxplot below represents the percentage (%) and right plot represents the distribution and rug of the relative frequency of the % of SERKs, SERK1/2-related and SERK3/4/5-related proteins in each lineage.

### SOBIR1 phylogenetic tree

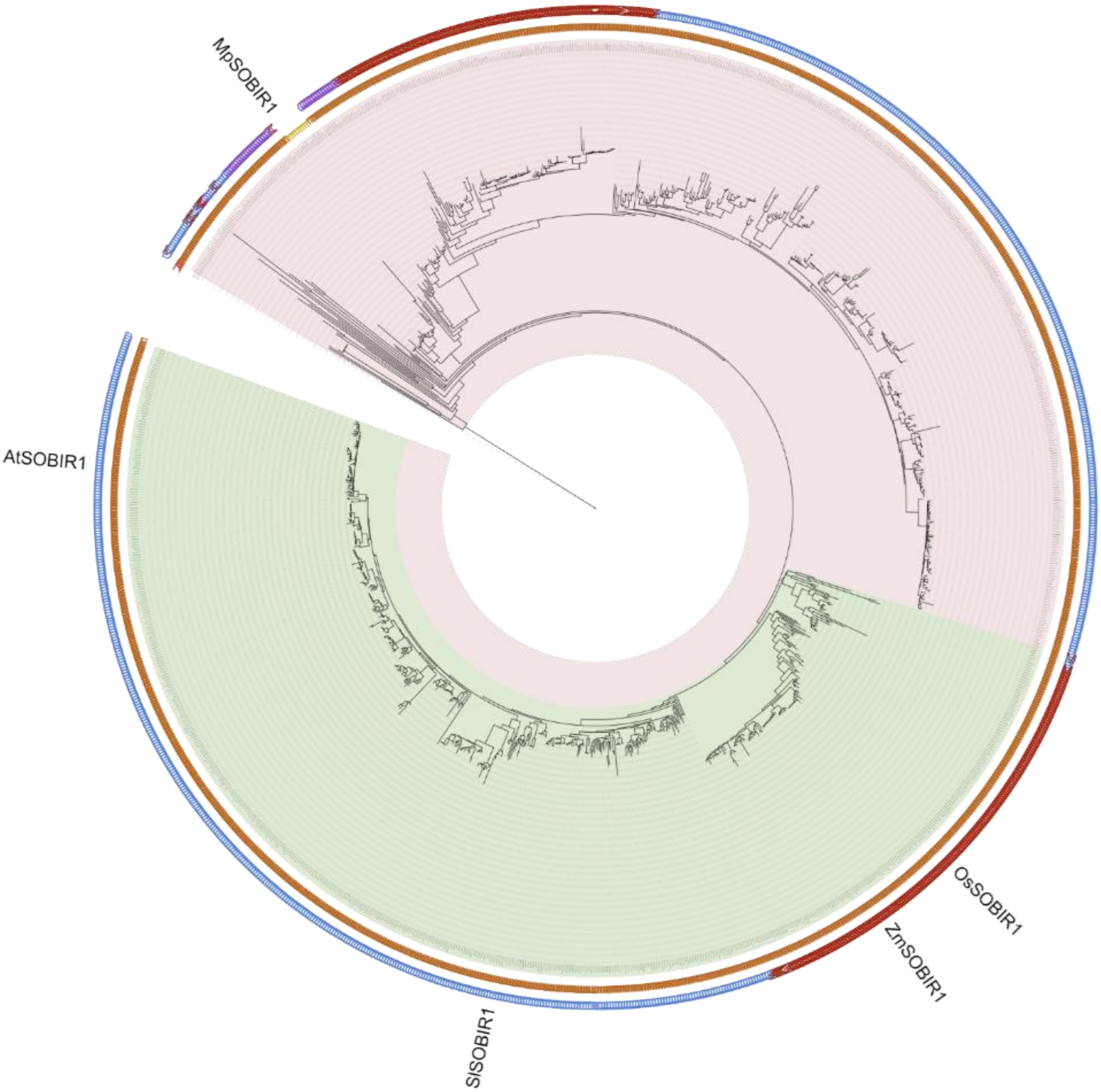


**Inner ring annotation :**

- **O** Glaucophyta
- **R** Rhodophyta
- **G** Green algae
- **L** Bryophytes
- **E** Tracheophytes

**Outer ring annotation:**

- **G** Gymnosperm
- **M** Monocot
- **D** Dicot

**Clade annotation:**

- SOBIR1/SOBIR1-related clade
- Angiosperm SOBIR1 clade

**Supplementary figure 4. Phylogenetic analysis of SOBIR1 in plants.** Phylogenetic tree of full length SOBIR1 members identified from 350 species. The inner ring indicates SOBIR1 members from Glaucophyta, red algae (Rhodophyta), green algae, Bryophytes or Tracheophytes. The outer ring indicates SOBIR1 members from gymnosperm, monocots or dicots. The SOBIR1/SOBIR1-related and angiosperm SOBIR1 clades are defined. Characterized SOBIR1 members are labelled in the tree. Abbreviations for plant species: *M. polymorpha*, Mp; *A. thaliana*, At; *S. lycopersicum*, Sl; *O. sativa*, Os; *Z. mays*, ZM.

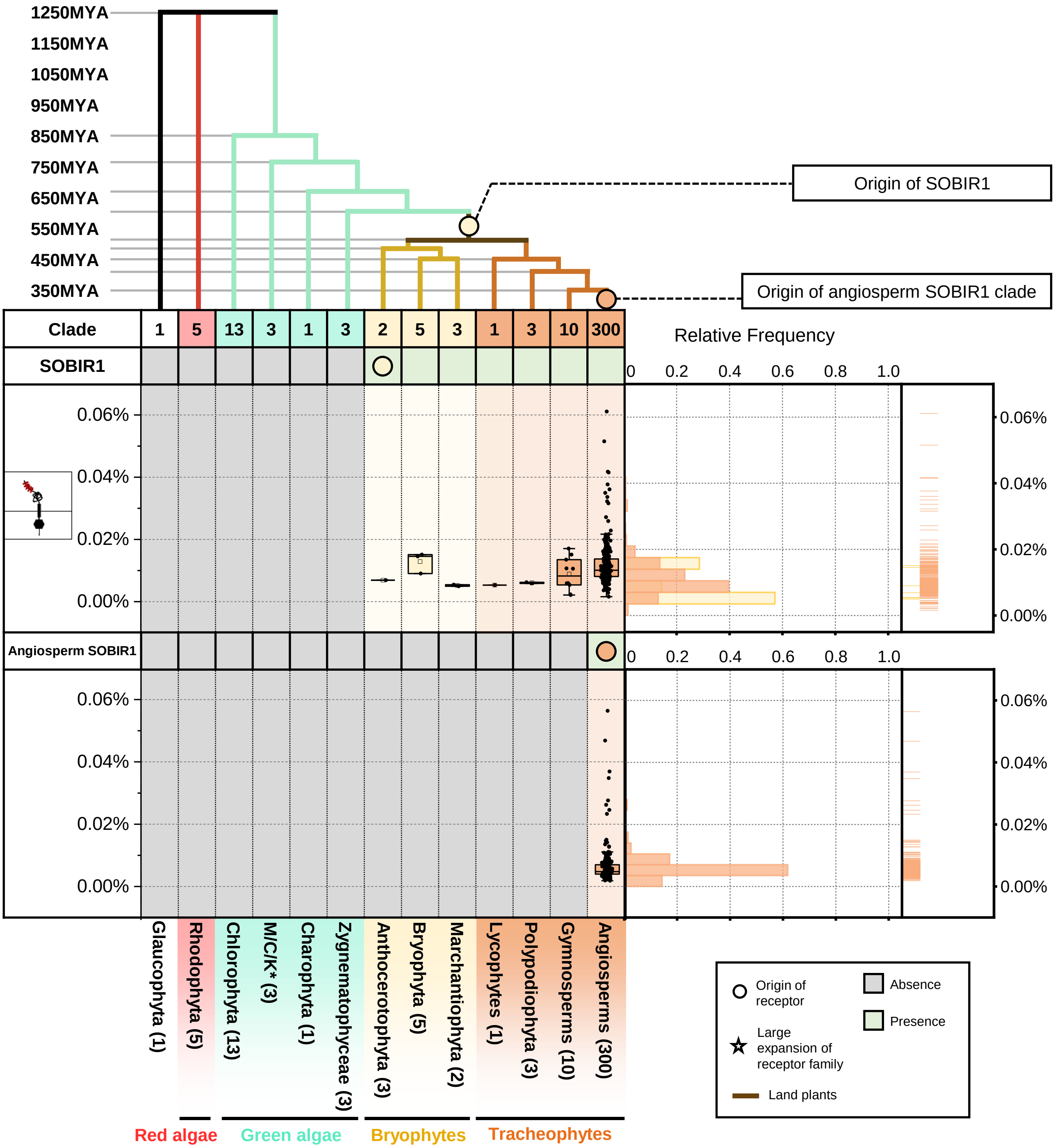


**Supplementary figure 5. The origin and expansion of SOBIR1 in plants.** Top panel represents a phylogenetic tree of multiple alga and plant lineages. Circles (○) and stars (☆) indicate the origin and expansion of receptor families. The timescale (in million years; MYA) of the phylogenetic tree is estimated by TIMETREE. Bottom panel represents the presence/absence of SOBIR1 and angiosperm SOBIR1 in different algae and plant lineages. Grey box indicates the absence of receptor and green box indicates the presence of receptors in each lineage. The origin of SOBIR1 is marked with a circle. *M/C/K represents Mesostigmatophyceae, Chlorokybophyceae and Klebsormidiophyceae. Number of available species from each algal and plant lineage are indicated by numbers in the boxes. Boxplot below represents the percentage (%) and right plot represents the distribution and rug of the relative frequency of the % of SOBIR1 and angiosperm SOBIR1 in each lineage.

### RLCK-VII phylogenetic tree

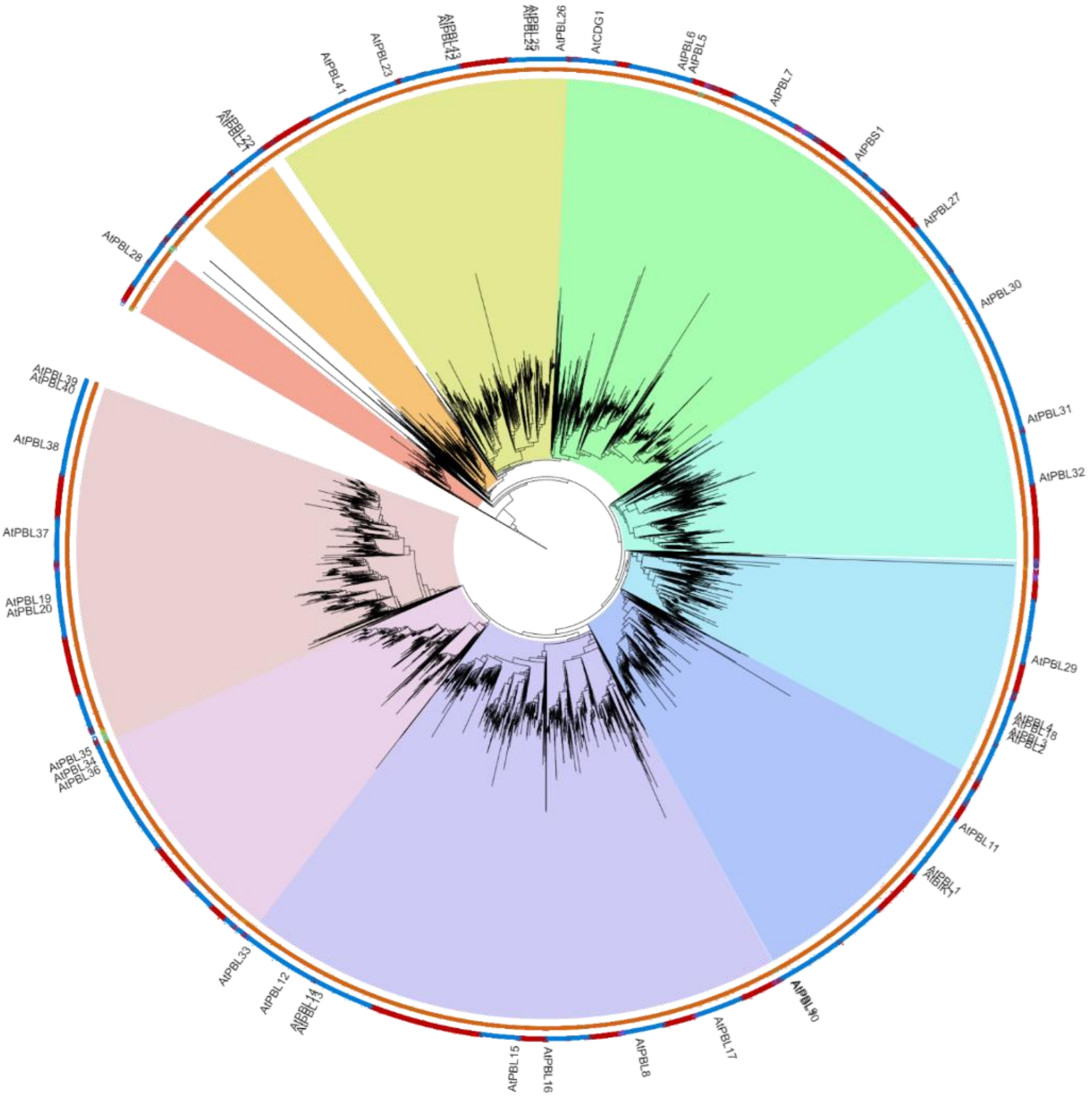


**Inner ring annotation :**

- O Glaucophyta
- R Rhodophyta
- G Green algae
- L Bryophytes
- E Tracheophytes

**Outer ring annotation:**

- G Gymnosperm
- M Monocot
- D Dicot

**Clade annotation:**

- RLCK-VII clade
- RLCK-VII-10 clade (AtPBL28)
- RLCK-VII-3 clade (AtPBL21,22)
- RLCK-VII-2 clade (AtPBL23-26,41-43)
- RLCK-VII-1 clade (AtPBL5-7,27, AtPBS1, AtCDG1)
- RLCK-VII-7 clade (AtPBL30-32)
- RLCK-VII-9 clade (AtPBL2-4,18,29)
- RLCK-VII-8 clade (AtPBL9-11, AtBIK1, AtPBL1)
- RLCK-VII-6 clade (AtPBL8,12-17,33)
- RLCK-VII-5 clade (AtPBL34-36)
- RLCK-VII-4 clade (AtPBL19,20,37-40)

**Supplementary figure 6. Phylogenetic analysis of RLCK-VII in plants.** Phylogenetic tree of full length RLCK-VII members identified from 350 species. The inner ring indicates RLCK-VII members from either Glaucophyta, red algae (Rhodophyta), green algae, Bryophytes or Tracheophytes. The outer ring indicates RLCK-VII members from either gymnosperm, monocots or dicots. The 10 RLCK-VII clades are defined based on previous annotation[101]. *Arabidopsis thaliana* RLCK-VIIs members are labelled in the tree. Abbreviations for plant species: *A. thaliana*, At.

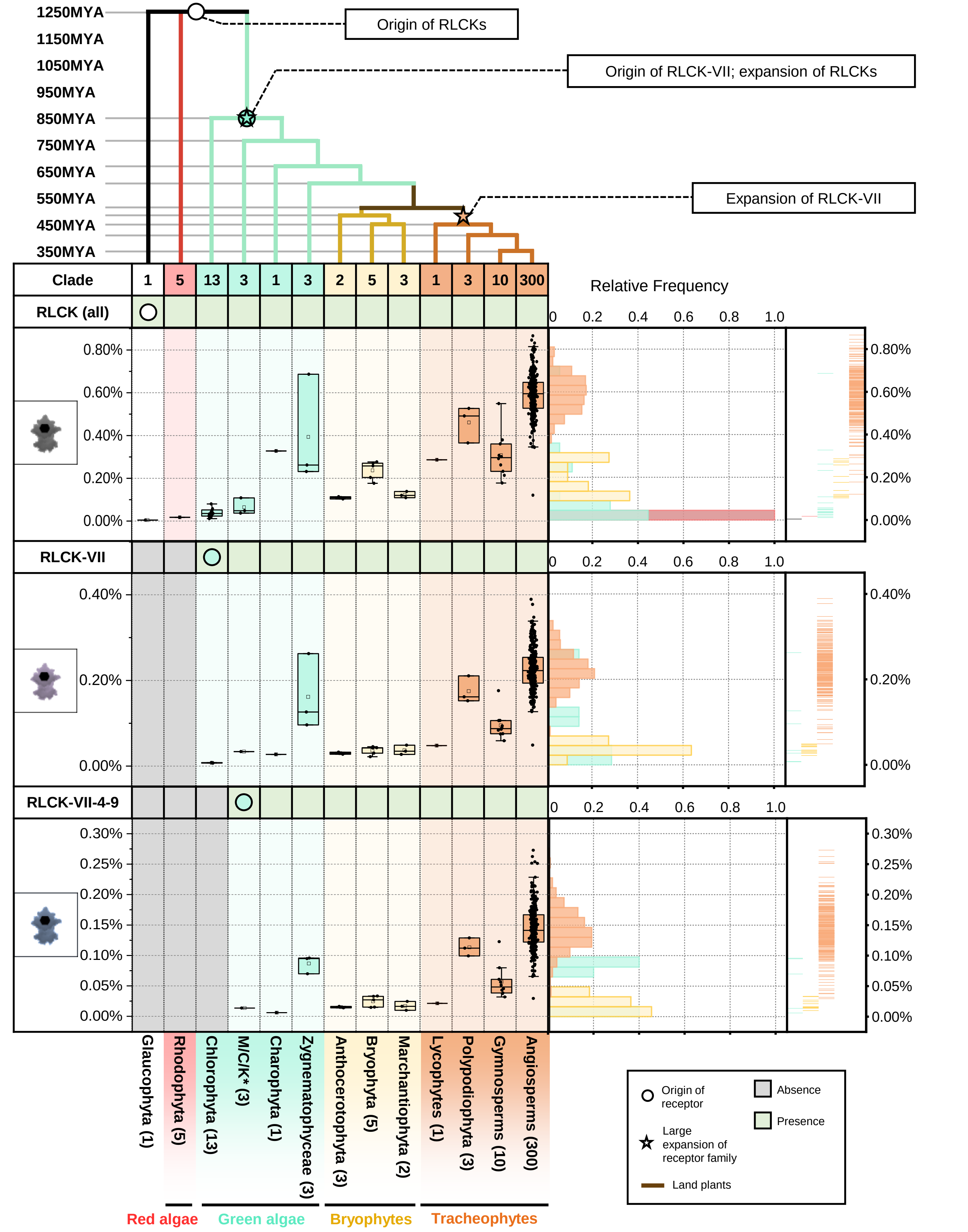


**Supplementary figure 7. The origin and expansion of RLCKs in plants.** Top panel represents the phylogenetic tree of multiple algae and plant lineage. With circle (○) and star (☆) indicate the origin and expansion of receptor families. The timescale (in million year; MYA) of the phylogenetic tree is estimated by TIMETREE. Bottom panel represent the present/absence of RLCKs, RLCK-VII and RLCK-VII-4,5,6,7,8,9 in different algae and plant lineages. Grey box indicates the absence of receptor and green box indicates the presence of receptors in each lineage. The origin of RLCKs are marked with a circle. *M/C/K represents Mesostigmatophyceae, Chlorokybophyceae and Klebsormidiophyceae. Number of available species from each algae and plant lineages are indicated by numbers in the boxes. Boxplot below represents the percentage (%) and right plot represents the distribution and rug of the relative frequency of the % of RLCKs, RLCK-VII and RLCK-VII-4,5,6,7,8,9 in each lineage.

### CDPK phylogenetic tree

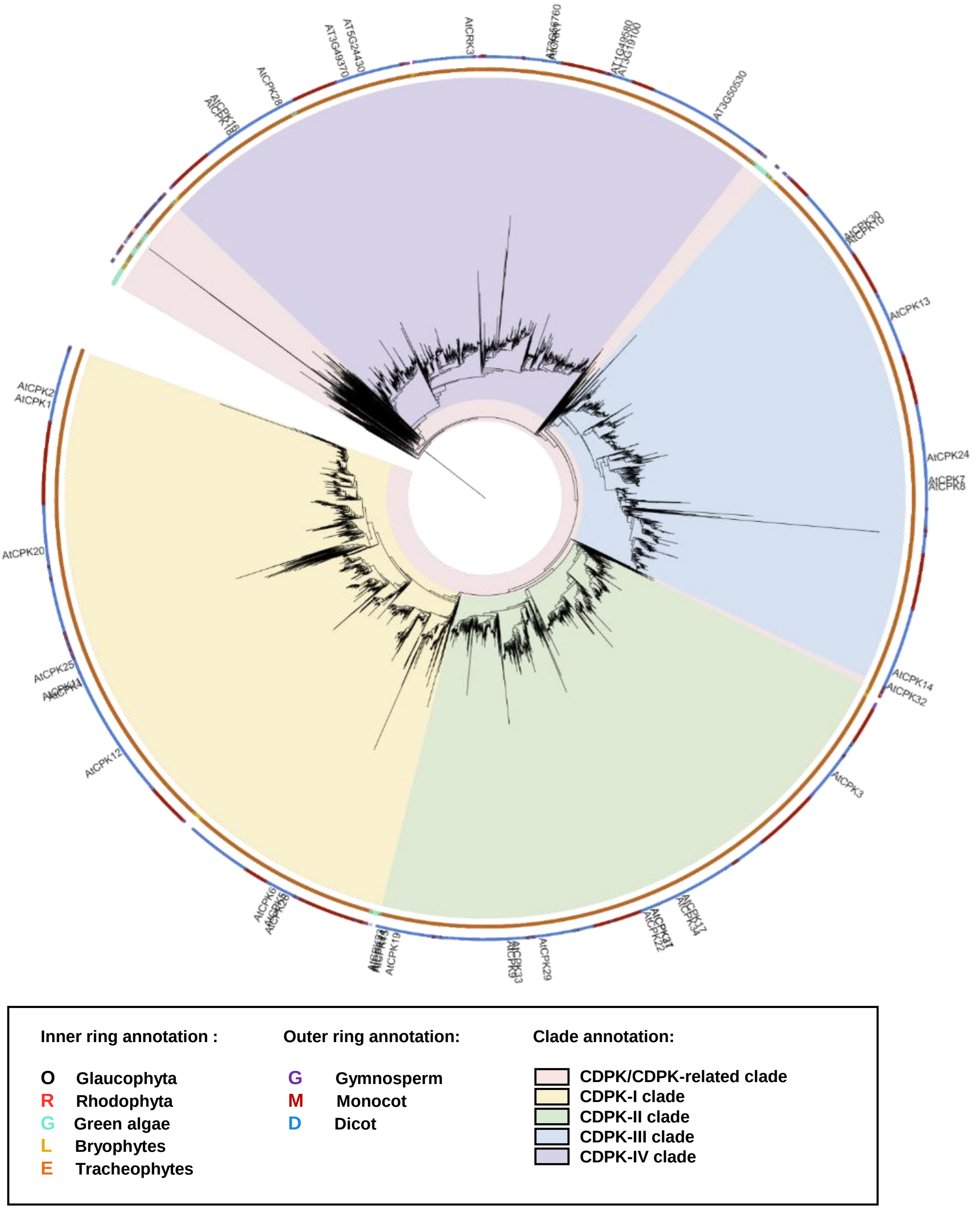


**Supplementary figure 8. Phylogenetic analysis of CDPK in plants.** Phylogenetic tree of full length CDPK members identified from 350 species. The inner ring indicates CDPK members from either Glaucophyta, red algae (Rhodophyta), green algae, Bryophytes or Tracheophytes. The outer ring indicates CDPK members from either gymnosperm, monocots or dicots. The CDPK-I/II/III/IV clades are defined based on previous annotation[102,103]. *Arabidopsis thaliana* CDPK members are labelled in the tree. Abbreviations for plant species: *A. thaliana*, At.

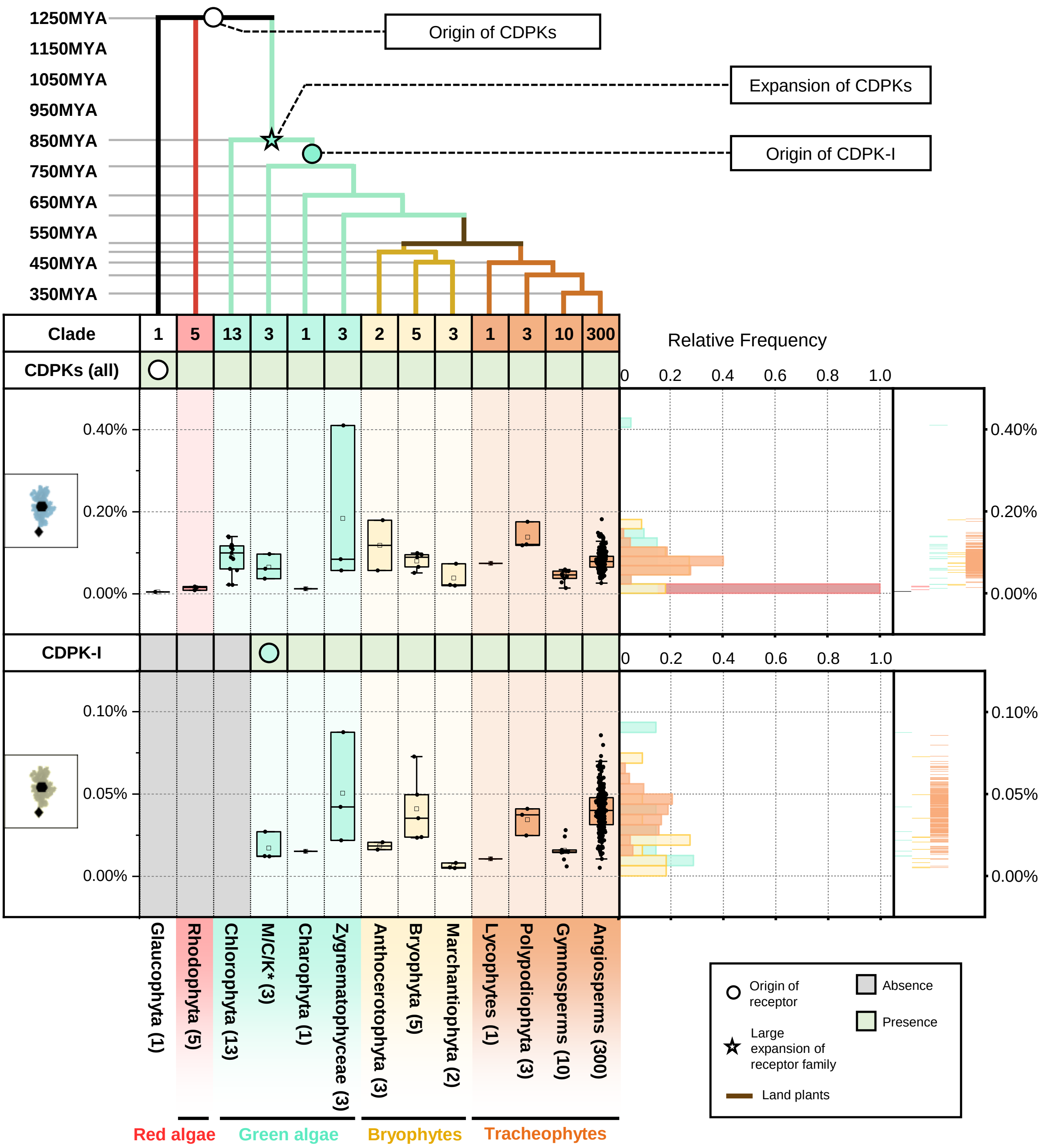


**Supplementary figure 9. The origin and expansion of CDPK in plants.** Top panel represents the phylogenetic tree of multiple algae and plant lineage. With circle (○) and star (☆) indicate the origin and expansion of receptor families. The timescale (in million year; MYA) of the phylogenetic tree is estimated by TIMETREE. Bottom panel represent the present/absence of CDPK (all) and CDPK-I in different algae and plant lineages. Grey box indicates the absence of receptor and green box indicates the presence of receptors in each lineage. The origin of CDPKs are marked with a circle. *M/C/K represents Mesostigmatophyceae, Chlorokybophyceae and Klebsormidiophyceae. Number of available species from each algae and plant lineages are indicated by numbers in the boxes. Boxplot below represents the percentage (%) and right plot represents the distribution and rug of the relative frequency of the % of CDPK (all) and CDPK-I in each lineage.

### MAPKKK phylogenetic tree

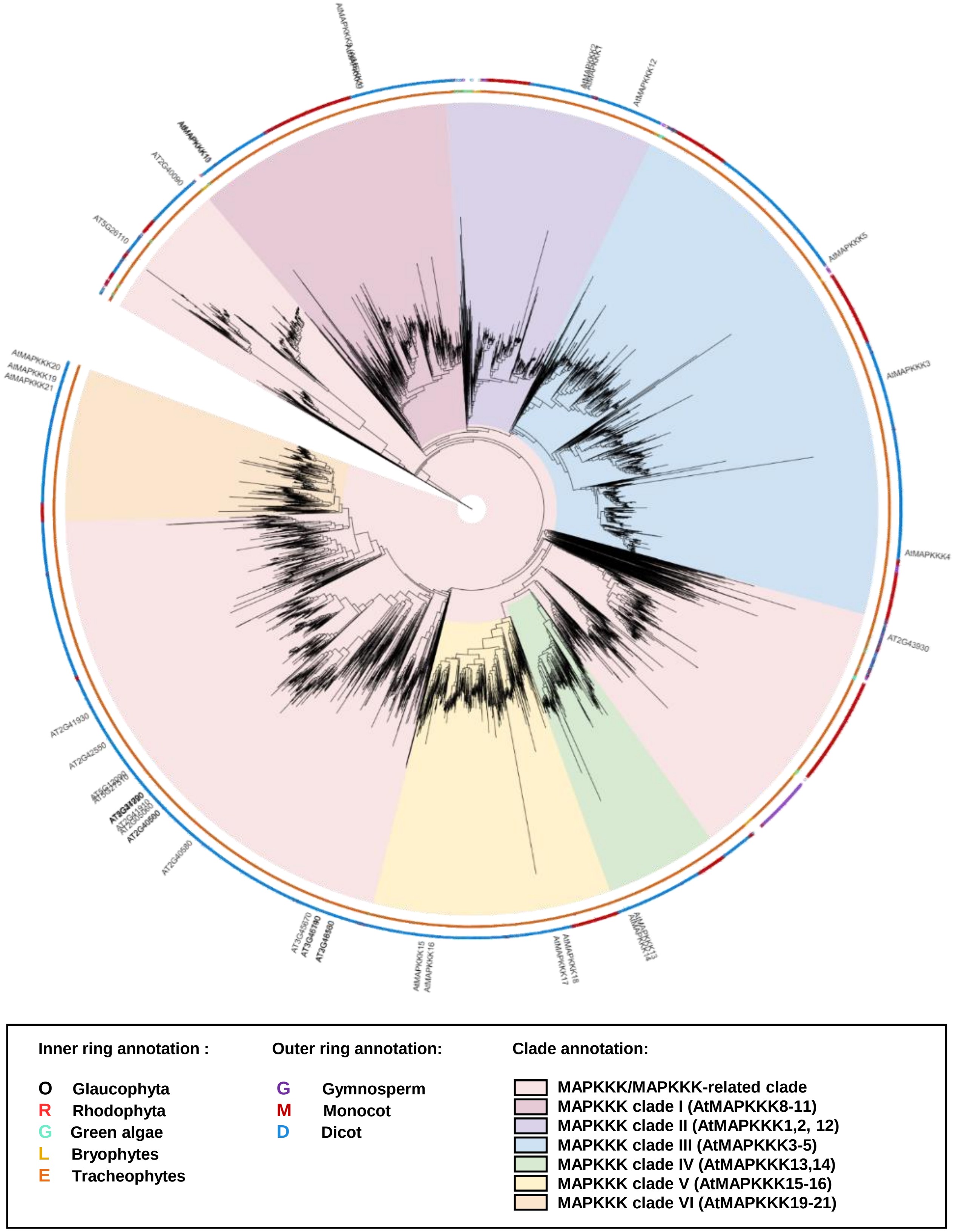


**Supplementary figure 10. Phylogenetic analysis of MAPKKK in plants.** Phylogenetic tree of MAPKKK members identified from 350 species. The inner ring indicates MAPKKK members from either Glaucophyta, red algae (Rhodophyta), green algae, Bryophytes or Tracheophytes. The outer ring indicates MAPKKK members from either gymnosperm, monocots or dicots. The MAPKKK-I/II/III/IV/V/VI clades are defined. *Arabidopsis thaliana* MAPKKKs members are labelled in the tree. Abbreviations for plant species: *A. thaliana*, At.

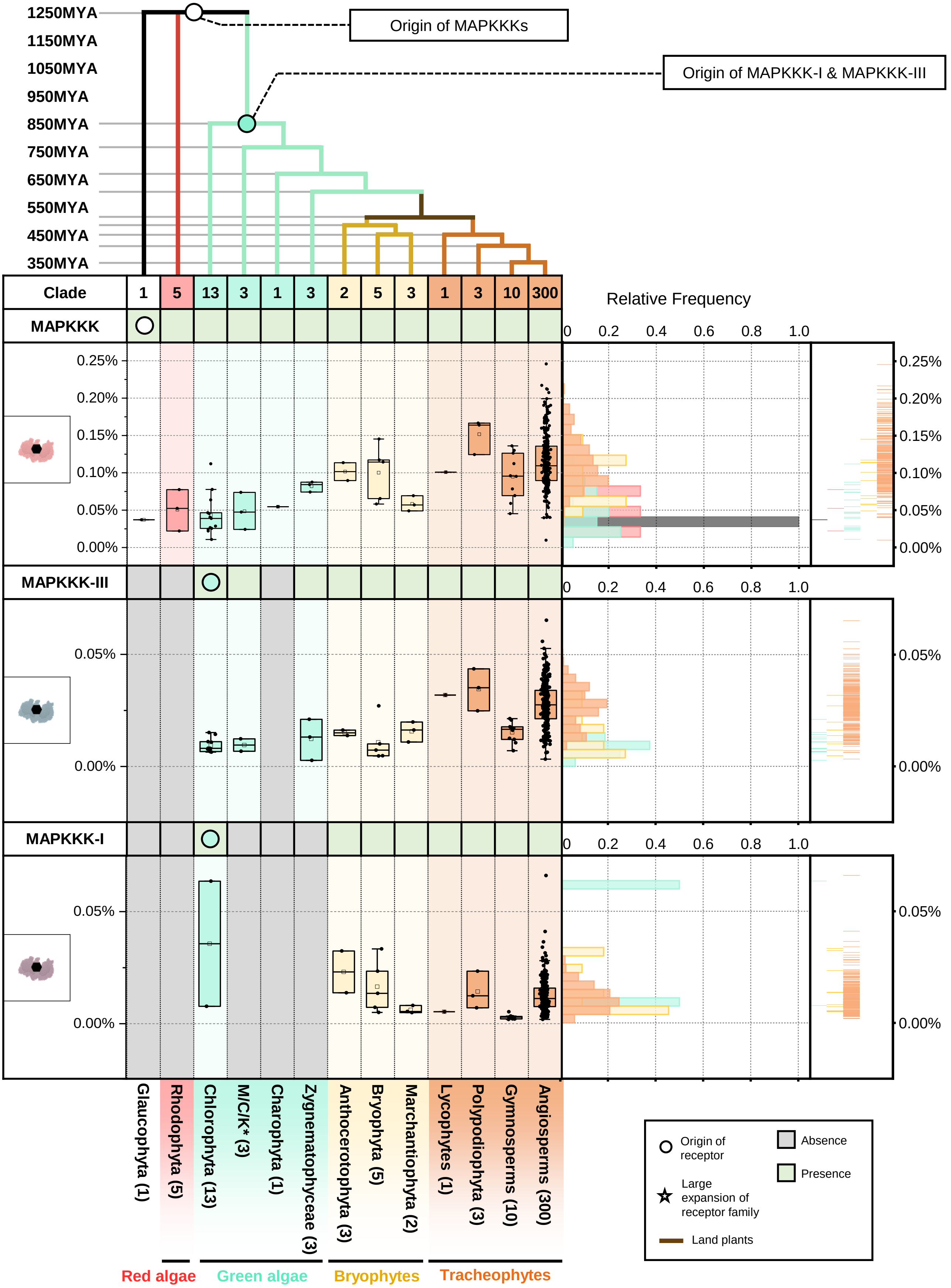


**Supplementary figure 11. The origin and expansion of MAPKKK in plants.** Top panel represents the phylogenetic tree of multiple algae and plant lineage. With circle (○) and star (☆) indicate the origin and expansion of receptor families. The timescale (in million year; MYA) of the phylogenetic tree is estimated by TIMETREE. Bottom panel represent the present/absence of MAPKKK, MAPKKK-III and MAPKKK-I members in different algae and plant lineages. Grey box indicates the absence of receptor and green box indicates the presence of receptors in each lineage. The origin of MAPKKKs are marked with a circle. *M/C/K represents Mesostigmatophyceae, Chlorokybophyceae and Klebsormidiophyceae. Number of available species from each algae and plant lineages are indicated by numbers in the boxes. Boxplot below represents the percentage (%) and right plot represents the distribution and rug of the relative frequency of the % of MAPKKK, MAPKKK-III and MAPKKK-I members in each lineage.

### MAPKK phylogenetic tree

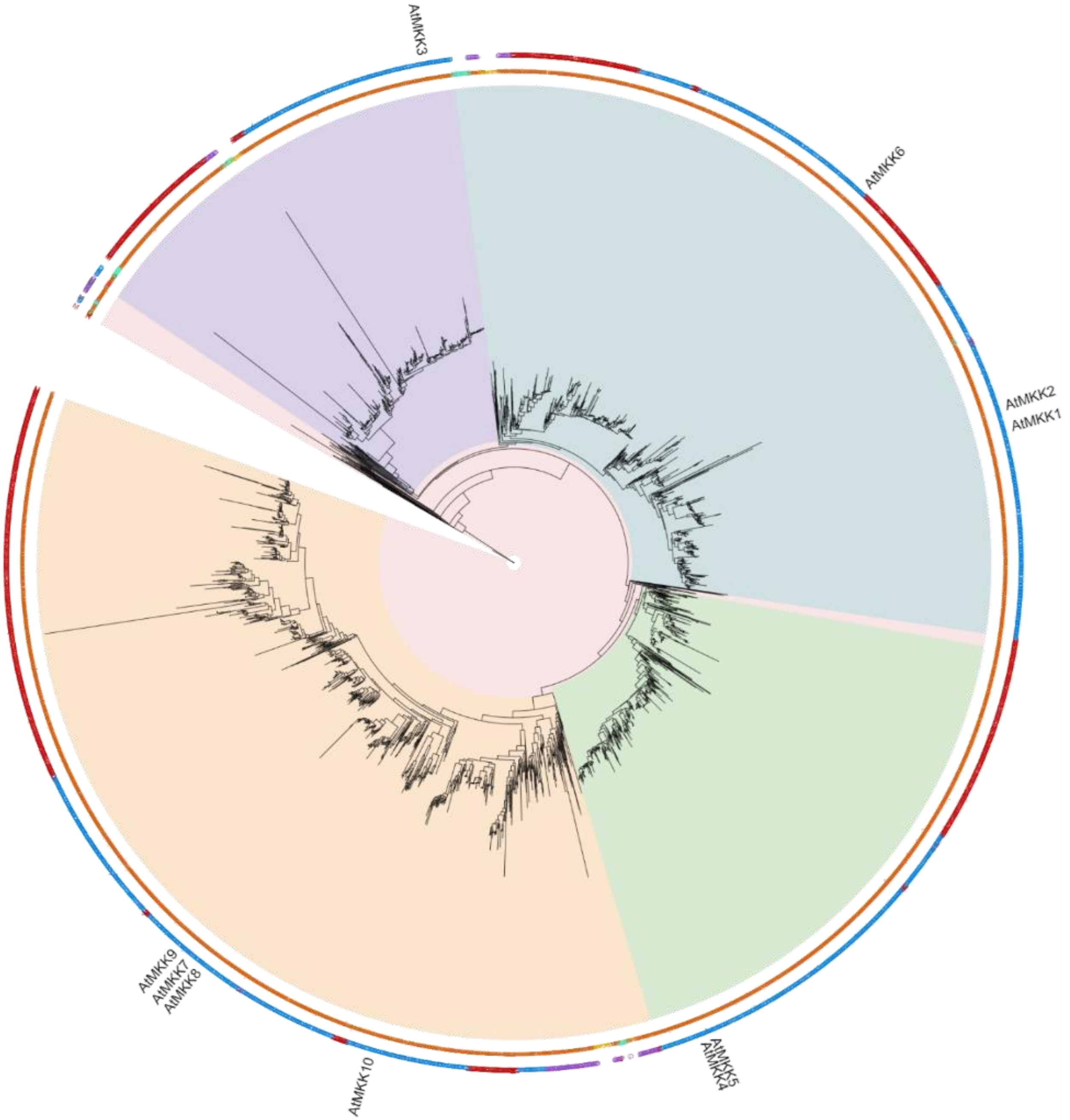


**Inner ring annotation :**

- **O** **Glaucophyta**
- **R** **Rhodophyta**
- **G** **Green algae**
- **L** **Bryophytes**
- **E** **Tracheophytes**

**Outer ring annotation:**

- **G** **Gymnosperm**
- **M** **Monocot**
- **D** **Dicot**

**Clade annotation:**

- **MAPKK/MAPKK-related clade**
- **MAPKK clade I (AtMKK3)**
- **MAPKK clade II (AtMKK1,2,6)**
- **MAPKK clade III (AtMKK4,5)**
- **MAPKK clade IV (AtMKK7-10)**

**Supplementary figure 12. Phylogenetic analysis of MAPKK in plants.** Phylogenetic tree of MAPKK members identified from 350 species. The inner ring indicates MAPKK members from either Glaucophyta, red algae (Rhodophyta), green algae, Bryophytes or Tracheophytes. The outer ring indicates MAPKK members from either gymnosperm, monocots or dicots. The MAPKK-I/II/III/IV clades are defined based on previous annotation[104]. *Arabidopsis thaliana* MAPKKs members are labelled in the tree. Abbreviations for plant species: *A. thaliana*, At.

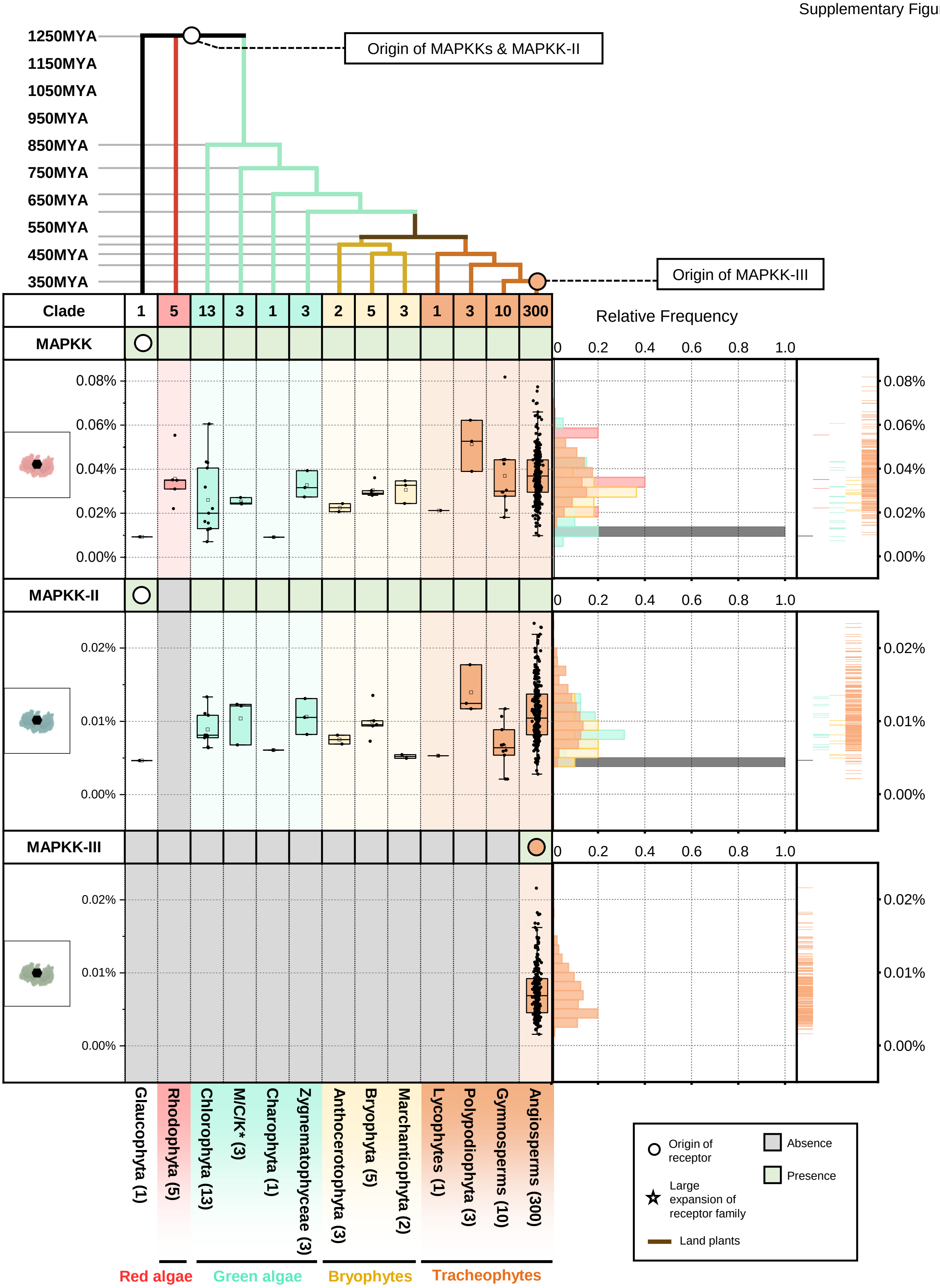


**Supplementary figure 13. The origin and expansion of MAPKK in plants.** Top panel represents the phylogenetic tree of multiple algae and plant lineage. With circle (○) and star (☆) indicate the origin and expansion of receptor families. The timescale (in million year; MYA) of the phylogenetic tree is estimated by TIMETREE. Bottom panel represent the present/absence of MAPKK, MAPKK-II and MAPKK-III members in different algae and plant lineages. Grey box indicates the absence of receptor and green box indicates the presence of receptors in each lineage. The origin of MAPKKs are marked with a circle. *M/C/K represents Mesostigmatophyceae, Chlorokybophyceae and Klebsormidiophyceae. Number of available species from each algae and plant lineages are indicated by numbers in the boxes. Boxplot below represents the percentage (%) and right plot represents the distribution and rug of the relative frequency of the % of MAPKK, MAPKK-II and MAPKK-III members in each lineage.

### MAPK phylogenetic tree

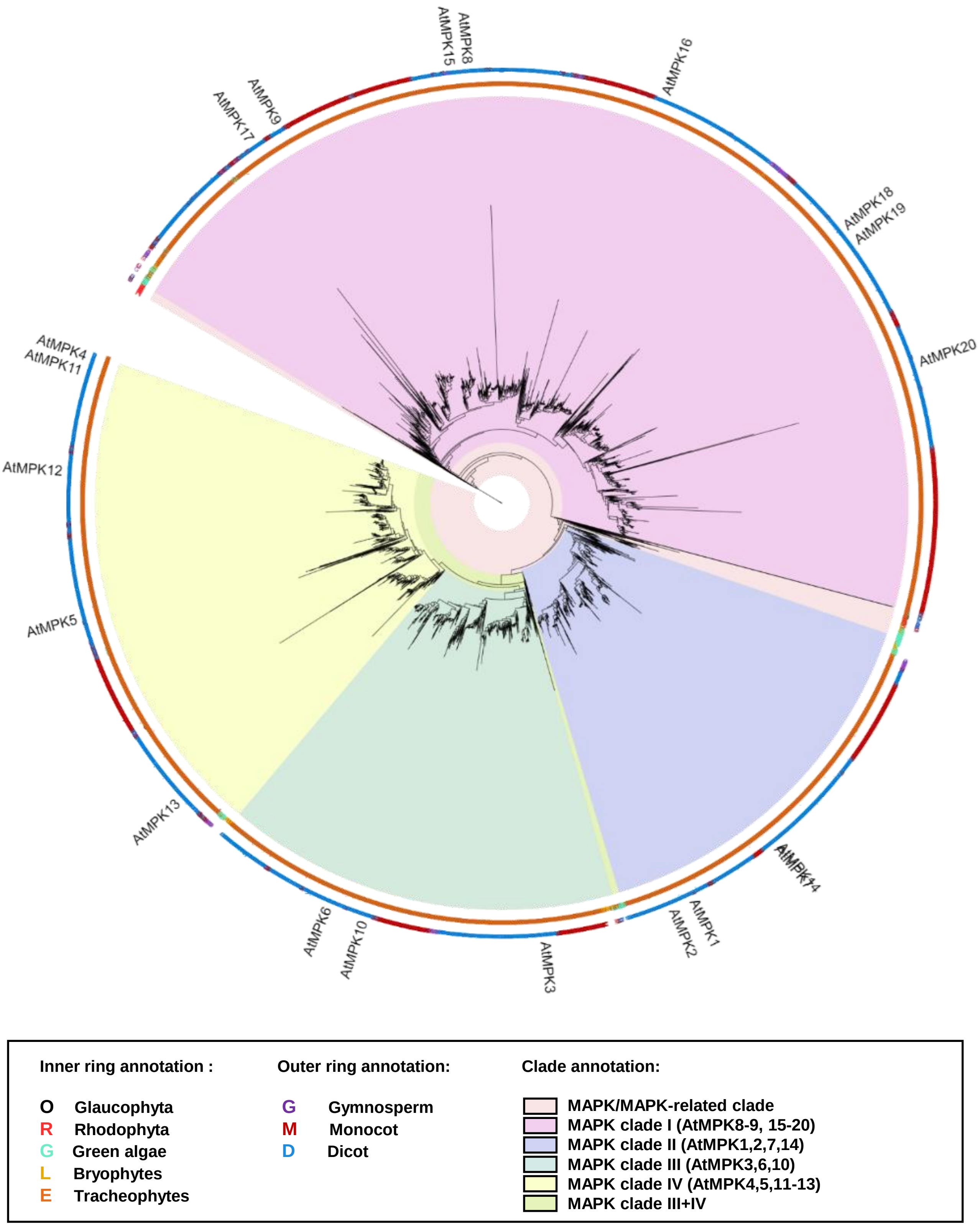


**Supplementary figure 14. Phylogenetic analysis of MAPK in plants.** Phylogenetic tree of MAPK members identified from 350 species. The inner ring indicates MAPK members from either Glaucophyta, red algae (Rhodophyta), green algae, Bryophytes or Tracheophytes. The outer ring indicates MAPK members from either gymnosperm, monocots or dicots. The MAPK-I/II/III/IV clades are defined based on previous annotation[104,105]. *Arabidopsis thaliana* MAPKs members are labelled in the tree. Abbreviations for plant species: *A. thaliana*, At.

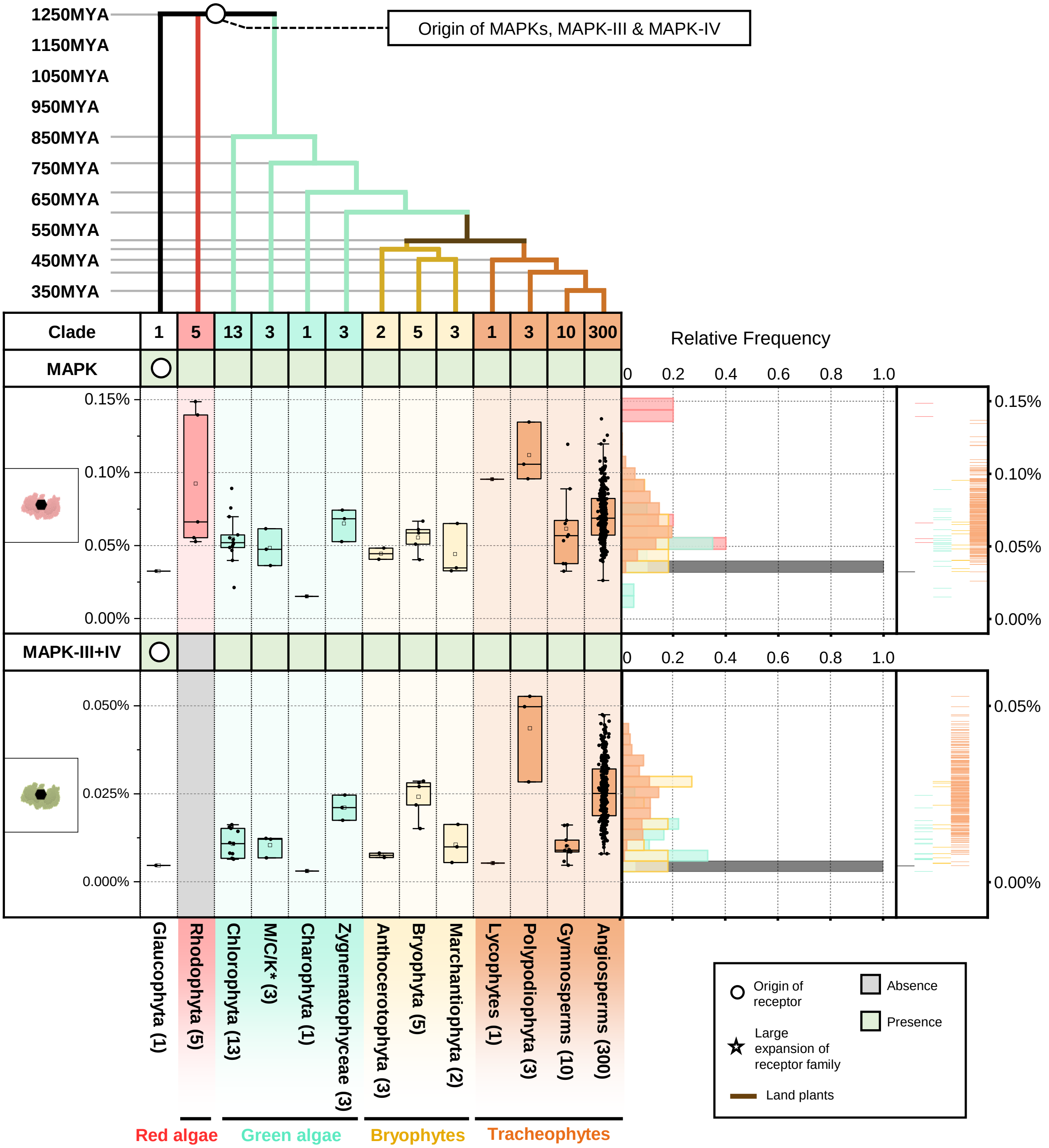


**Supplementary figure 15. The origin and expansion of MAPK in plants.** Top panel represents the phylogenetic tree of multiple algae and plant lineage. With circle (○) and star (☆) indicate the origin and expansion of receptor families. The timescale (in million year; MYA) of the phylogenetic tree is estimated by TIMETREE. Bottom panel represent the present/absence of MAPK, MAPK-III+IV members in different algae and plant lineages. Grey box indicates the absence of receptor and green box indicates the presence of receptors in each lineage. The origin of MAPKs are marked with a circle. *M/C/K represents Mesostigmatophyceae, Chlorokybophyceae and Klebsormidiophyceae. Number of available species from each algae and plant lineages are indicated by numbers in the boxes. Boxplot below represents the percentage (%) and right plot represents the distribution and rug of the relative frequency of the % of MAPK, MAPK-III+IV members in each lineage.

### CNGC phylogenetic tree

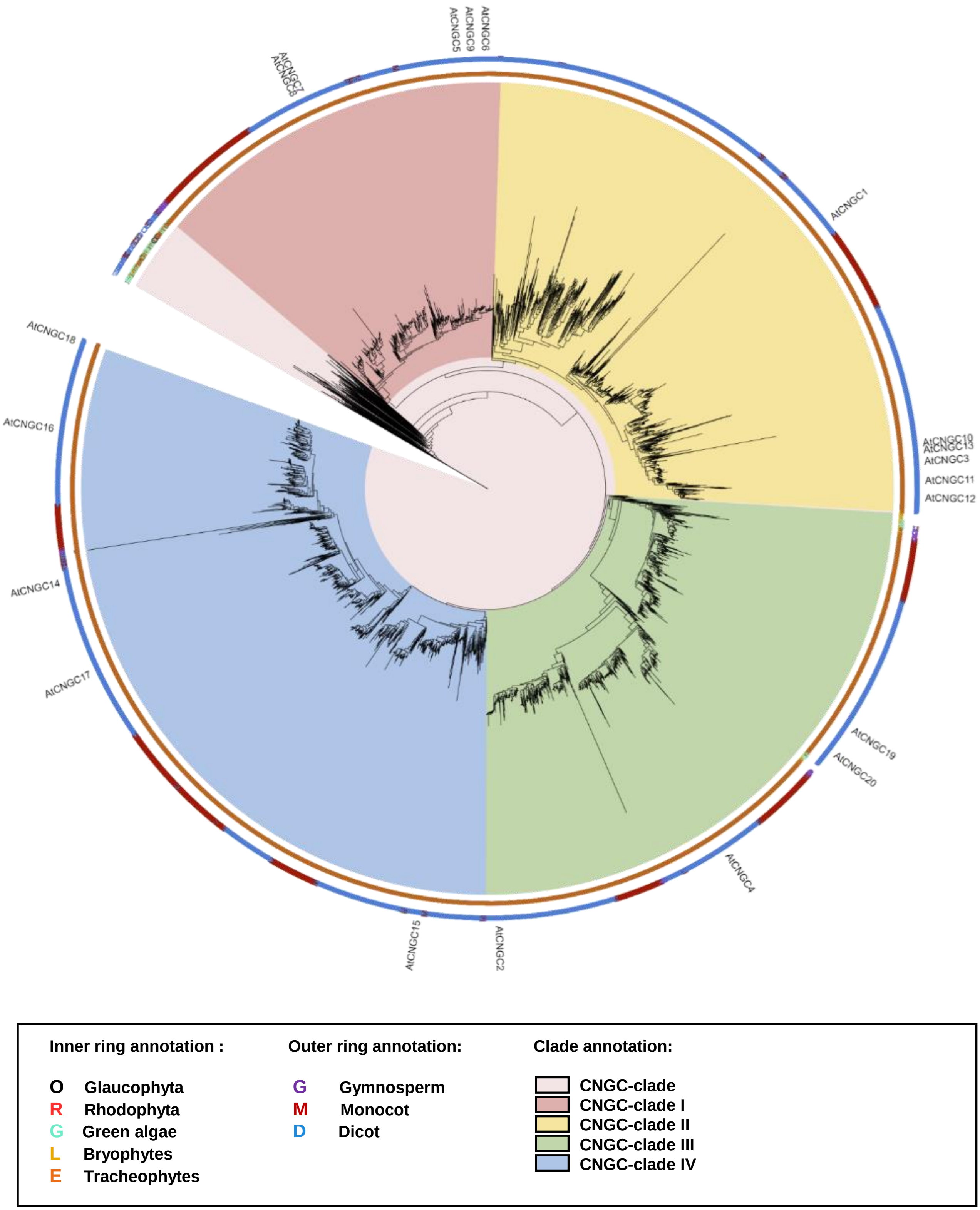


**Supplementary figure 16. Phylogenetic analysis of CNGCs in plants.** Phylogenetic tree of CNGCs identified from 350 species. The inner ring indicates CNGC members from either Glaucophyta, red algae (Rhodophyta), green algae, Bryophytes or Tracheophytes. The outer ring indicates CNGC members from either gymnosperm, monocots or dicots. The CNGC-I/II/III/IV clades are defined based on previous annotation[106]. *Arabidopsis thaliana* CNGC members are labelled in the tree. Abbreviations for plant species: *A. thaliana*, At.

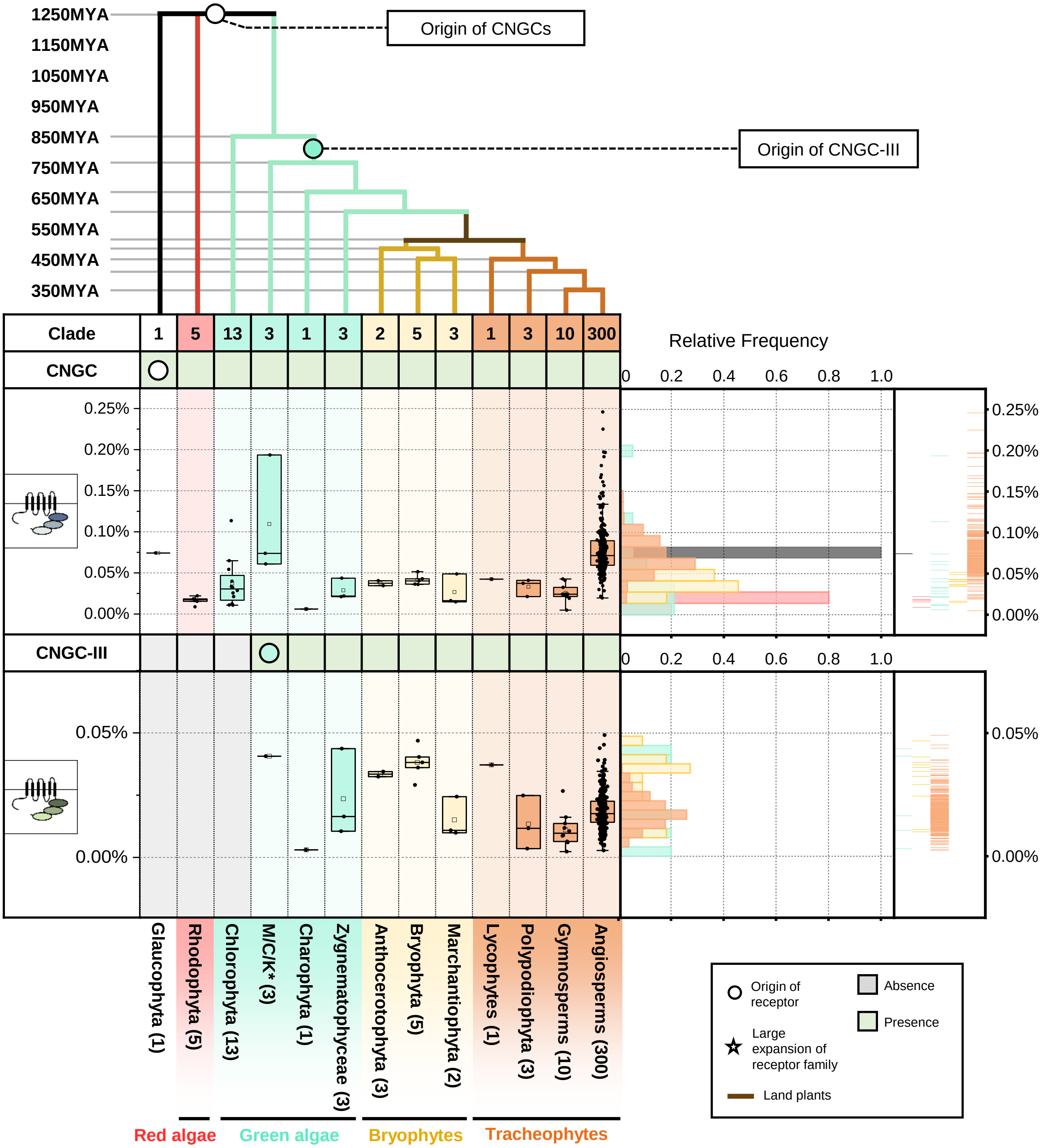


**Supplementary figure 17. The origin and expansion of CNGC in plants.** Top panel represents the phylogenetic tree of multiple algae and plant lineage. With circle (○) and star (☆) indicate the origin and expansion of receptor families. The timescale (in million year; MYA) of the phylogenetic tree is estimated by TIMETREE. Bottom panel represent the present/absence of CNGC and CNGC-I members in different algae and plant lineages. Grey box indicates the absence of receptor and green box indicates the presence of receptors in each lineage. The origin of CNGCs are marked with a circle. *M/C/K represents Mesostigmatophyceae, Chlorokybophyceae and Klebsormidiophyceae. Number of available species from each algae and plant lineages are indicated by numbers in the boxes. Boxplot below represents the percentage (%) and right plot represents the distribution and rug of the relative frequency of the % of CNGC and CNGC-I members in each lineage.

### OSCA phylogenetic tree

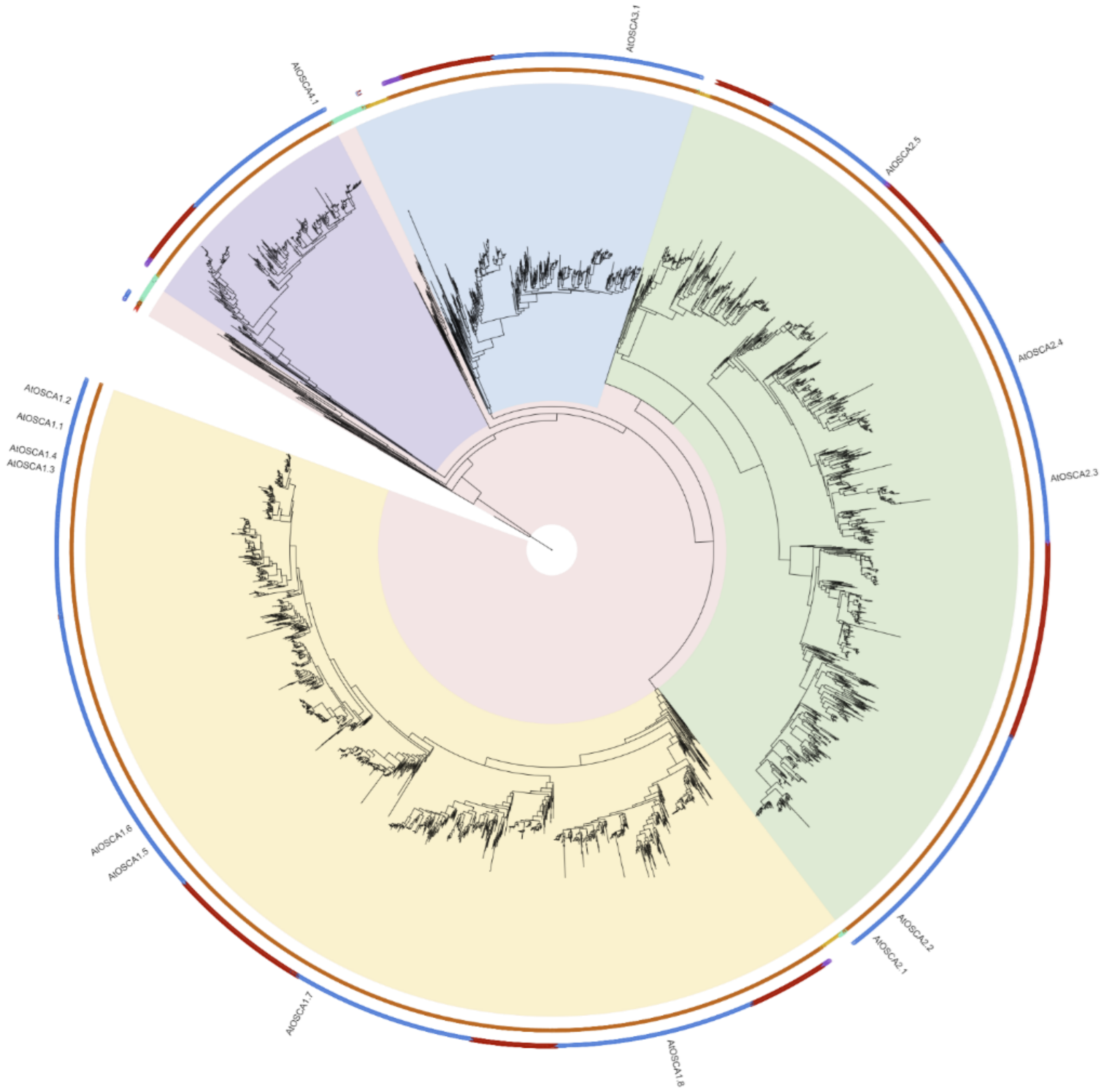


**Inner ring annotation :**

- **O** **Glaucophyta**
- **R** **Rhodophyta**
- **G** **Green algae**
- **L** **Bryophytes**
- **E** **Tracheophytes**

**Outer ring annotation:**

- **G** **Gymnosperm**
- **M** **Monocot**
- **D** **Dicot**

**Clade annotation:**

- **OSCA-clade**
- **OSCA1 clade**
- **OSCA2 clade**
- **OSCA3 clade**
- **OSCA4 clade**

**Supplementary figure 18. Phylogenetic analysis of OSCA in plants.** Phylogenetic tree of OSCA members identified from 350 species. The inner ring indicates OSCA members from either Glaucophyta, red algae (Rhodophyta), green algae, Bryophytes or Tracheophytes. The outer ring indicates OSCA members from either gymnosperm, monocots or dicots. The OSCA1/2/3/4 clades are defined based on previous annotation[107]. *Arabidopsis thaliana* OSCAs members are labelled in the tree. Abbreviations for plant species: *A. thaliana*, At.

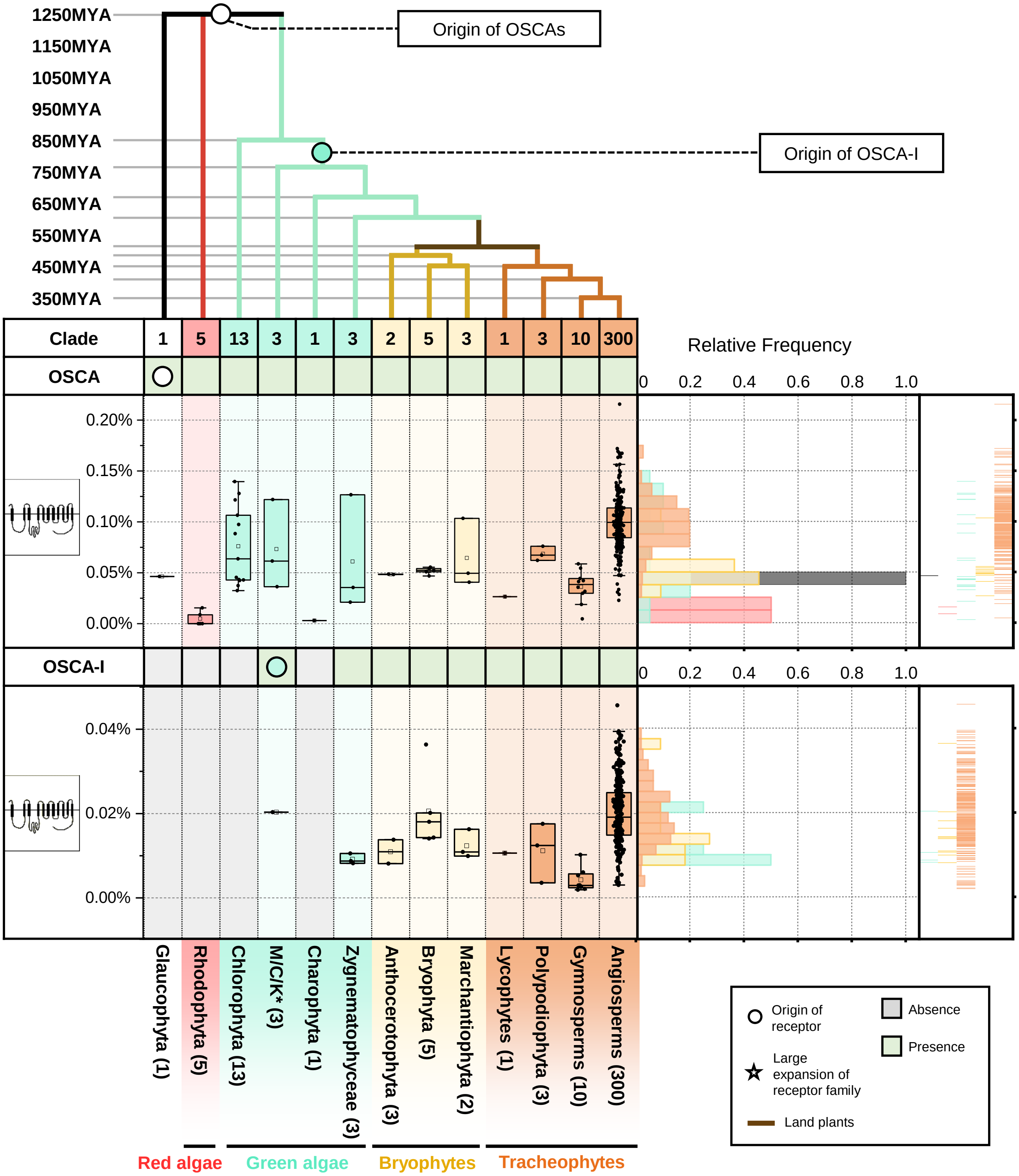


**Supplementary figure 19. The origin and expansion of OSCA in plants.** Top panel represents the phylogenetic tree of multiple algae and plant lineage. With circle (○) and star (☆) indicate the origin and expansion of receptor families. The timescale (in million year; MYA) of the phylogenetic tree is estimated by TIMETREE. Bottom panel represent the present/absence of OSCA and OSCA-I members in different algae and plant lineages. Grey box indicates the absence of receptor and green box indicates the presence of receptors in each lineage. The origin of OSCAs are marked with a circle. *M/C/K represents Mesostigmatophyceae, Chlorokybophyceae and Klebsormidiophyceae. Number of available species from each algae and plant lineages are indicated by numbers in the boxes. Boxplot below represents the percentage (%) and right plot represents the distribution and rug of the relative frequency of the % of OSCA and OSCA-I members in each lineage.

### NADPH oxidases phylogenetic tree

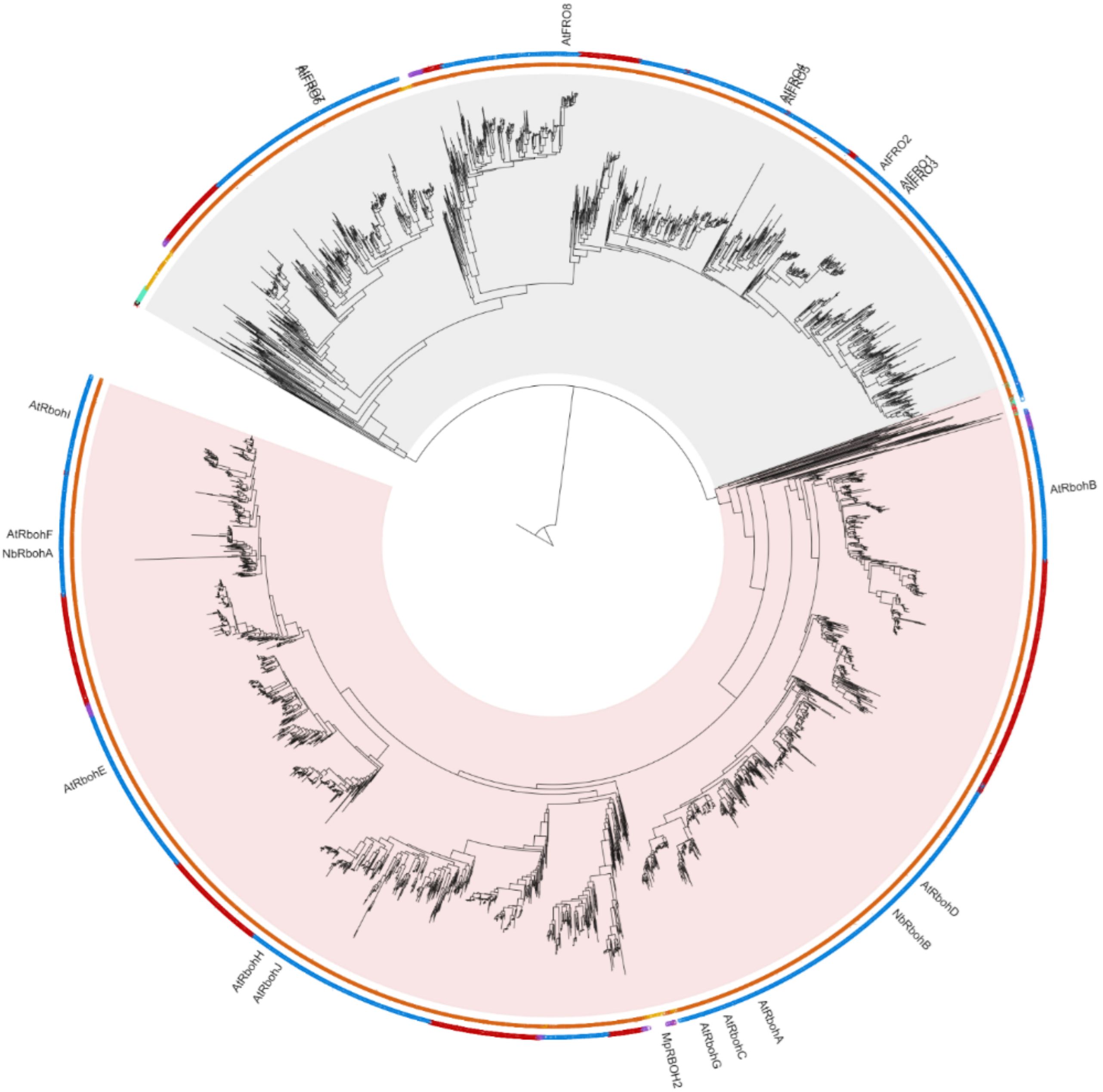


**Inner ring annotation :**

- **O** Glaucophyta
- **R** Rhodophyta
- **G** Green algae
- **L** Bryophytes
- **E** Tracheophytes

**Outer ring annotation:**

- **G** Gymnosperm
- **M** Monocot
- **D** Dicot

**Clade annotation:**

- RBOH clade
- FRO clade

**Supplementary figure 20. Phylogenetic analysis of NADPH oxidases in plants.** Phylogenetic tree of NADPH oxidase members identified from 350 species. The inner ring indicates NADPH oxidase members from either Glaucophyta, red algae (Rhodophyta), green algae, Bryophytes or Tracheophytes. The outer ring indicates NADPH oxidase members from either gymnosperm, monocots or dicots. The FRO and RBOH clades are defined. Characterized RBOH members and *Arabidopsis thaliana* NADPH oxidases are labelled in the tree. Abbreviations for plant species: *M. polymorpha*, Mp; *A. thaliana*, At; *N. benthamiana*, Nb.

### RBOH phylogenetic tree

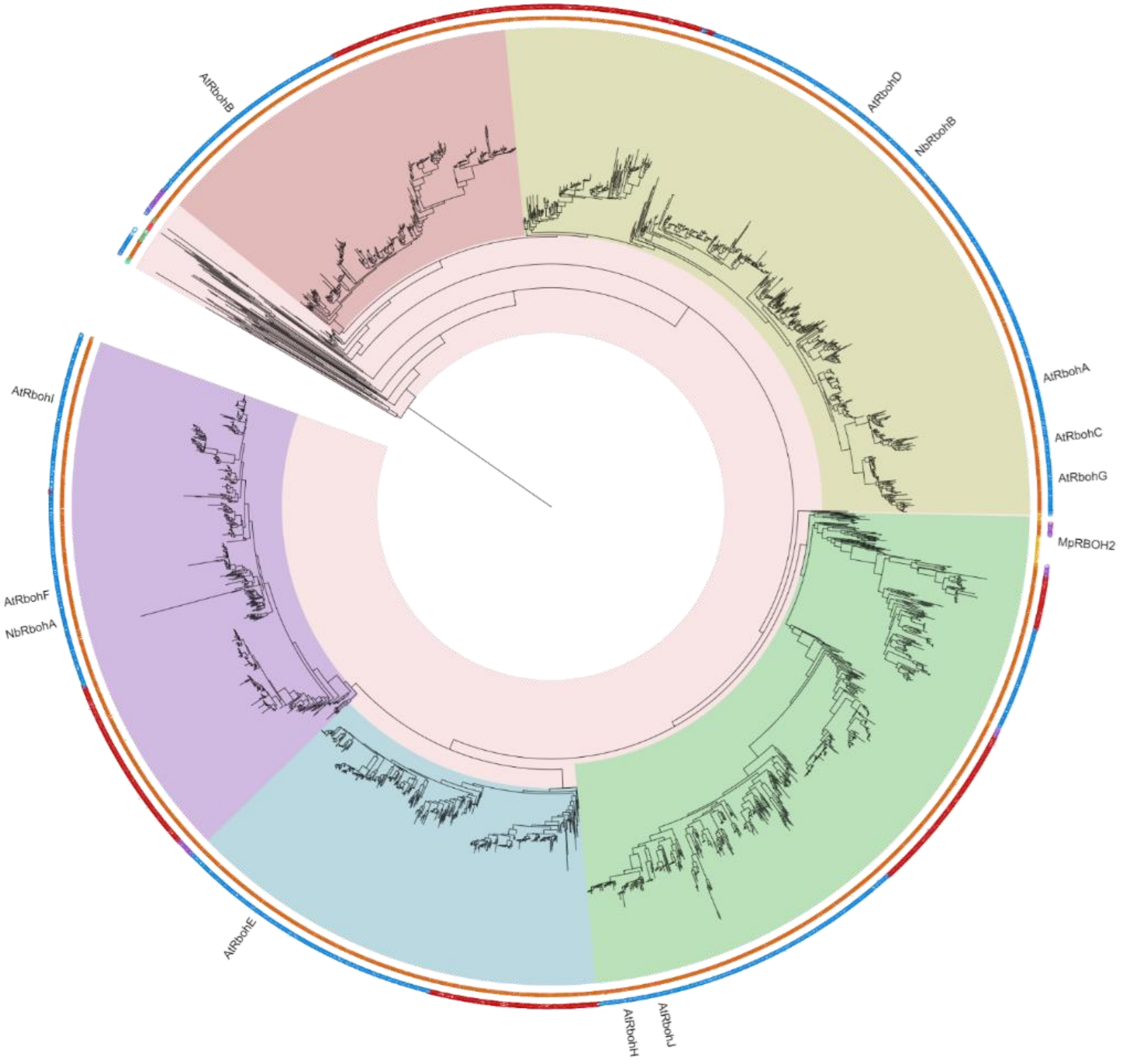


**Inner ring annotation :**

- O Glaucophyta
- R Rhodophyta
- G Green algae
- L Bryophytes
- E Tracheophytes

**Outer ring annotation:**

- G Gymnosperm
- M Monocot
- D Dicot

**Clade annotation:**

- RBOH clade
- RBOH clade-I
- RBOH clade-II
- RBOH clade-III
- RBOH clade-IV
- RBOH clade-V

**Supplementary figure 21. Phylogenetic analysis of RBOHs in plants.** Phylogenetic tree of RBOH members identified from 350 species. The inner ring indicates RBOH members from either Glaucophyta, red algae (Rhodophyta), green algae, Bryophytes or Tracheophytes. The outer ring indicates RBOH members from either gymnosperm, monocots or dicots. The RBOH-I/II/III/IV/V clades are defined based on previous annotation[108,109]. *Arabidopsis thaliana* RBOH members are labelled in the tree. Abbreviations for plant species: *M. polymorpha*, Mp; *A. thaliana*, At; *N. benthamiana*, Nb.

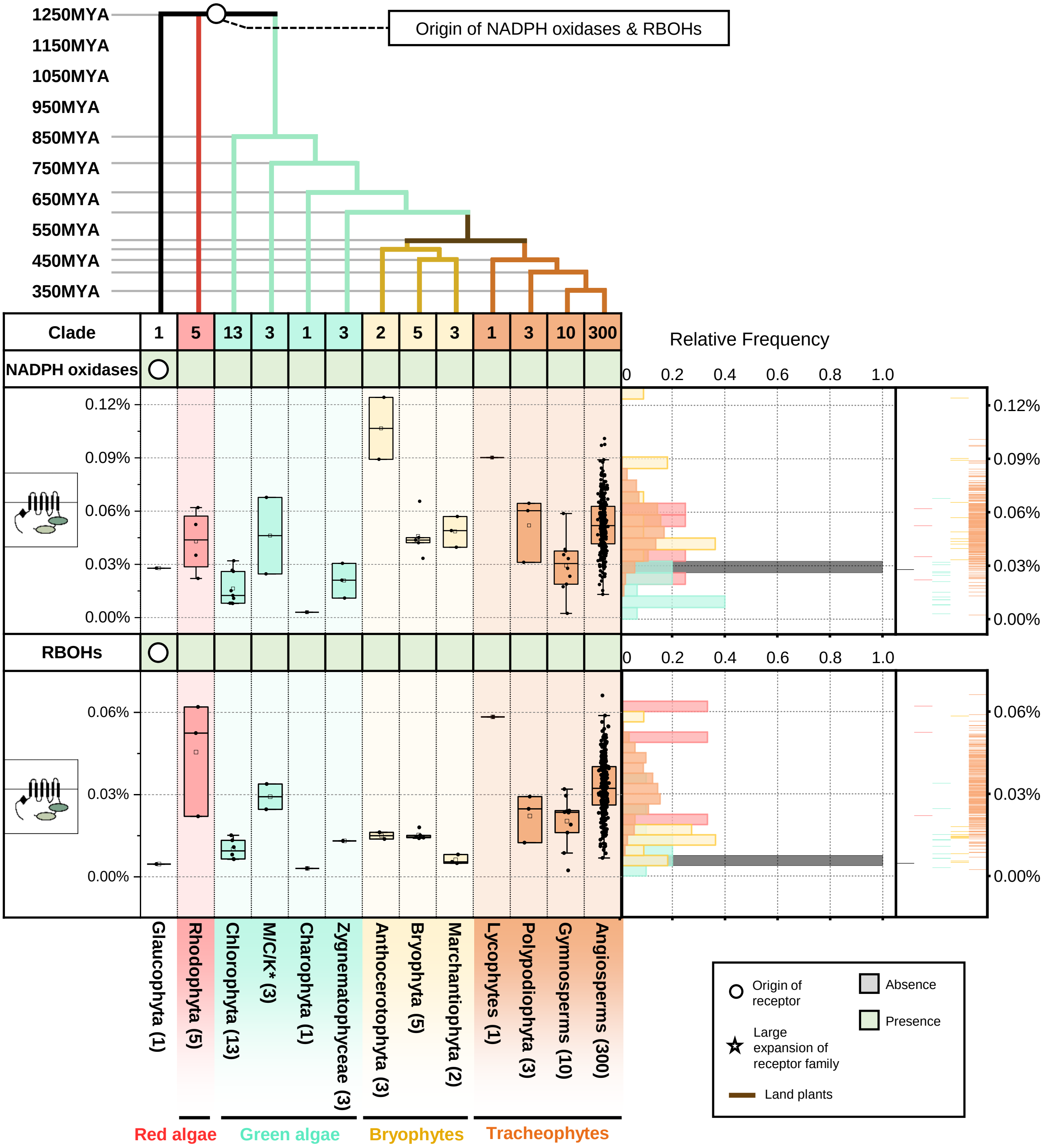


**Supplementary figure 22. The origin and expansion of NADPH oxidases in plants.** Top panel represents the phylogenetic tree of multiple algae and plant lineage. With circle (○) and star (☆) indicate the origin and expansion of receptor families. The timescale (in million year; MYA) of the phylogenetic tree is estimated by TIMETREE. Bottom panel represent the present/absence of NADPH oxidase and RBOHs in different algae and plant lineages. Grey box indicates the absence of receptor and green box indicates the presence of receptors in each lineage. The origin of NADPH oxidases are marked with a circle. *M/C/K represents Mesostigmatophyceae, Chlorokybophyceae and Klebsormidiophyceae. Number of available species from each algae and plant lineages are indicated by numbers in the boxes. Boxplot below represents the percentage (%) and right plot represents the distribution and rug of the relative frequency of the % of NADPH oxidase and RBOHs in each lineage.

### EP proteins phylogenetic tree

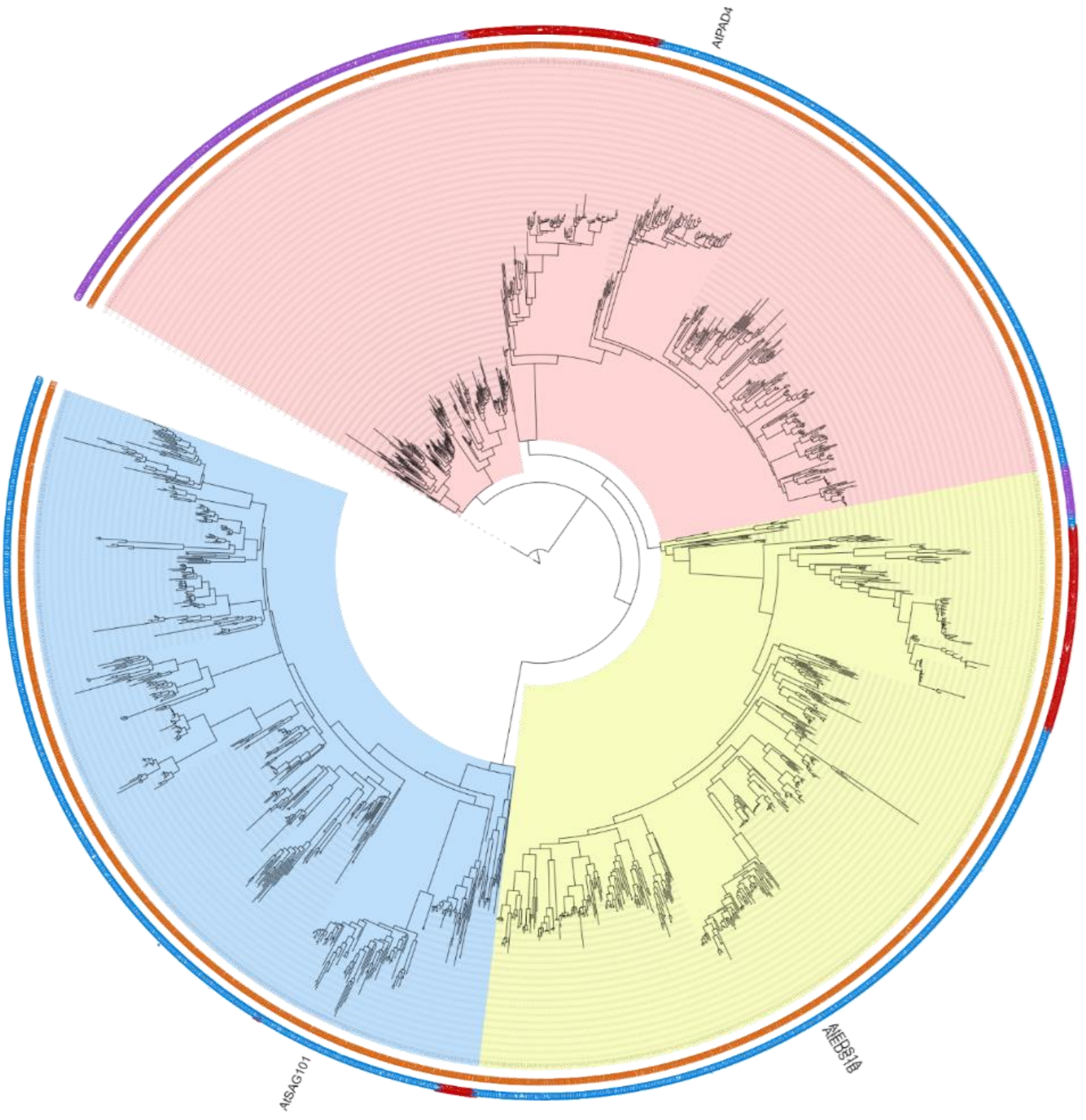


**Inner ring annotation :**

- **O** **Glaucophyta**
- **R** **Rhodophyta**
- **G** **Green algae**
- **L** **Bryophytes**
- **E** **Tracheophytes**

**Outer ring annotation:**

- **G** **Gymnosperm**
- **M** **Monocot**
- **D** **Dicot**

**Clade annotation:**

- **PAD4 clade**
- **EDS1 clade**
- **SAG101 clade**

**Supplementary figure 23. Phylogenetic analysis of EP proteins in plants.** Phylogenetic tree of EP proteins identified from 350 species. The inner ring indicates EP proteins from either Glaucophyta, red algae (Rhodophyta), green algae, Bryophytes or Tracheophytes. The outer ring indicates EP proteins from either gymnosperm, monocots or dicots. The PAD4, EDS1 and SAG101 clades are defined based on previous annotation[110]. *Arabidopsis thaliana* EP protein members are labelled in the tree. Abbreviations for plant species: *A. thaliana*, At.

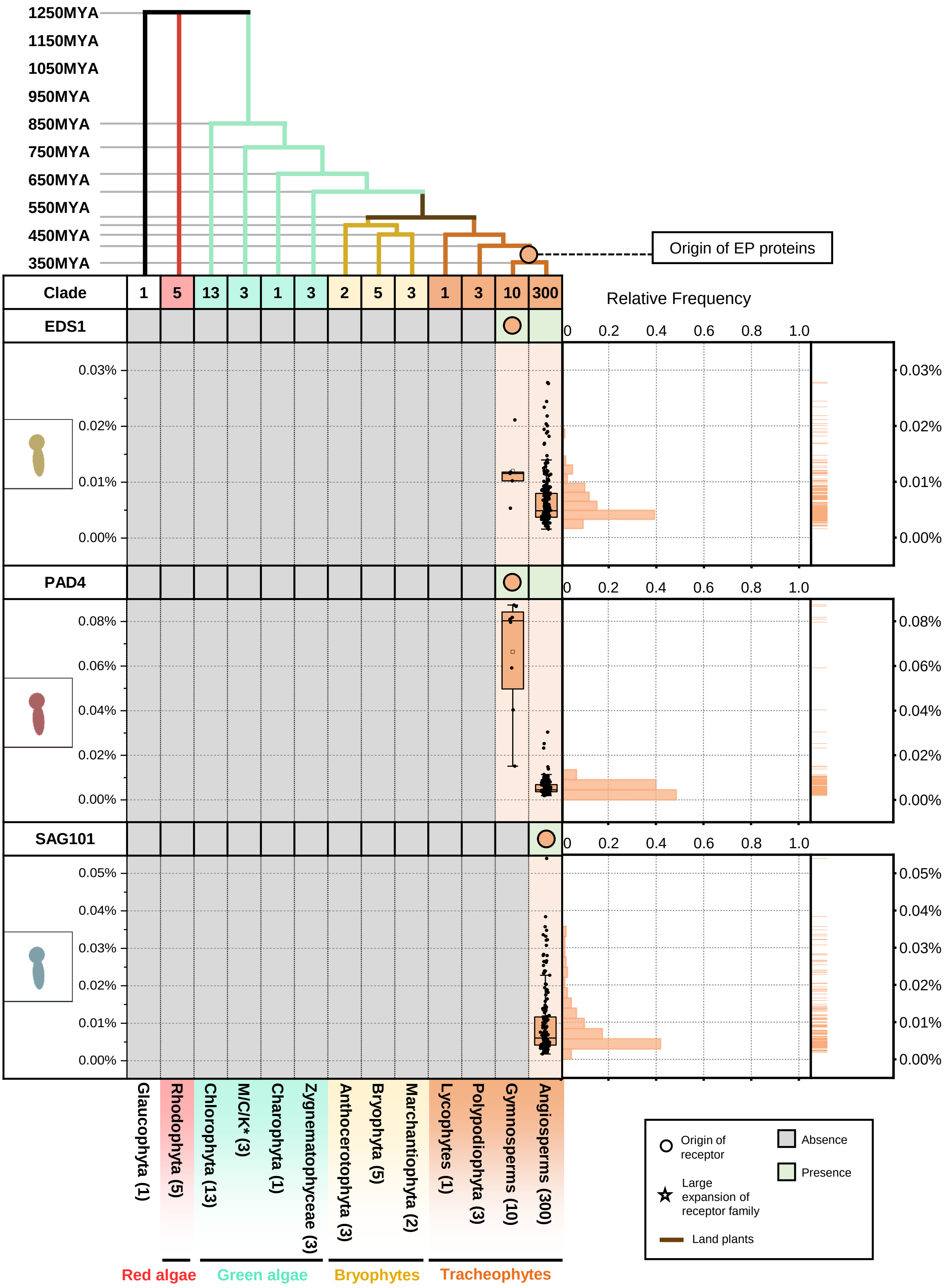


**Supplementary figure 24. The origin and expansion of EP proteins in plants.** Top panel represents a phylogenetic tree of multiple algae and plant lineages. Circles (○) and stars (☆) indicate the origin and expansion of receptor families. The timescale (in million years; MYA) of the phylogenetic tree is estimated by TIMETREE. Bottom panel represent the present/absence of EDS1, PAD4 and SAG101 in different algae and plant lineages. Grey box indicates the absence of receptor and green box indicates the presence of EDS1, PAD4 and SAG101 in each lineage. The origin of EP proteins are marked with a circle. *M/C/K represents Mesostigmatophyceae, Chlorokybophyceae and Klebsormidiophyceae. Number of available species from each algae and plant lineages are indicated by numbers in the boxes. Boxplot below represents the percentage (%) and right plot represents the distribution and rug of the relative frequency of the % of EDS1, PAD4 and SAG101 in each lineage.

### RWP8-NLR phylogenetic tree

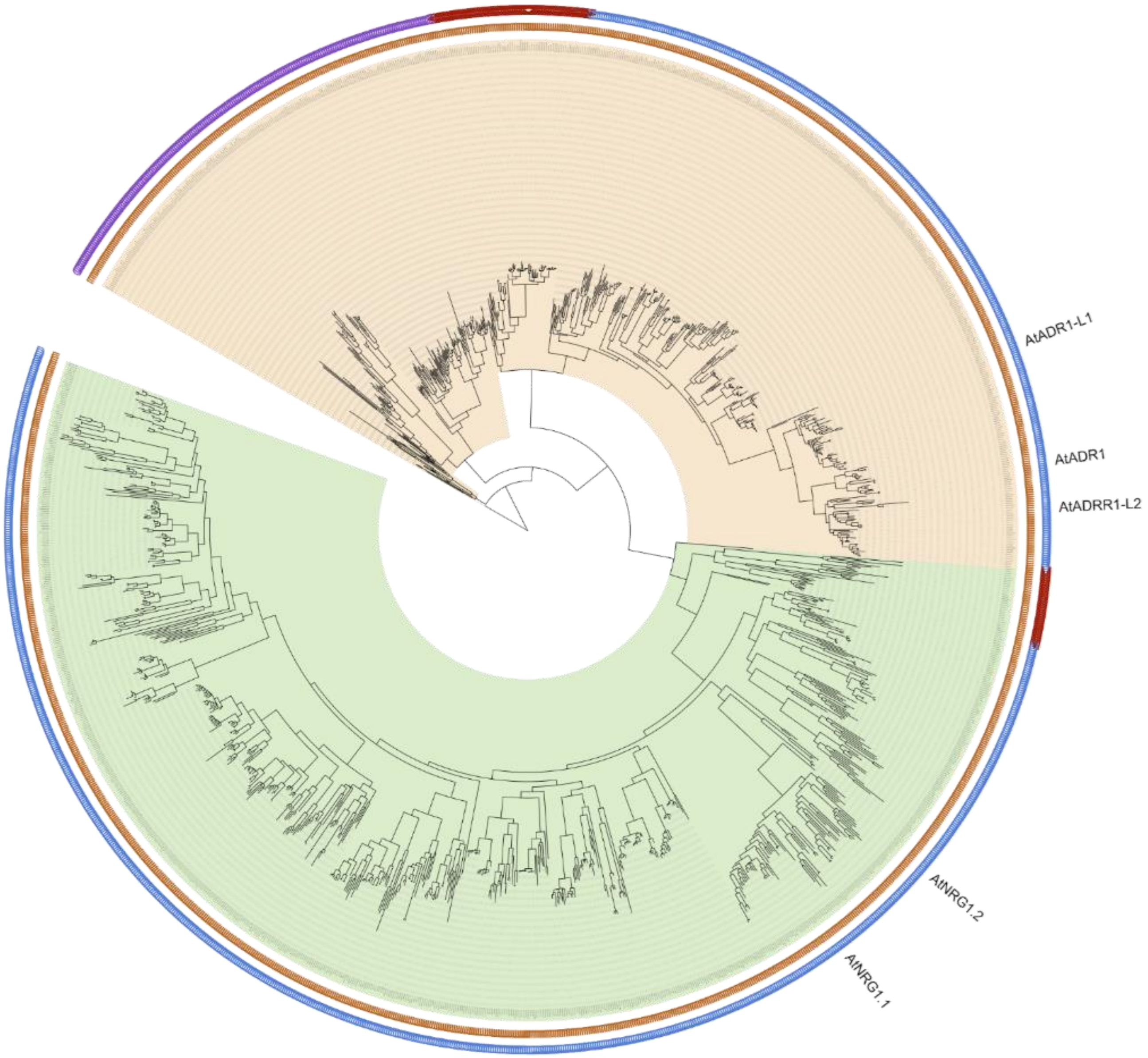


| Inner ring annotation : | | Outer ring annotation: | | Clade annotation: |
|---|---|---|---|---|
| O | Glaucophyta | G | Gymnosperm | ADR1 clade |
| R | Rhodophyta | M | Monocot | NRG1 clade |
| G | Green algae | D | Dicot | |
| L | Bryophytes | | | |
| E | Tracheophytes | | | |

**Supplementary figure 25. Phylogenetic analysis of RPW8-NLRs (RNLs/helper NLRs) in plants.** Phylogenetic tree of helper NLRs identified from 350 species. The inner ring indicates helper NLRs from either Glaucophyta, red algae (Rhodophyta), green algae, Bryophytes or Tracheophytes. The outer ring indicates helper NLRs from either gymnosperm, monocots or dicots. The ADR1 and NRG1 clades are defined based on previous annotations[111]. *Arabidopsis thaliana* helper NLRs are labelled in the tree. Abbreviations for plant species: *A. thaliana*, At.

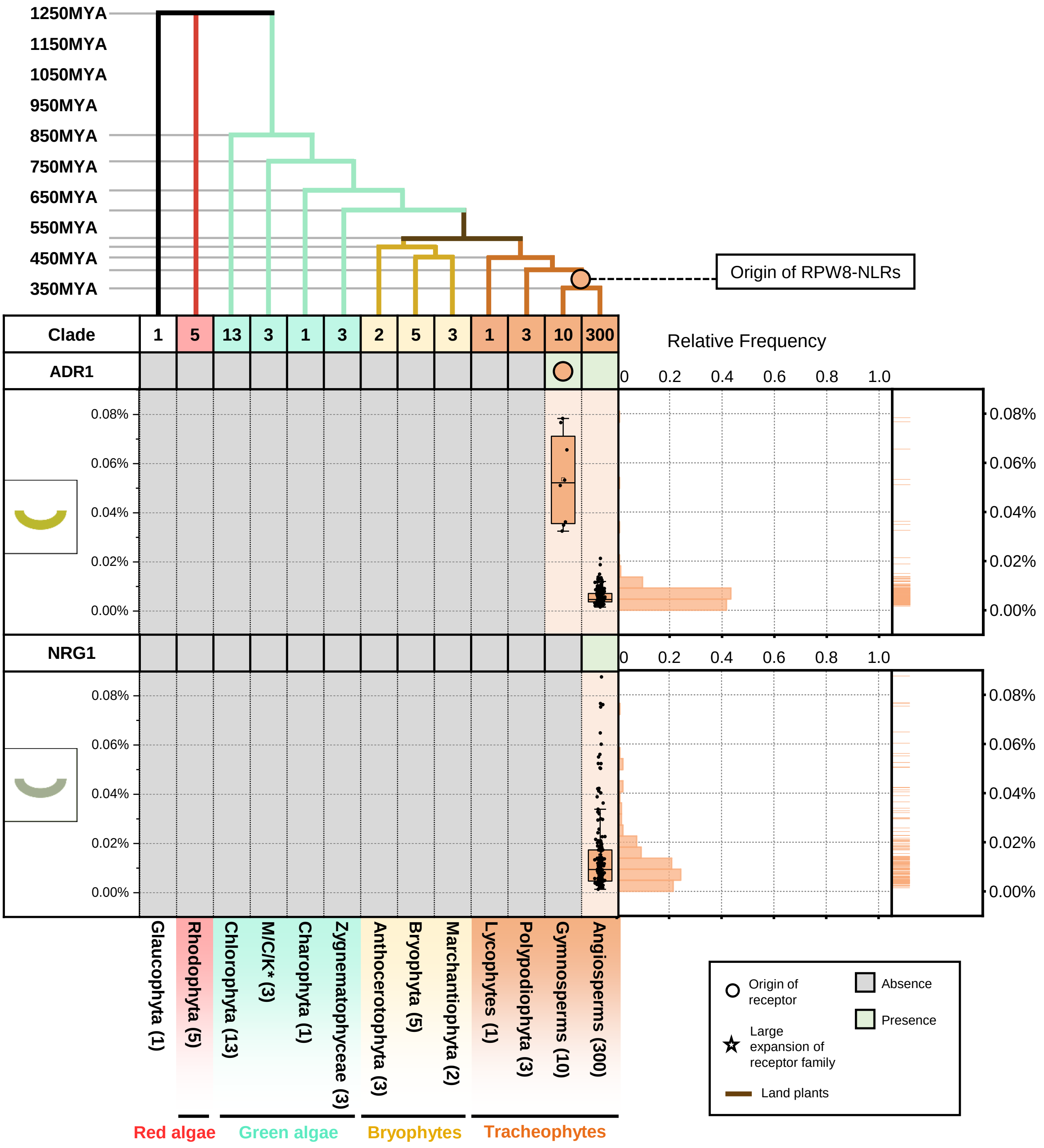


**Supplementary figure 26. The origin and expansion of RWP8-NLRs in plants.** Top panel represents a phylogenetic tree of multiple algae and plant lineages. Circles (○) and stars (☆) indicate the origin and expansion of receptor families, respectively. The timescale (in million years; MYA) of the phylogenetic tree was estimated by TIMETREE. Bottom panel represents the presence or absence of RPW8-NLRs in different algal and plant lineages. Grey box indicates the absence of receptors and green box indicates the presence of receptors in each lineage. The origin of RWP8-NLRs is marked with a circle. *M/C/K represents Mesostigmatophyceae, Chlorokybophyceae and Klebsormidiophyceae. Number of available species from algae and plant lineages are indicated by numbers in the boxes. Boxplot below represents the percentage (%) and right plot represents the distribution and rug of the relative frequency of the % RPW8-NLRs in each lineage.

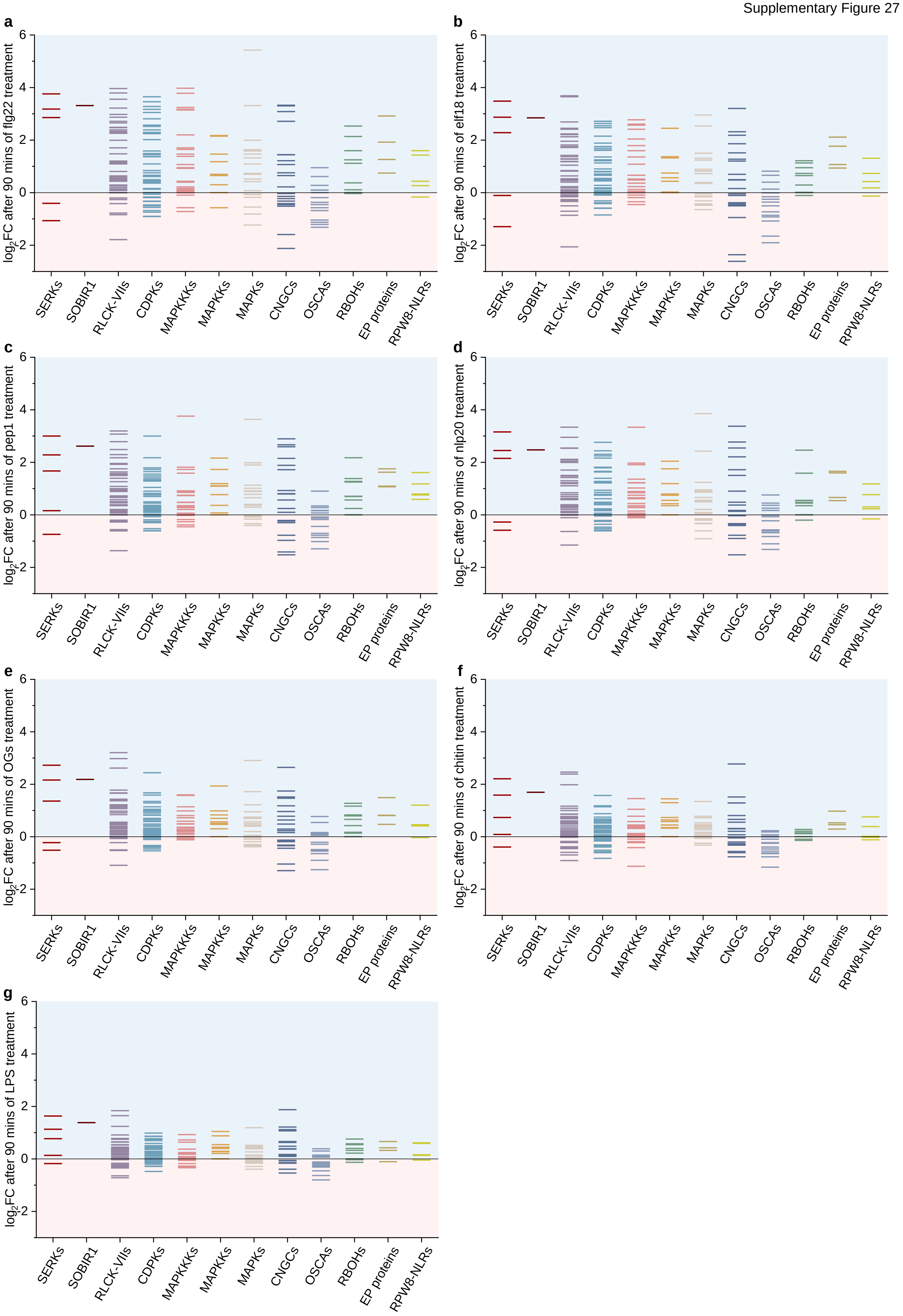
a
b
c
d
e
f
g
log$_2$FC after 90 mins of flg22 treatment
log$_2$FC after 90 mins of elf18 treatment
log$_2$FC after 90 mins of pep1 treatment
log$_2$FC after 90 mins of nlp20 treatment
log$_2$FC after 90 mins of OGs treatment
log$_2$FC after 90 mins of chitin treatment
log$_2$FC after 90 mins of LPS treatment
6
4
2
0
-2
SERKs
SOBIR1
RLCK-VIIs
CDPKs
MAPKKKs
MAPKKs
MAPKs
CNGCs
OSCAs
RBOHs
EP proteins
RPW8-NLRs

**Supplementary figure 27. Expression of PTI signalling components during PTI in *Arabidopsis thaliana*. (a-g)** The expression of PTI signalling components during PTI in *Arabidopsis thaliana*. *Arabidopsis thaliana* seedlings were treated with flg22, elf18, pep1, nlp20, OGs, chitin, or LPS to activate PTI. Light blue represents increased expression and light pink represents decreased expression during PTI. X-axis values represent log2 (fold change during PTI relative to samples at 0 min after treatment). RNA-seq data analyzed here were reported previously, where PTI was activated by different PAMPs/DAMPs in *A. thaliana* for 90mins. RNA-seq data were obtained from Bjornson et al, Nature Plants 2021 (reference 15 in main text).

Supplementary Figure 28

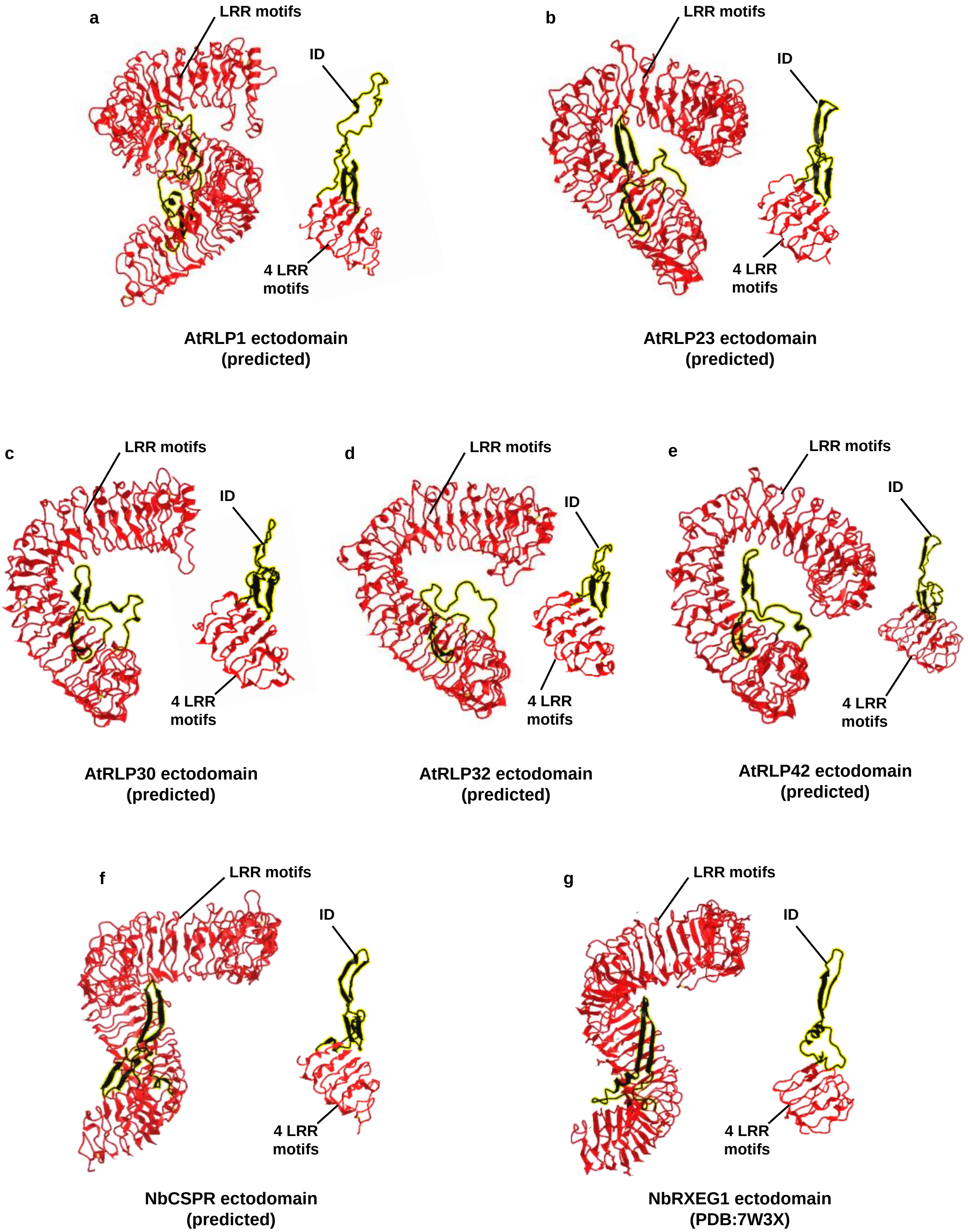

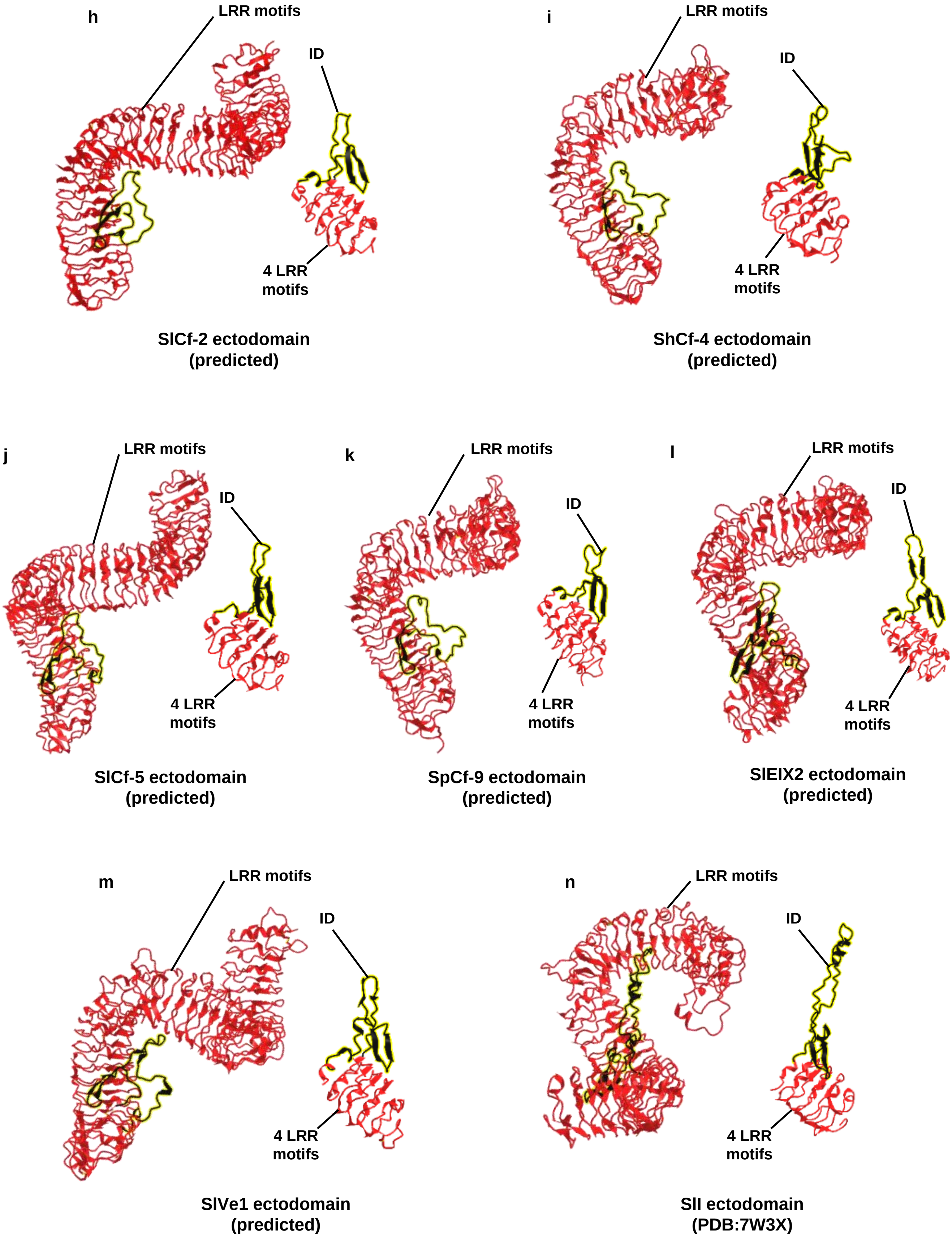
h
LRR motifs
ID
4 LRR motifs
SlCf-2 ectodomain (predicted)
i
LRR motifs
ID
4 LRR motifs
ShCf-4 ectodomain (predicted)
j
LRR motifs
ID
4 LRR motifs
SlCf-5 ectodomain (predicted)
k
LRR motifs
ID
4 LRR motifs
SpCf-9 ectodomain (predicted)
l
LRR motifs
ID
4 LRR motifs
SlEIX2 ectodomain (predicted)
m
LRR motifs
ID
4 LRR motifs
SlVe1 ectodomain (predicted)
n
LRR motifs
ID
4 LRR motifs
SlI ectodomain (PDB:7W3X)

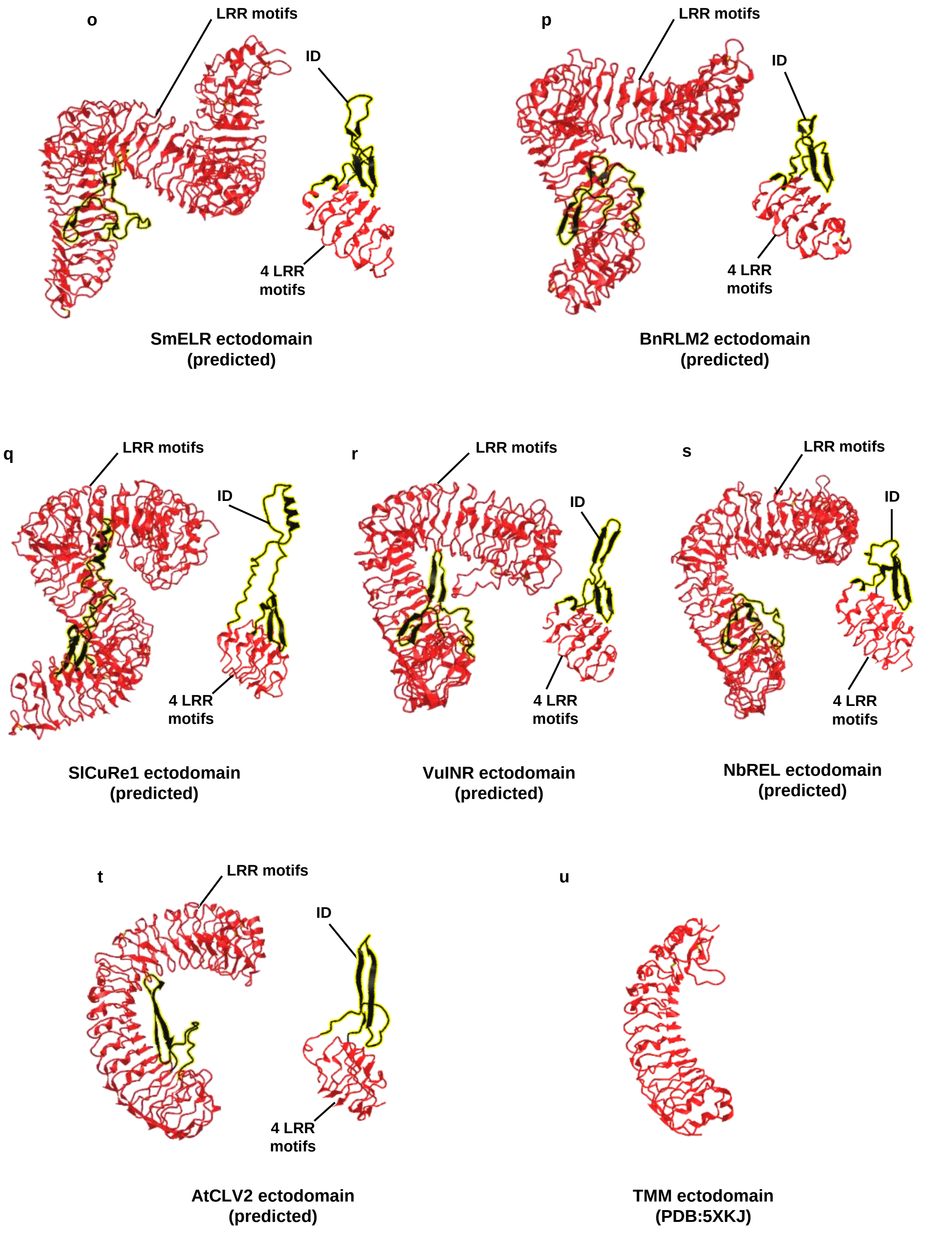


**Supplementary figure 28. Published or predicted structures of ectodomains of LRR-RLP in plants.** Ectodomain structures of characterized LRR-RLPs in plants. Structure of RXEG1 and TMM were obtained from PDB[112.113]. Structures of the other LRR-RLPs were predicted by Alphafold2[*,114.] Ectodomains were trimmed by PDBeditor and visualized in iCn3D[115]. Island domains (ID) are highlighted in black with yellow background.

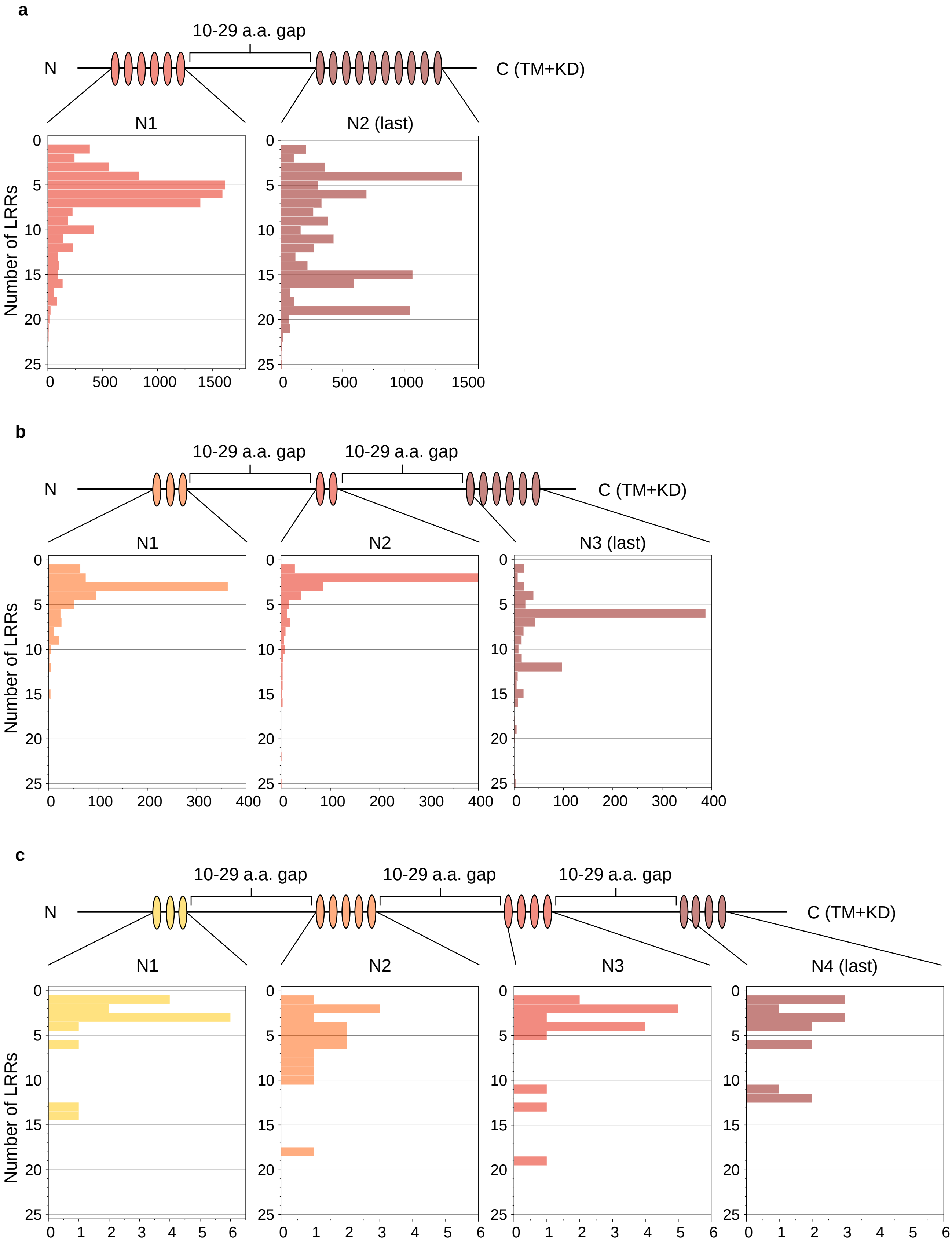


**Supplementary figure 29. Distribution of small gaps within LRR motifs in LRR-RLKs. (a-c)** Position of small gaps (NLs; 10-29 amino acids) in LRR-RLKs with **(a)** one, **(b)** two and **(c)** three small gaps. N1, N2, N3, and N4 represents the number of LRR motifs between the small gaps.

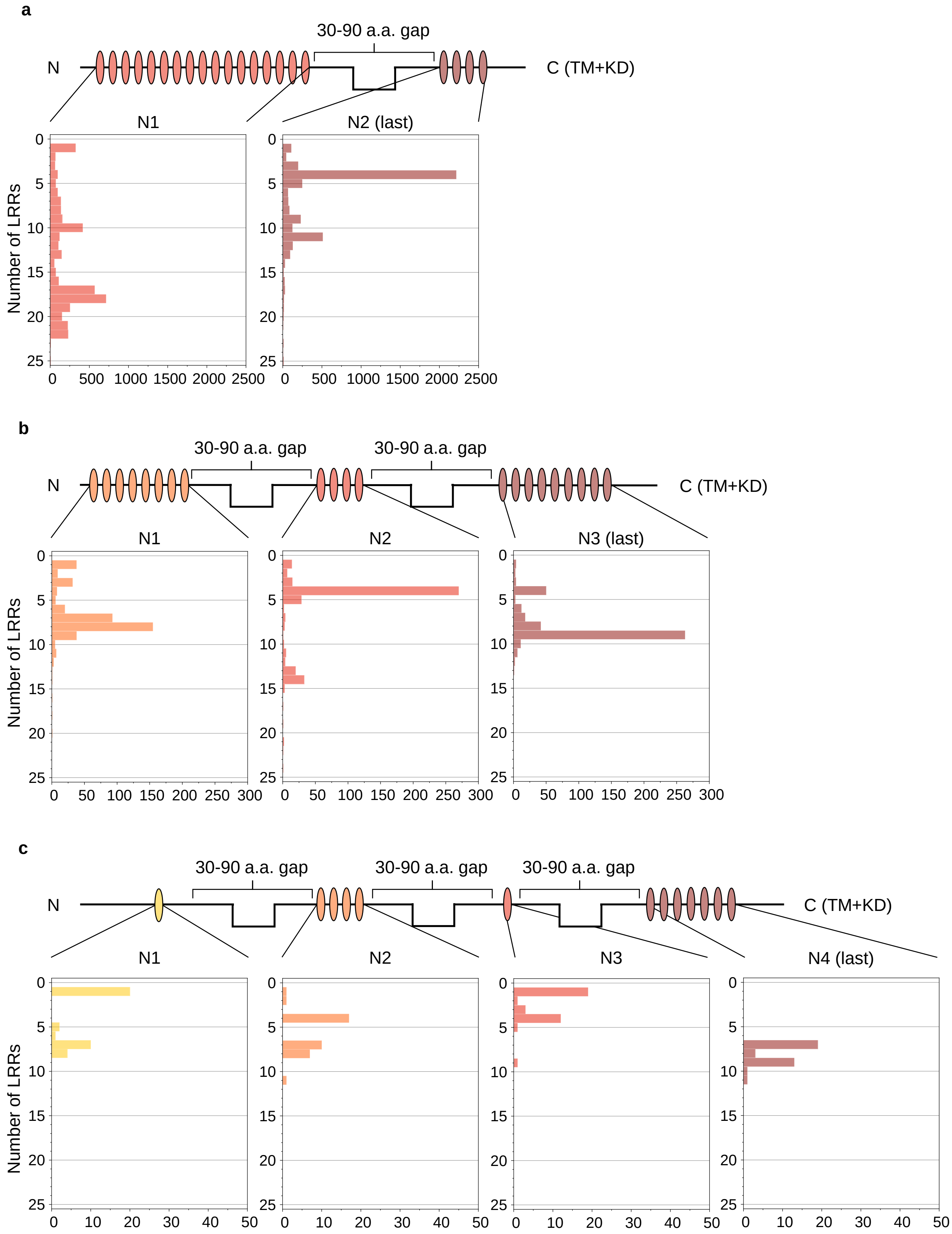


**Supplementary figure 30. Distribution of large gaps within LRR motifs in LRR-RLKs. (a-c)** Position of large gaps (IDs; 30-90 amino acids) in LRR-RLKs with **(a)** one, **(b)** two and **(c)** three large gaps. N1, N2, N3, and N4 represents the number of LRR motifs between the large gaps.

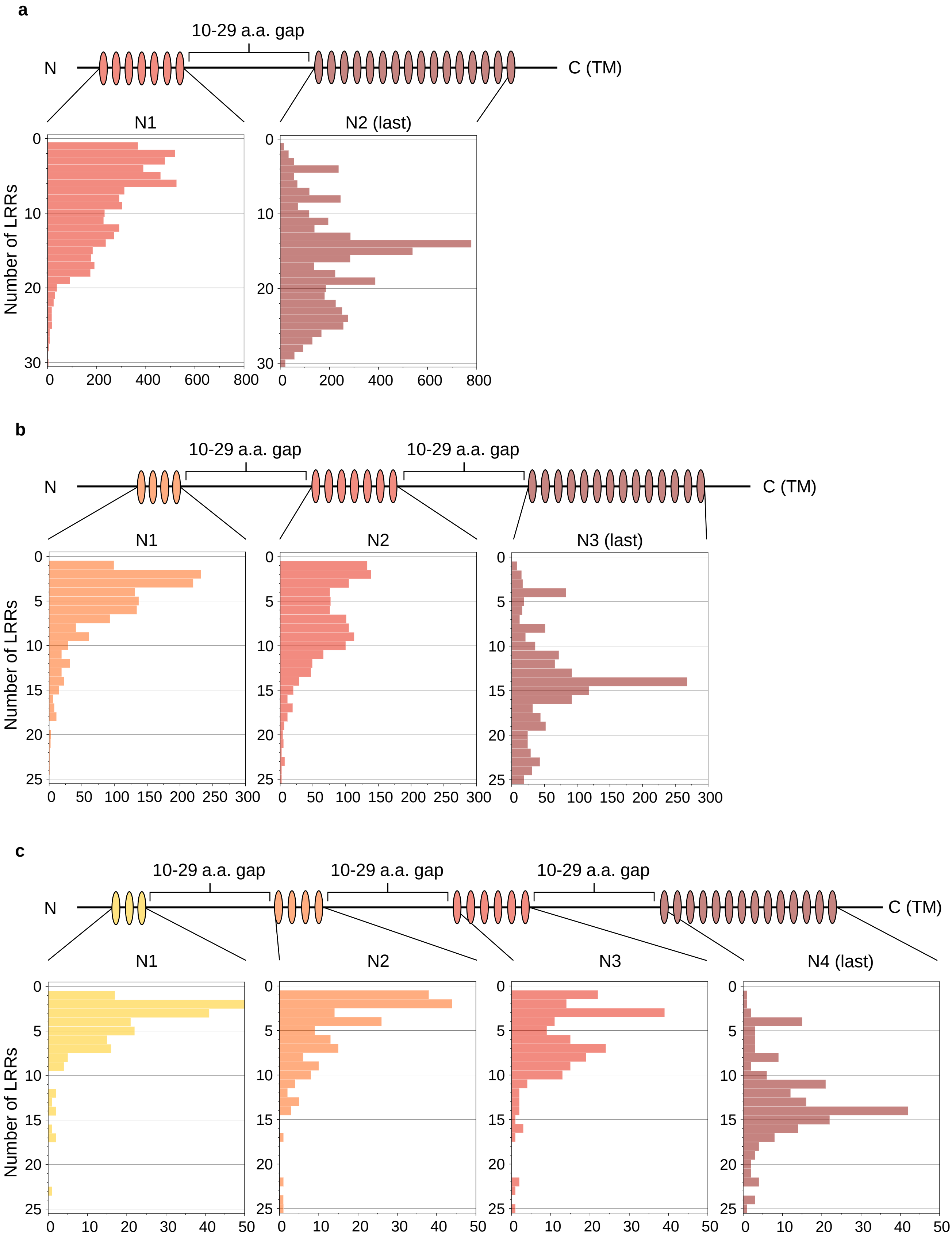


**Supplementary figure 31. Distribution of small gaps within LRR motifs in LRR-RLPs. (a-c)** Position of small gaps (NLs; 10-29 amino acids) in LRR-RLPs with **(a)** one, **(b)** two and **(c)** three small gaps. N1, N2, N3, and N4 represents the number of LRR motifs between the small gaps.

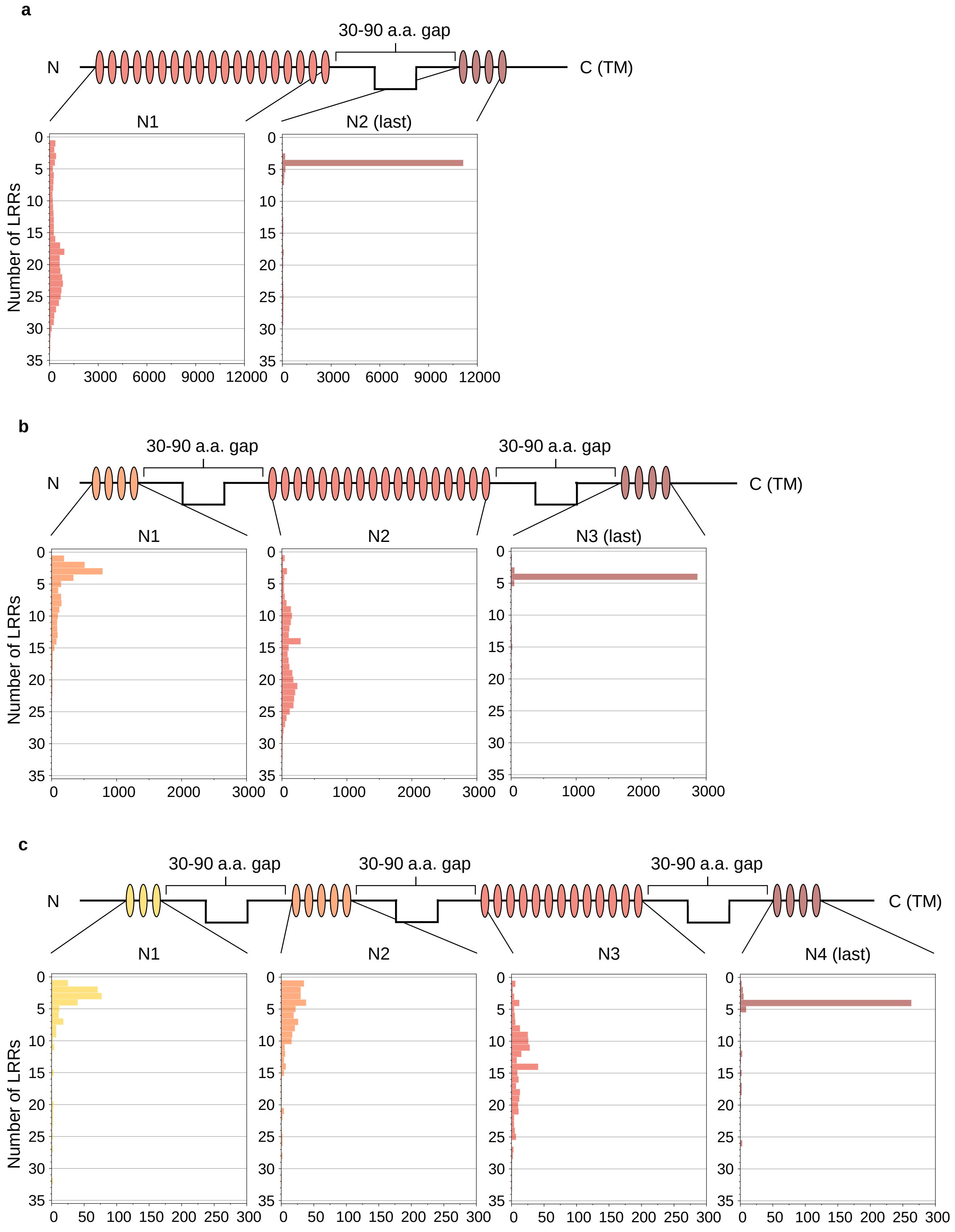


**Supplementary figure 32. Distribution of large gaps within LRR motifs in LRR-RLPs. (a-c)** Position of large gaps (IDs; 30-90 amino acids) in LRR-RLPs with **(a)** one, **(b)** two and **(c)** three large gaps. N1, N2, N3, and N4 represents the number of LRR motifs between the large gaps.

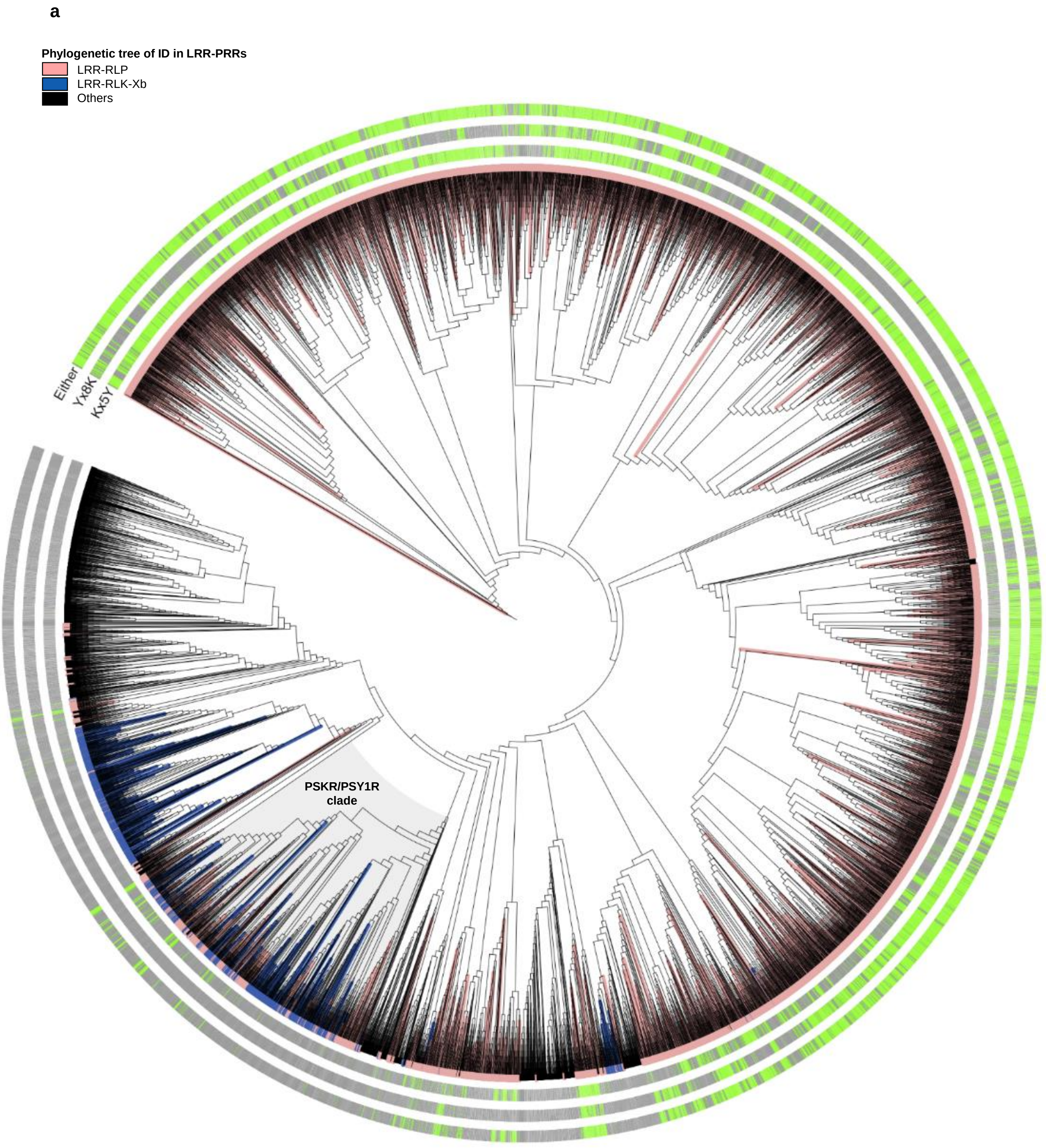


**Supplementary figure 33a. Phylogenetic tree of IDs of all LRR-containing PRRs from 350 species.** Branches are labelled in colours as indicated. Grey clade represents the PSKR/PSY1R clade. Presence of $Kx_5Y$, $Yx_8K$, or either Kx5Y or Yx8K in the IDs are indicated in green, and absence of these motifs are indicated in grey.

**b**

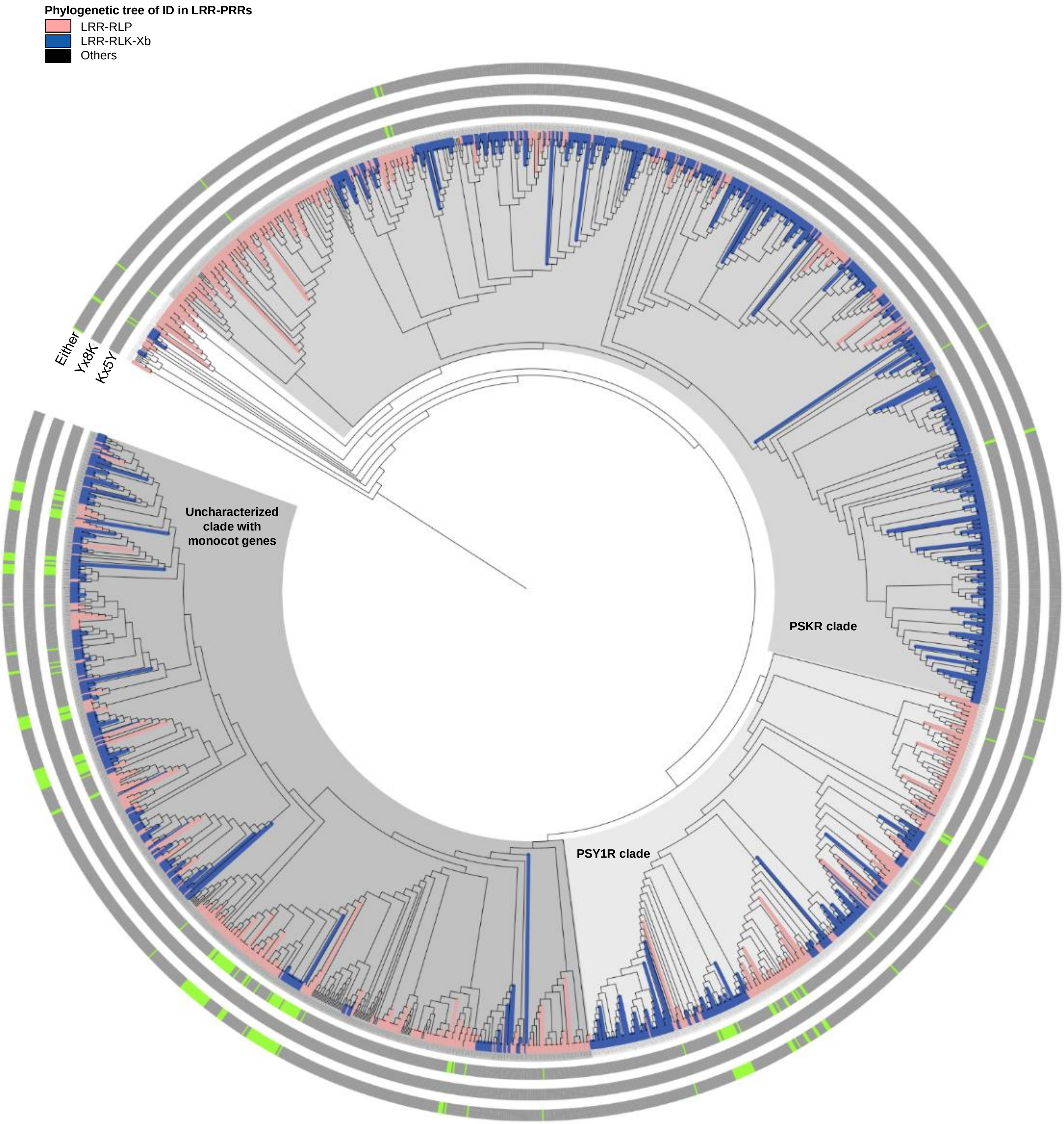


**Supplementary figure 33b. Phylogenetic tree of IDs from LRR-PRRs from the PSKR/PSY1R-clade.** Phylogenetic tree was extracted from the grey clade in supplementary figure 33a. Branches are labelled in colours as indicated. Presence of $Kx_5Y$, $Yx_8K$, or either Kx5Y or Yx8K in the IDs is indicated in green, and absence of these motifs is indicated in grey.

a

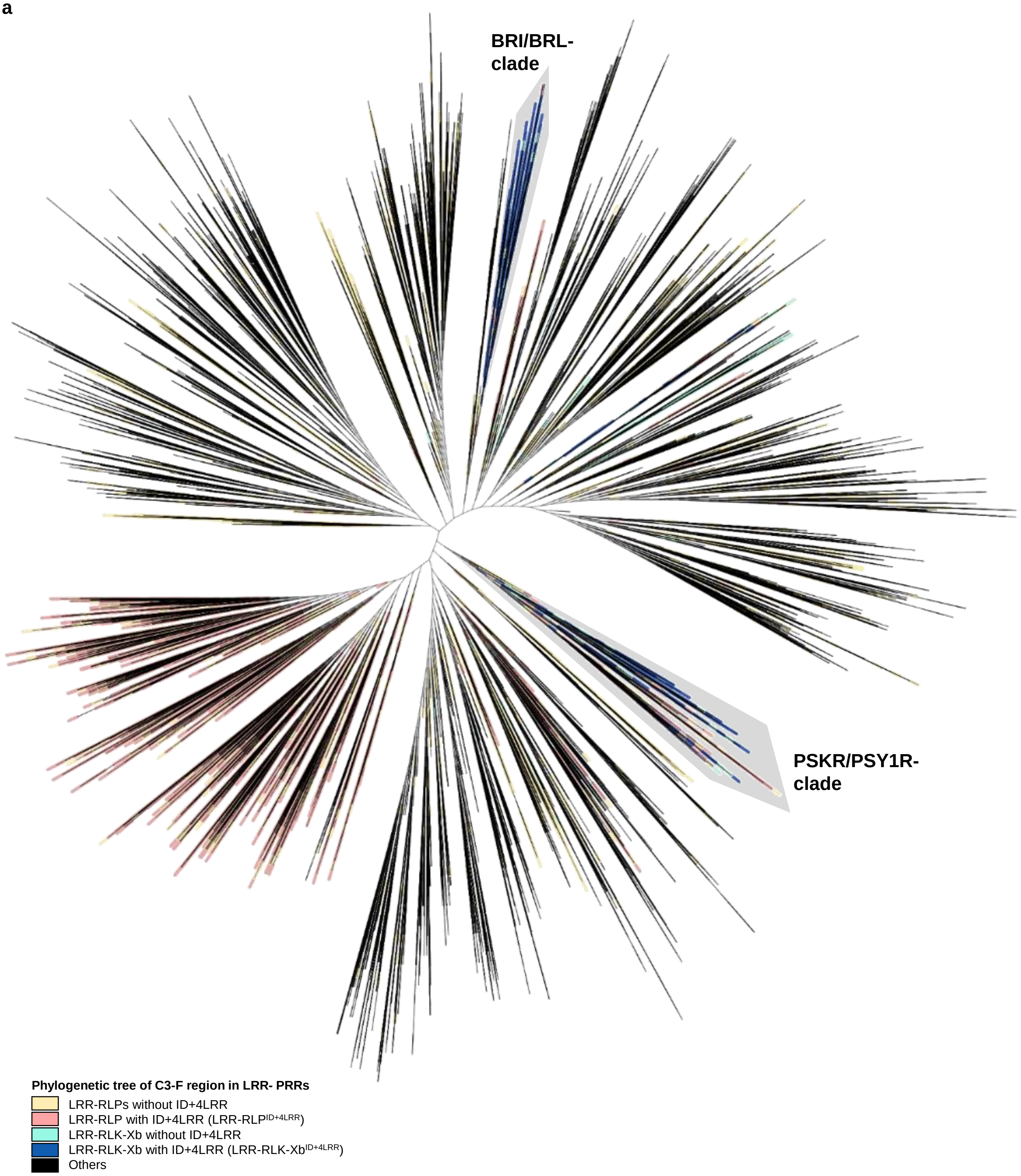


**Supplementary figure 34a. Phylogenetic tree of the ectodomain region of all LRR-containing PRRs from 350 species.** Clades and branches are labelled as indicated.

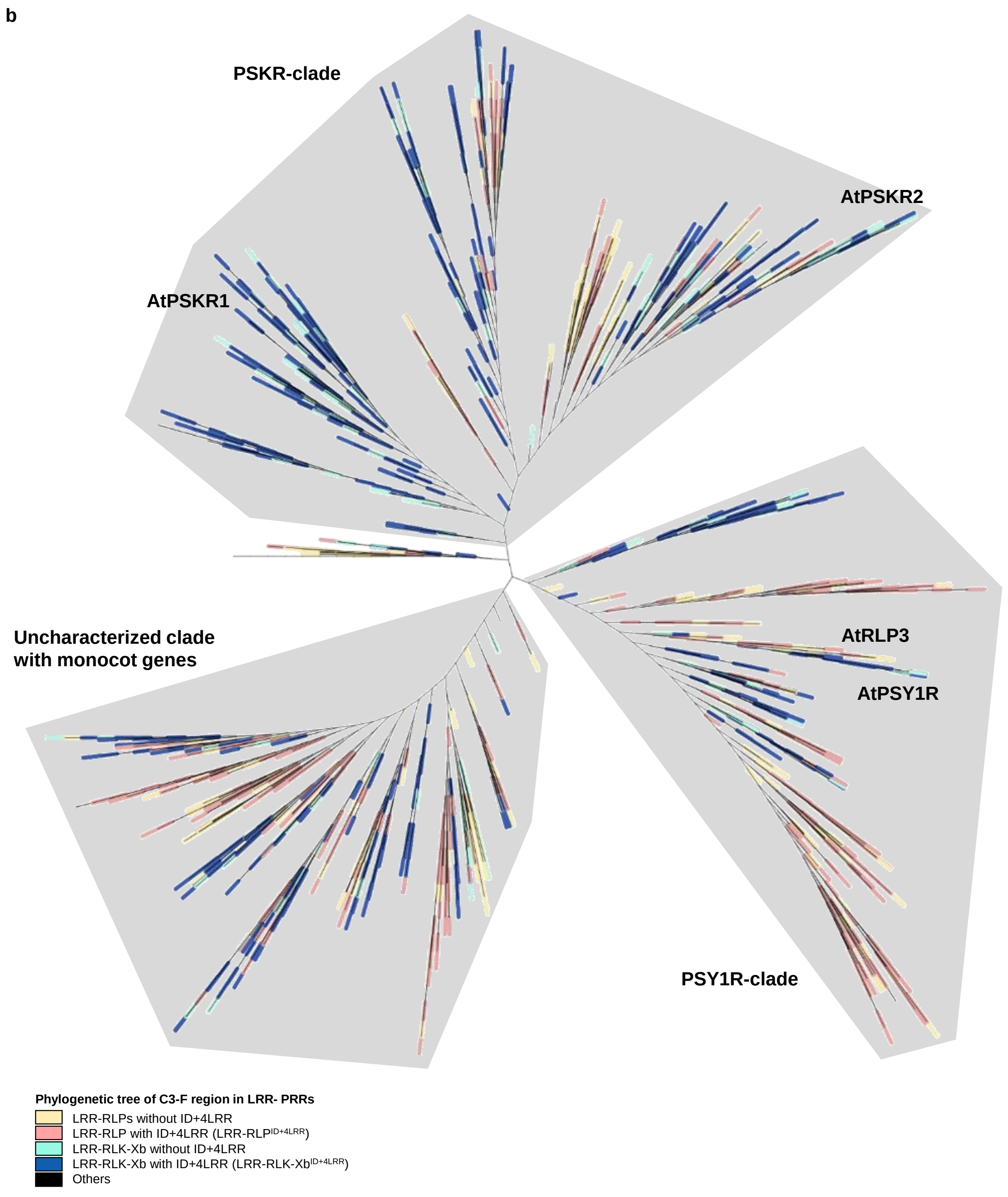


**Supplementary figure 34b. Phylogenetic tree of the ectodomain region of all LRR-containing PRRs from 350 species within the PSKR/PSY1R-clade.** Clades and branches are labelled as indicated.

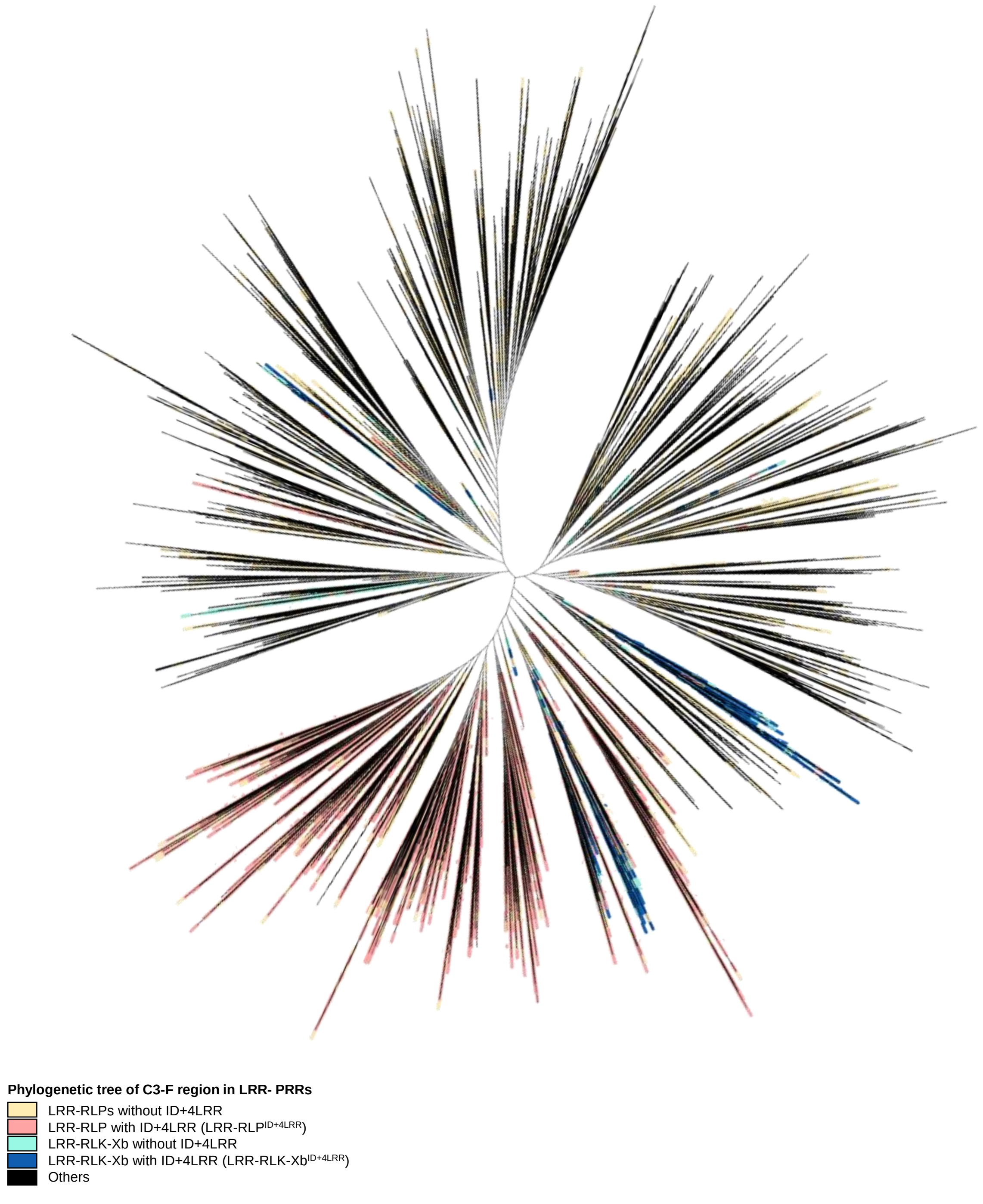


**Supplementary figure 35. Phylogenetic tree of the C3-F region of all LRR-containing PRRs from 350 species.** Clades and branches are labelled as indicated.

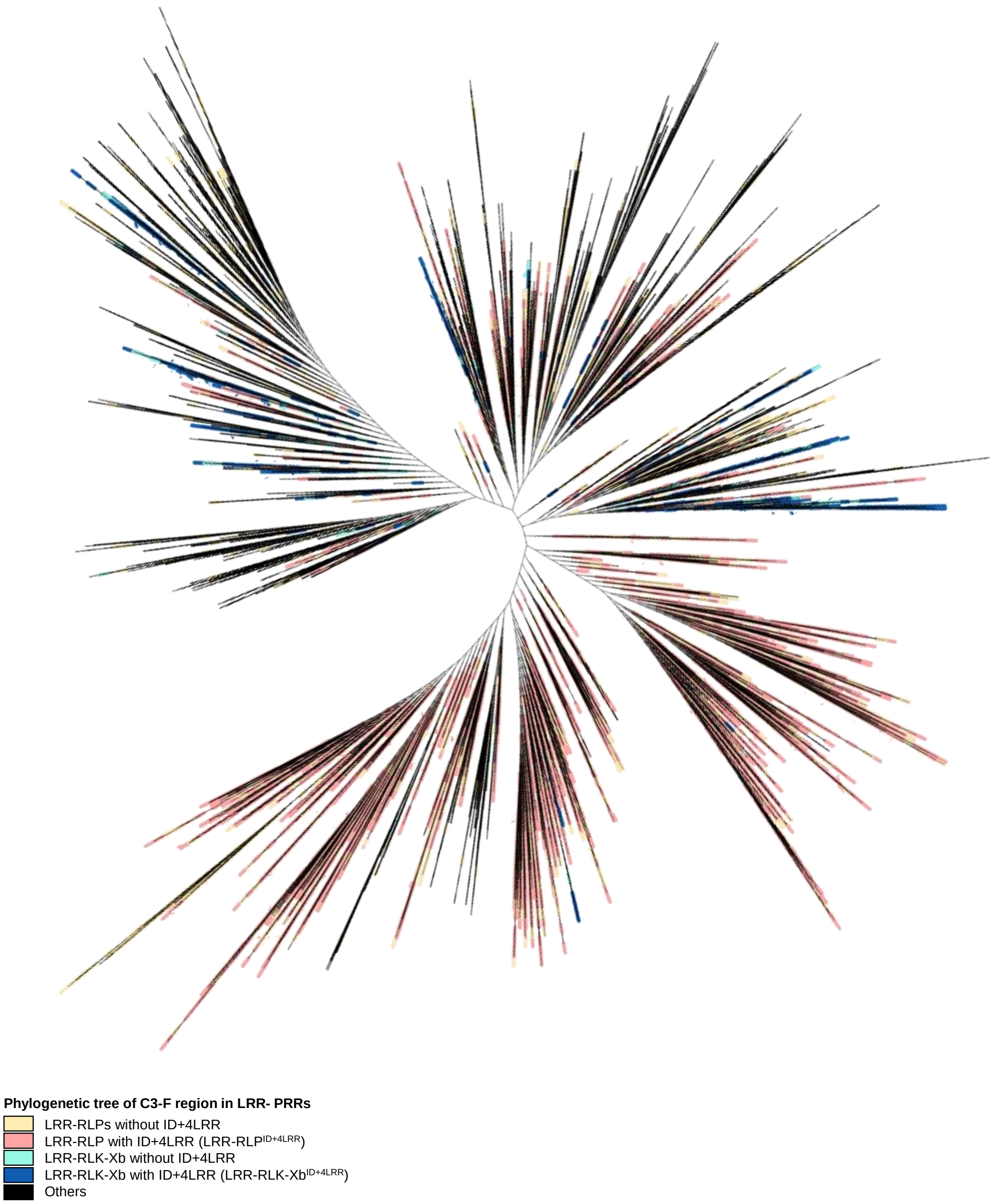


**Supplementary figure 36. Phylogenetic tree of the eJM region of all LRR-containing PRRs from 350 species.** Clades and branches are labelled as indicated.

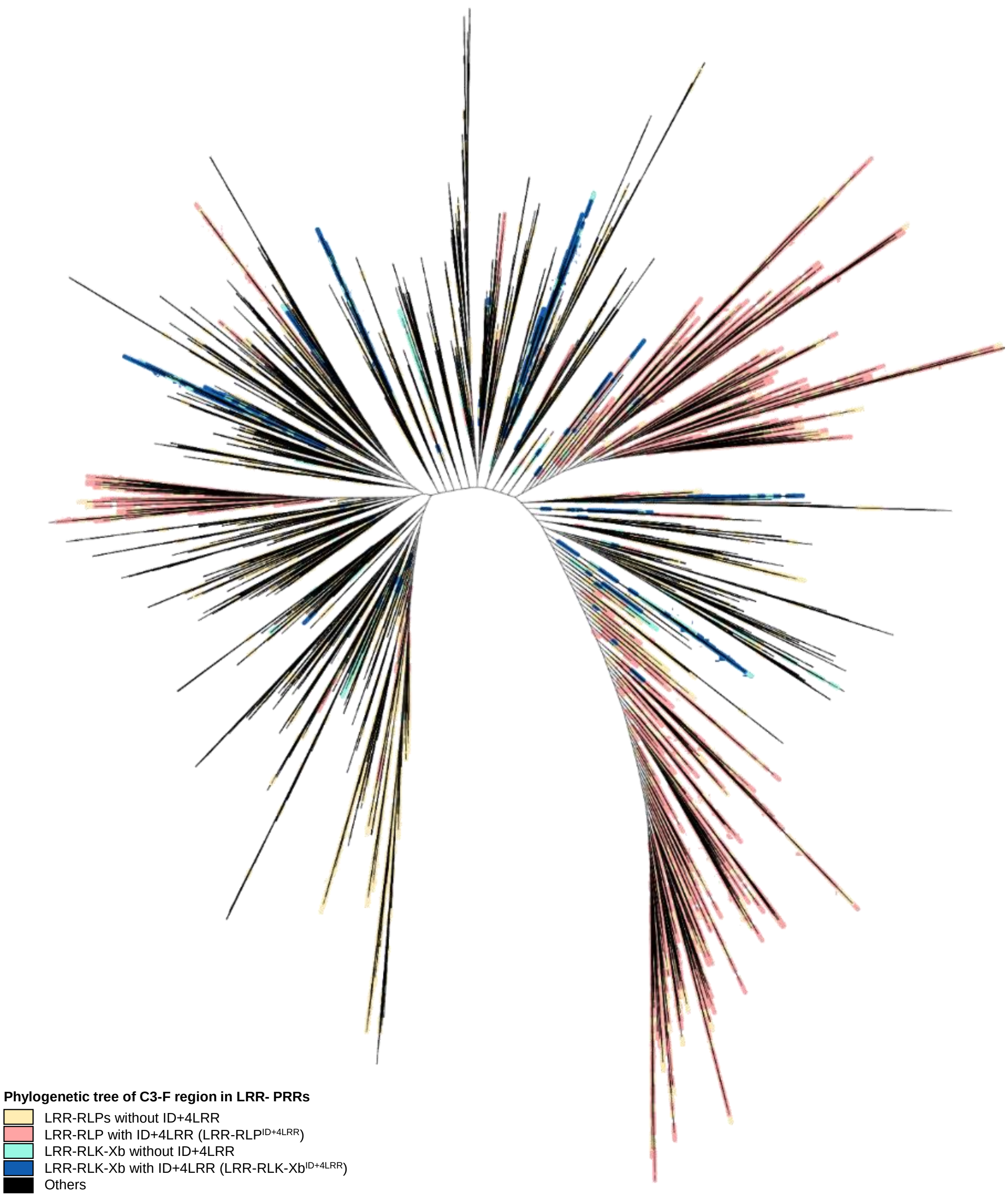


**Supplementary figure 37. Phylogenetic tree of the TM region of all LRR-containing PRRs from 350 species.** Clades and branches are labelled as indicated.
